# Frequency-dependent communication of information in networks of non-oscillatory neurons in response to oscillatory inputs

**DOI:** 10.1101/2025.07.09.664007

**Authors:** Andrea Bel, Horacio G. Rotstein

**Author notes:** Corresponding Researcher, CONICET, Argentina;. Graduate Faculty, Graduate Program in Neuroscience (GPN), Center for Molecular and Behavioral Neuroscience (CMBN), Rutgers University.

## Abstract

Understanding how neuronal networks process oscillatory inputs is key for deciphering the brain’s information processing dynamics. Neuronal filters describe the frequency-dependent relationship of neuronal outputs (e.g., membrane potential amplitude, firing rate) and their inputs for the level of neuronal organization (e.g., cellular, network) considered. Band-pass filters are associated to the notion of resonance and reflect the system’s ability to respond maximally to inputs at a nonzero (resonant) frequency or a limited (resonant) frequency band. The complementary notion of phasonance refers to the ability of a system to exhibit a zero-phase response for a nonzero (phasonant) input frequency. The biophysical and dynamic mechanisms that shape neuronal filters and give raise to preferred frequency responses to oscillatory inputs are poorly understood beyond single cells. Moreover, the mechanisms that control the frequency-dependent communication of information across cells in a network remain unclear. Here, we use mathematical modeling, analytical calculations, computational simulations and dynamical systems tools to investigate how the complex and nonlinear interaction of the systems’s biophysical properties and interacting time scales shape neuronal filters in minimal network models receiving oscillatory inputs with frequencies (*f*) within some range. The minimal networks consist of one directly stimulated cell (cell 1) connected to another (not directly stimulated) cell (cell 2) via graded chemical synapses. Individual cells are either passive or resonators and chemical synapses are either excitatory or inhibitory. The network outputs consist of the voltage peak envelopes and the impedance amplitude and phase profiles (as a function of *f*) for the two cells. We introduce the frequency-dependent amplitude *K*(*f*) and phase ΔΦ(*f*) communication coefficients, defined as the ratio of the amplitude responses of the indirectly and directly stimulated cells and the phase difference between these two cells, respectively. Extending previous work, we also introduce the *K*-curve, parametrized by *f*, in the phase-space diagram for the voltage variables of the two participating cells. This curve joins the peak voltage values of the two cells in response to the oscillatory inputs and is a geometric representation of the communication coefficient. It allows to interpret the results and explain the dependence of the properties of the communication coefficient in terms of the biophysical and dynamic properties of the participating cells and synaptic connectivity when analytical calculations are not possible. We describe the conditions under which one or the two cells in the network exhibit resonance and phasonance and the conditions under which the network exhibits *K*-resonance and ΔΦ-phasonance and more complex network responses depending as the complexity of the participating cells increases. For linear networks (linear nodes and linear connectivity), *K* is proportional to the impedance of the indirectly activated cell 2 and ΔΦ is equal to the phase of the indirectly stimulated cell 2, independent of the directly stimulated cell 1 in both cases. We show that the presence of nonlinear connectivity in the network creates (nonlinear) interactions between the two cells that give rise to *K*-resonance, ΔΦ-phasonance and more complex responses that are absent in the corresponding linear networks. The results and methods developed in this paper have implications for the processing of information in more complex networks.

## 1 Introduction

Understanding how neuronal networks process information is crucial to comprehending the complex workings of the brain. Neurons and neuronal circuits are subject to inputs that often have oscillatory components within certain frequency range. Neuronal filters describe the frequency-dependent relationship of neuronal outputs (e.g., membrane potential amplitude, firing rate) on their inputs (e.g., sinusoidal functions, spike trains, white noise). Neuronal filters shape neuronal communication [1–6] and play a major role in information processing, rhythm generation and computations within the brain [3,4,7–18] by selectively amplifying and attenuating specific signal components. In systematic and mechanistic studies, neuronal filters are obtained by computing the system’s response (e.g., membrane potential amplitude, firing rate) to periodic inputs (with frequency *f*), and therefore the two notions (filters and response) are typically used indistinctly.

Band-pass filters are closely tied to the notion of resonance defined as the ability of a system to respond maximally to input frequencies at a nonzero (resonant) frequency or limited frequency band (e.g., Figs. 2-A2 and -B2 at *f* = *f*_*res*_) [1, 4, 19, 20]. This is in contrast to low-pass filters that are monotonically decreasing functions of the input frequency (e.g., Figs. 2-A1 and -B1). A more complete description of the input-output relationships requires the use of frequency-dependent phase profiles characterizing de frequency ranges for which there is a phase lag (e.g., Fig. 2-C1 and Fig. 2-C2 for *f > f*_*phas*_) and phase lead (e.g., Fig. 2-C2 for *f < f*_*phas*_). Together, these two frequency-dependent quantities constitute the system’s transfer function, which can be used to reconstruct the system’s dynamics in response to arbitrary inputs with small enough amplitude (for linear systems, there is no restriction on the input amplitude). For nonlinear systems under certain conditions the transfer function is informative enough of the input-output relationship even if the amplitude restriction is not met.

Neuronal resonance has been investigated both experimentally and theoretically in a variety of brain areas at the single cell and, to a lesser extent, the synaptic and network levels of organization [3, 4, 12, 18–36]. In networks, resonance can be inherited from lower levels of organization (e.g., cellular) or generated independently by the interplay of the network building blocks [4, 35]. Some of these studies have extended their scope to neuronal phasonance, defined as the ability a system to exhibit a zero-phase response for a nonzero (phasonant) frequency (e.g., Fig. 2-C2 for *f* = *f*_*phas*_) [25–27, 35, 37] and more complex frequency-dependent profiles (e.g., antiresonance and antiphasonance) [19, 26, 27, 37].

Neuronal filters are shaped by the complex and nonlinear interaction of the systems’s biophysical components and involve the interplay of a multiplicity of time scales (e.g., ionic currents, synaptic currents) [1–4, 19, 21, 22, 26, 27, 31, 32, 37–40]. However, the biophysical and dynamic mechanisms of generation of neuronal resonance and phasonance beyond single cells remain to be understood. It is unclear how the response of a directly stimulated cell is transmitted to other (not directly stimulated) cells in the network, and how this frequency-dependent communication is controlled by the biophysical properties of the participating nodes and synapses. It is also unclear whether, and if yes how and under what conditions, there are preferred frequencies (or frequency bands) at which this communication is maximal.

In this paper, we address these issues by using minimal two-cell network models (Fig. 1) consisting of one directly stimulated (cell 1), one not directly stimulated cell (cell 2) and instantaneous graded synapses [41–49] (slaved to voltage). The cells can be either passive (1D; red nodes in Fig. 1) or resonators (2D; blue nodes in Fig. 1). The latter describe the dynamics of linearized models having a resonant current (e.g., hyperpolarization-activated, or h-, M-type slow potassium, calcium inactivation), and an instantaneously fast amplifying current (e.g., persistent sodium, calcium activation), which are incorporated in the linearized leak current. Graded synapses are inherently nonlinear and active at subthreshold levels. In previous work [50], we used this model in the absence of external stimulation to describe the resonant-based mechanism of generation of oscillations in networks of non-oscillatory neurons. Here we focus on a parameter regime where network oscillations are not present in the absence of external inputs. The extension of our study to cellular models having more complex dynamics such as resonators with subthreshold nonlinearities of quadratic type in the phase-space [27] is straightforward.

**Figure 1:**
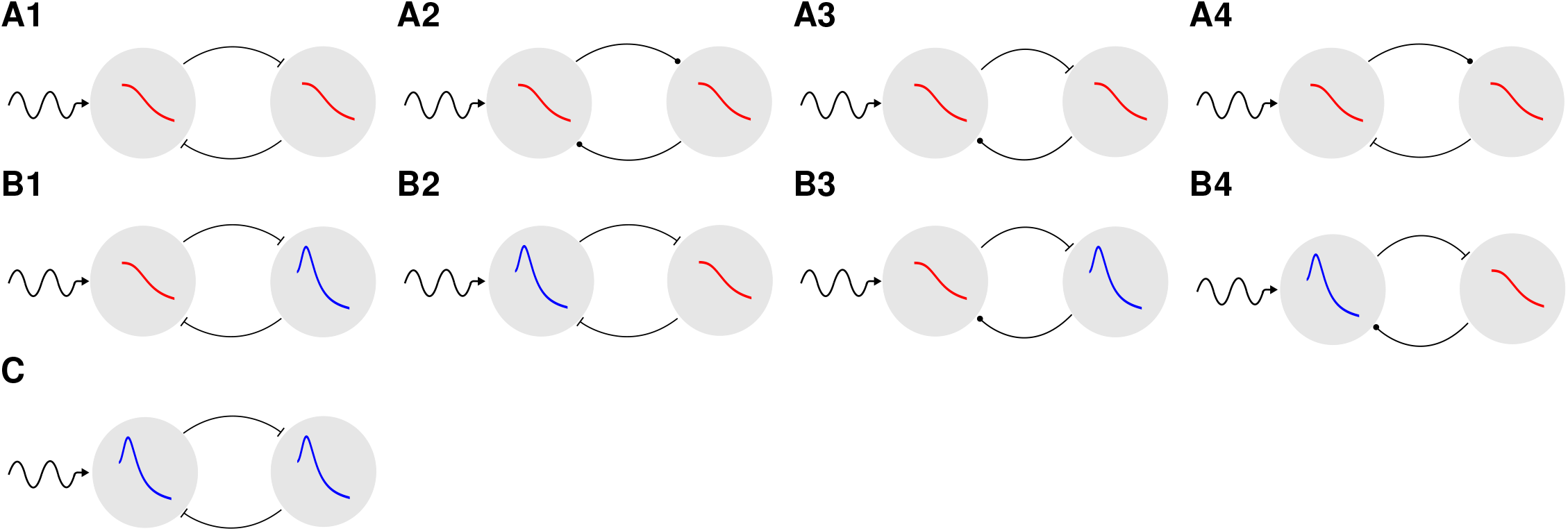
Network diagrams. A circle means that the connection is excitatory and a bar means that it is inhibitory. The external input arrives to cell 1 (left cell). **First row (A)**. Passive cells (1D/1D). **A1**. Mutual inhibition. **A2**. Mutual excitation. **A3**. Inhibition-excitation. **A4**. Excitation-inhibition. **Second row (B)**. Passive and resonator cells (1D/2D). **B1**. Mutual inhibition. Cell 1 passive. **B2**. Mutual inhibition. Cell 1 resonator. **B3**. Inhibition-excitation. Cell 1 passive. **B4**. Inhibition-excitation. Cell 1 resonator. **Third row (C)**. Resonator cells (2D/2D). Mutual inhibition.

In a network, the response profiles of the indirectly and directly stimulated cells are different (and both are different from the response profiles of isolated cells when directly stimulated). The quotient *K*(*f*) of their amplitudes is a measure of the effective communication of information between the cells or their relative synaptic strength. This extends the notion of the coupling coefficient for gap junctions in response to constant (DC) inputs [51–65] and oscillatory inputs [65, 66], which has been used to measure the strength of the connection or the efficacy of the signal transmission. The difference ΔΦ(*f*) of the response phase profiles, referred to as the phase communication coefficient, provides information about the synchrony properties (if they exist) of the two cells. Together, *K*(*f*) and ΔΦ(*f*) are useful tools to characterize the frequency-dependent properties of the effective communication of information between the two cells in response to external, time-dependent inputs.

We investigate the frequency-dependent properties of the *K*(*f*) and ΔΦ(*f*) profiles (curves of these quantities as a function of the input frequency *f*) as a function of the cellular and synaptic biophysical and dynamic properties, identify the conditions under which the network exhibits *K*-resonance and ΔΦ-phasonance, defined as the ability of the network to exhibit a peak in the *K*(*f*) profile and a zero ΔΦ response at a non-zero input frequency, and determine the relationship (or lack of thereof) between these phenomena and the occurrence of (network) resonance and phasonance in the participating cells in the network.

For linear networks, both *K*(*f*) and ΔΦ(*f*) can be computed analytically, which is not possible for non-linear networks. To investigate the dynamic mechanisms that give rise to the network response profiles and the preferred frequency responses (resonances) they may exhibit, we extend the methods introduced in [27, 37] to develop dynamical system tools for the qualitative, phase-space analysis of the dependence of the frequency-dependent response profiles on the model parameters, and the relationship between the responses of the individual cells in the network and the communication coefficient. We introduce the notion of the *K*-curve associated to the *K* profile. The *K*-curve is a tool to visualize the simultaneous and relative changes of the *v*_1_ and *v*_2_ peaks in the “moving” phase-plane diagram (the projection of the phase-space diagram for the participating variables and the oscillatory input) as a function of the input frequency *f*. We derive a geometric condition between the position and tangent vectors on the *K*-curve for the occurrence of resonance. These geometric tools allow us to better understand how *K*(*f*) is shaped by the network building blocks by looking at the dynamic interaction between the vector field of the autonomous (unforced) system and the oscillatory input that cyclically modulates this vector field over a range of input frequencies. We extend this analysis to higher-dimensional nodes, which requires the use of projections.

The dynamic structure of the models we use in this paper is qualitatively similar to these for firing rate models of Wilson-Cowan type [67–69] and firing rate models with adaptation [47–49, 70]. Therefore, our results and the tools developed in this paper have implications for the investigation of the response of neuronal populations to external inputs [35].

## 2 Methods

We consider a two-cell network with graded synaptic connections receiving oscillatory inputs. The dynamics of the individual cells are described by linearized biophysical (conductance-based) models [19, 26] and the graded synaptic connectivity is nonlinear [43, 71, 72]. We refer the reader to Supplementary Material S1 for details on the linearization process for conductance-based models (see also [19, 26]). The formalism and tools we develop and use in this paper can be extended to recurrently connected networks with an arbitrary number of nodes having both linear and nonlinear dynamics.

### 2.1 Networks of linearized cells with graded synapses

The network model is described by the following system of equations

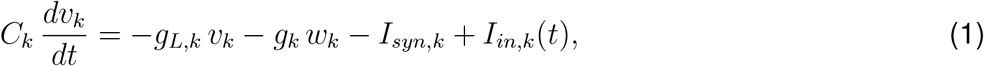

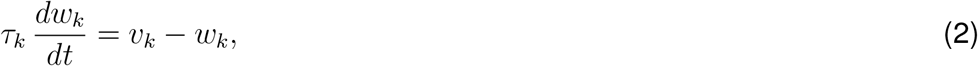

for *k* = 1, 2. In eqs. (1)-(2) *t* is time, *v*_*k*_ represent the membrane potential, *w*_*k*_ represent the gating variable, *C*_*k*_ is the capacitance, *g*_*L,k*_ is the linearized leak maximal conductance, *g*_*k*_ is the ionic current linearized conductance, *τ*_*k*_ is the linearized time constant controlling the evolution of *w*_*k*_, *I*_*in,k*_(*t*) is a time-dependent input current, and *I*_*syn,k*_ is the graded synaptic current. The linearized component of ionic currents with instantaneously fast dynamics are incorporated in *g*_*L,k*_. Examples include the persistent sodium current (*I*_*Nap*_) and T-type calcium (*I*_*Ca,T*_) activation gating variable. Examples of ionic currents with relatively slow dynamics include the hyperpolarization-activated mixed-cation *I*_*h*_ current, the M-type slow-potassium current *I*_*Ks*_ and the T-type calcium inactivation *I*_*CaT*_ process. The variables *v*_*k*_ are relative to the voltage coordinate of the equilibrium (resting) potential, 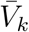 of the original conductance-based model, and the variables *w*_*k*_ are relative to the gating variable coordinate of the fixed-point 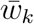 of the original conductance-based model, normalized by the derivative of the corresponding activation curve at 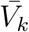 (see Supplementary Material S1).

The synaptic current (from cell *j* to cell *k*) is given by

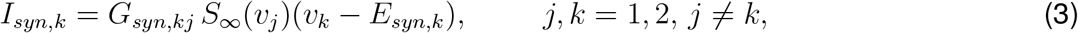

where *G*_*syn,kj*_ is the maximal synaptic conductance, *E*_*syn,k*_ is the synaptic reversal potential (relative to 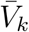) and

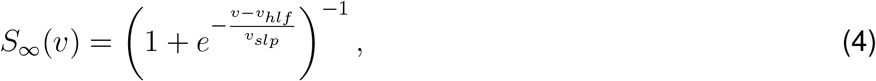

where the half-activation point *v*_*hlf*_ is also relative to 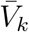.

We use the following units: mV for voltage, ms for time, *µ*F/cm^2^ for capacitance, *µ*A/cm^2^ for current, mS/cm^2^ for the maximal conductances and Hz for frequency.

Note that the heterogeneity due to different values of the biophysical parameters in the original conductance-based model is translated into the linearized model both directly and indirectly through the equilibria (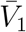 and 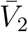). Unless stated otherwise, we used the following parameter values: *C*_1_ = *C*_2_ = 1, *v*_*hlf*_ = 0, *v*_*slp*_ = 1, *E*_*in*_ = *−*20, *E*_*ex*_ = 60.

### 2.2 Single cell voltage response to oscillatory inputs: low- and band-pass filters

The voltage response of a neuron receiving an oscillatory input current (with frequency *f*) is typically measured in terms of the so-called impedance function **Z**(*f*), a complex quantity with amplitude *Z*(*f*) = | **Z**(*f*) | and phase Φ(*f*). For simplicity, we refer to *Z*(*f*) simply as the impedance unless the full notation is necessary to avoid confusion or ambiguity. We refer to the graphs of these quantities as the impedance (*Z*-) and phase (Φ-) profiles. Note that *Z*(*f*) and Φ(*f*) are steady state quantities.

A system exhibits *resonance* if *Z*(*f*) peaks at a non-zero (resonant) frequency *f*_*res*_ (e.g., Figs. 2-A2) and *phasonance* if Φ(*f*) vanishes at a non-zero (phasonant) frequency *f*_*phas*_ (e.g., Figs. 2-C2). Following circuit theory, we refer to a system exhibiting resonance as a *band-pass filter* (Figs. 2-A2) and to a system exhibiting a monotonically decaying *Z*-profile as a *low-pass filter* (Fig. 2-A1). The generation of cellular resonance requires the interplay of negative and positive feedback effects, which are mediated by the ionic current gating variables or related processes. Resonant ionic processes (e.g., *I*_*h*_, *I*_*Ks*_ and *I*_*CaT*_ activation) oppose changes in voltage, while amplifying ionic processes (e.g., persistent sodium *I*_*Nap*_ and *I*_*Ca,T*_ activation) favor these changes.

**Figure 2:**
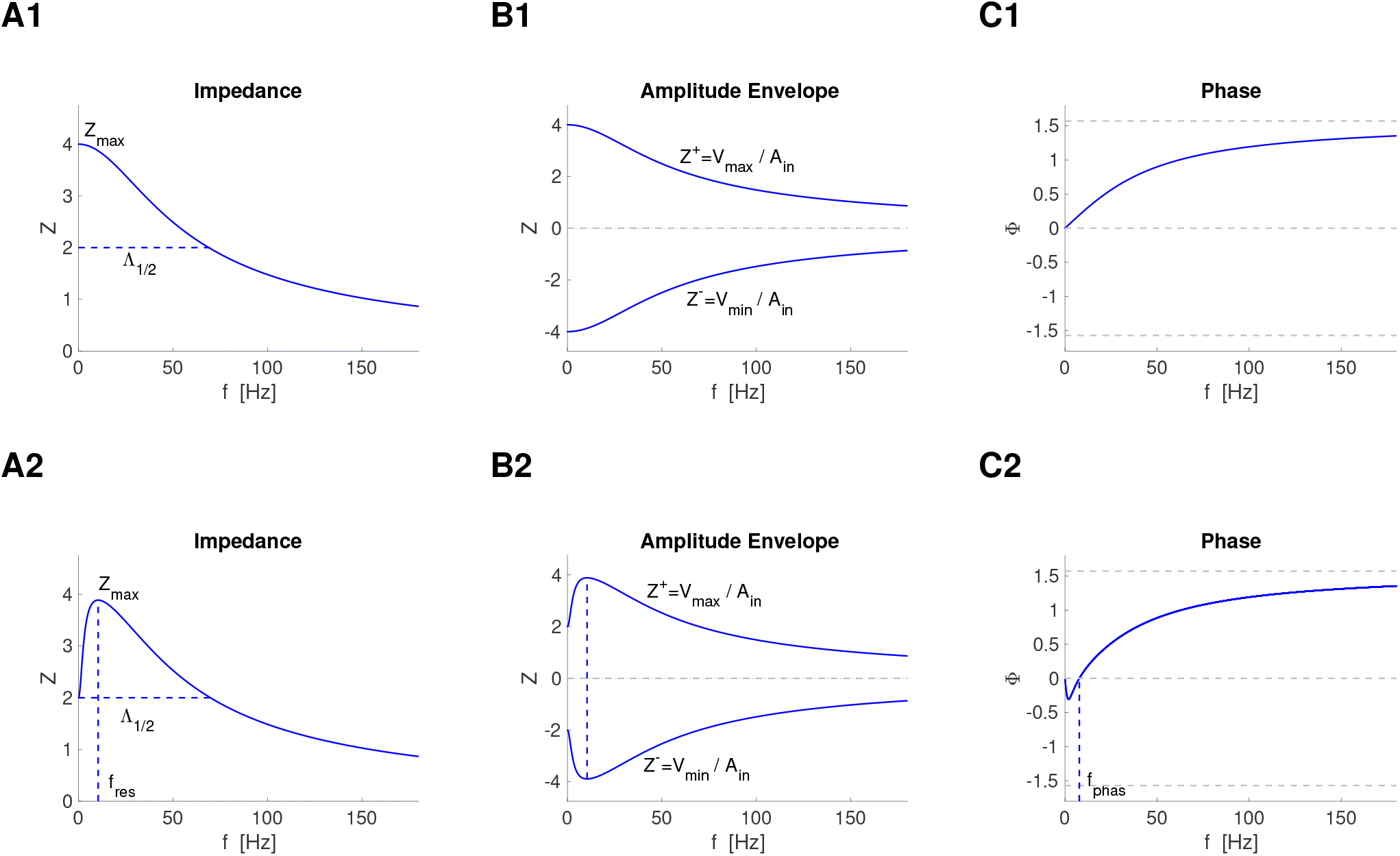
Schematic diagrams of the impedance (A), amplitude envelope (B) and phase (C) profiles for a low-pass filter cell (top) and a resonant cell / band-pass filter (bottom). Curves of the impedance (*Z*), amplitude envelope (*V*_*max*_ and *V*_*min*_) and phase (Φ) as a function of the input frequency *f*. **A**. Λ_1*/*2_ is the frequency range for which *Z* is above *Z*_*max*_*/*2. **A1**. *Z*_*max*_ = *Z*(0). **A2**. The resonant frequency *f*_*res*_ is the non-zero input frequency at which *Z*(*f*) peaks. **B1 - B2** *V*_*max*_/*V*_*min*_ are the maximum/minimum steady state voltage response values to the oscillatory inputs. **C1**. The voltage response is delayed for all input frequencies *f*. **C2**. The phasonant frequency *f*_*phas*_ is the non-zero input frequency at which Φ = 0. The voltage response is advanced for *f < f*_*phas*_ and delayed for *f > f*_*phas*_.

The resonant properties of 2D linear neuronal systems, including their relationship between the cellular intrinsic biophysical properties and the dynamic mechanism of generation of resonance have been investigated extensively by us and other authors [19, 26, 37].

For a linear system receiving a sinusoidal input current of the form

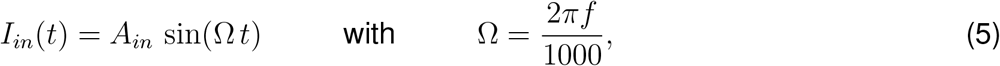

the voltage response is given by

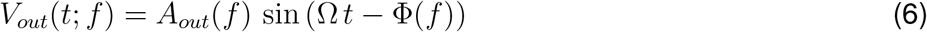

where *A*_*out*_(*f*) is the amplitude and Φ(*f*) is the phase-shift (or phase), which captures the difference between the peaks of *I*_*in*_(*t*) and *V*_*out*_(*t*; *f*) normalized by the period. The impedance is defined as

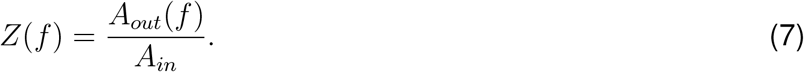

In the Supplementary Material S2 we present the expressions of *Z*(*f*) and Φ(*f*) for linear neuronal 2D and 1D models.

For nonlinear systems we extend the notion of impedance by defining the amplitude envelope impedance profile [37, 73] as

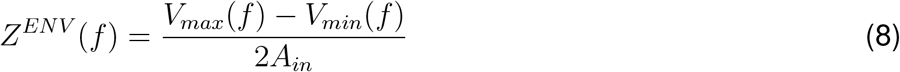

where *V*_*max*_(*f*) and *V*_*min*_(*f*) are the peak and trough envelope profiles, respectively, defined as the maximum and minimum values, respectively, of the steady-state oscillatory voltage *V*_*out*_(*f*) as a function of the input frequency *f*. We refer to the graphs of *V*_*max*_(*f*) and *V*_*min*_(*f*) divided by the input amplitude *A*_*in*_ as the upper (*Z*^+^) and lower (*Z*^*−*^) *Z*-envelopes, respectively (Fig. 2-B1 and -B2). For linear systems, *Z*^*ENV*^ (*f*) = *Z*(*f*). The resonant frequency *f*_*res*_ is then the peak frequency of *Z*^*ENV*^ in eq. (8). Similarly, the phase Φ is computed as the distance between the peaks of the output and closest input normalized by period, and *f*_*phas*_ is the non-zero frequency where Φ vanishes. These quantities extend the concept of impedance and phase to nonlinear systems under certain conditions (the input and output frequencies coincide and the output amplitude is uniform across cycles for a given input with constant amplitude).

An important aspect to note is that resonance can occur in the absence of intrinsic damped oscillations [19, 26]; i.e., when the fixed-point is a stable node. In this paper we focus on *resonators* defined as neurons, and in general dynamical systems, that exhibit resonance in the absence of intrinsic oscillations.

### 2.3 Network voltage response to oscillatory inputs

We now consider a two-cell network receiving sinusoidal inputs of the form (5) *I*_*in,k*_(*t*) = *A*_*in,k*_ sin(Ω *t*) (*k* = 1, 2). In most of this paper, we consider inputs to only one cell (cell 1); i.e., *I*_*in*,1_(*t*) = *A*_*in*_ sin(Ω *t*) and *I*_*in*,2_(*t*) = 0. However, for generality we develop the mathematical tools for *A*_*in*,1_ *≥ A*_*in*,2_ *≥* 0.

#### 2.3.1 Linear networks

For networks having linear nodes and linear connectivity *I*_*syn,k*_ = *−* (Γ_*k*,1_*v*_1_ +Γ_*k*,2_ *v*_2_), for *k* = 1, 2, the voltage response of each cell has the form *V*_*out,k*_(*t*; *f*) = *A*_*out,k*_(*f*) sin(Ω *t* − Φ_*ntwk,k*_(*f*)). The network (amplitude) impedance for each node is defined as

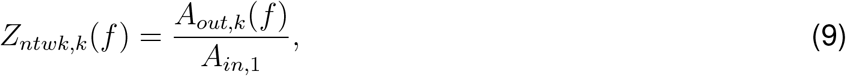

for *k* = 1, 2 and the network phase Φ_*ntwk,k*_(*f*) (phase-shift) is the difference between the peaks of *I*_*in*,1_ and *V*_*out,k*_ normalized by the period (for each value of the input frequency *f*). These are the component of the complex network impedance presented in the Supplementary Material S4 (for networks of *N* cells with 2D linear dynamics, linear connectivity and sinusoidal inputs to all cells in the network). In (9) we chose to normalize the network impedance by the input amplitude to the first node in the network, which we assume is the one receiving the oscillatory input with the highest amplitude.

For two-cell networks, the complex network impedances are given by (see Supplementary Material S4)

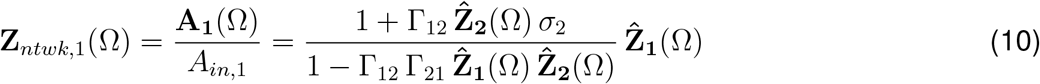

and

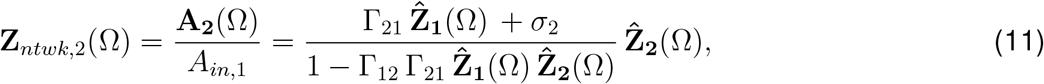

where

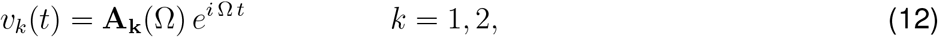

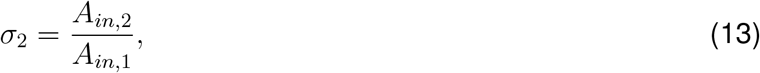

and 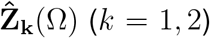 are the extended (complex) impedances of the individual nodes in the network where the parameter *−g*_*L,k*_ is substituted by *−g*_*L,k*_ + Γ_*kk*_ in eq. (S15) in the Supplementary Material S2 (the self-connectivity term Γ_*kk*_*v*_*k*_ is incorporated into the autonomous part of the corresponding nodes *k*).

If *A*_*in*,2_ = 0 (*σ*_2_ = 0),

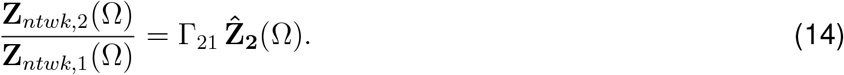

#### 2.3.2 Network resonance and phasonance

The notions of resonance and phasonance extend naturally to networks. We use the notation *f*_*ntwk,res,k*_ and *f*_*ntwk,phas,k*_ (*k* = 1, 2) to refer to the resonant and phasonant frequencies, respectively, of cell *k* when it is part of a network. We note that looking at the response of each cell in a network is not the only option to characterize the network response to oscillatory inputs. For example, one can choose a population metric by taking, for example, the average network voltage response across the population (of part of a population in a large network).

#### 2.3.3 Nonlinear networks

For nonlinear networks with graded synaptic connections, we adapt the envelope amplitude response *Z*^*ENV*^ (*f*) defined in the previous subsection and define the network impedance (amplitude) as

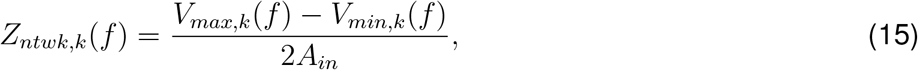

for *k* = 1, 2 where *V*_*max,k*_(*f*) and *V*_*min,k*_(*f*) are the peak and trough envelope profiles, respectively defined as the maximum and minimum of the steady-state oscillatory voltage response of cell *k* as a function of the input frequency *f*. The network phase Φ_*ntwk,k*_(*f*) is defined as the difference between the peaks of *I*_*in*,1_ and *V*_*out,k*_ normalized by the period, for each value of the input frequency *f*.

#### 2.3.4 Network impedance vs. peak envelope profiles

The impedance amplitude is symmetric for linear, but not for nonlinear systems. Therefore, while the impedance amplitude captures the frequency content of the voltage response signal, it does not necessarily describe the frequency-dependent properties of the peak and trough envelope profiles. Because neuronal signal occurs via voltage threshold-like mechanisms, we will look at the peak envelope profiles *V*_*max,k*_(*f*) for *k* = 1, 2, and adapt the notion of resonance to this profile. More specifically, a cell in the network exhibits resonance if the *V*_*max,k*_ peaks at some nonzero (resonant) frequency *f*_*res,k*_.

### 2.4 Communication coefficient: Frequency-dependent profiles

#### 2.4.1 Amplitude and phase

To analyze the frequency dependent properties of the communication of information between the two nodes in the network in response to oscillatory inputs, we define the amplitude communication coefficient from the cell receiving the input (cell 1) to the coupled cell (cell 2)

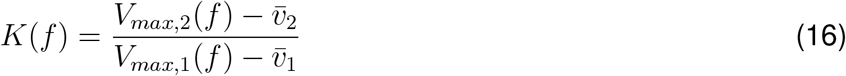

where 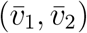 is the fixed point of system (1)-(2), and the phase communication coefficient

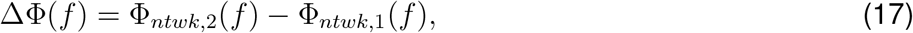

consisting of the phase difference between the voltage responses of the two cells. For linear systems,

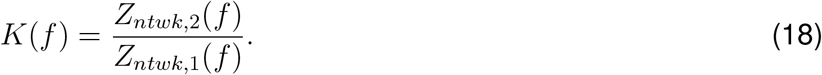

The communication coefficient *K*(*f*) is a frequency-dependent adaptation of the steady-state coupling coefficient used in networks connected trough gap junctions [53–57] in the presence of constant (DC) inputs, which has been used to measure the strength of the connection in response to the constant inputs. Here, we extend the concept of coupling coefficient to neurons connected with chemical synapses (nonlinear) and receiving periodic inputs over a frequency range. Because the response to oscillatory inputs has amplitude and phase, we also extend our metrics to include the phase communication coefficient ΔΦ(*f*), which has not been considered before.

In fact, for linear systems, *K*(*f*) and ΔΦ(*f*) are the components of the following complex quantity

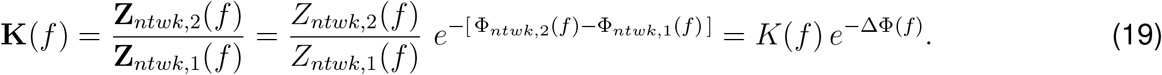

The communication coefficient *K*(*f*) allows us to study how the communication of information between the two cells depends on the interplay of the input frequency, the intrinsic properties of the nodes and the synaptic connectivity. From a different perspective, *K*(*f*) captures the frequency-dependent properties of the effective connectivity strength in terms of the properties of the network components. The phase communication coefficient ΔΦ gives us information about the synchrony between response peaks of the two cells.

#### 2.4.2 *K*-resonance and ΔΦ-phasonance

*K*-resonance refers to the ability of the frequency-dependent communication coefficient *K*(*f*) to exhibit a peak at a non-zero input frequency *f*_*res,K*_. Similarly, *K*-antiresonance refers to the ability of the frequency-dependent communication coefficient *K*(*f*) to exhibit a trough at a non-zero input frequency *f*_*ares,K*_. ΔΦ-phasonance refers to the ability of the frequency-dependent phase communication coefficient ΔΦ(*f*) to vanish for a nonzero input frequency *f*_*phas*,ΔΦ_.

These quantities are in fact a property of the connected cell pair. Here, we refer to them indistinctly as a property of the cell pair or the corresponding *K* and ΔΦ profiles.

From eq. (16), a necessary condition for the existence of *K*-resonance (or *K*-antiresonance) is given by

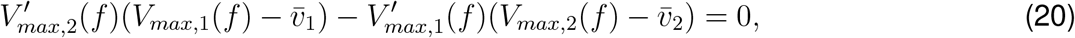

for some value *f >* 0.

#### 2.4.3 Attributes

We characterize the *K* profile by using two attributes: (i) the resonant frequency *f*_*res,K*_, defined as the peak frequency of *K*(*f*), and (ii) the resonance amplitude *Q*_*K*_ = *K*_*max*_ *− K*(0). If the *K* profile does not exhibit resonance, then *f*_*res,K*_ = *Q*_*K*_ = 0. If the network exhibits *K*-antiresonance, we use the attribute *f*_*ares,K*_. We characterize the ΔΦ profile by using the phasonant frequency *f*_*phas*,ΔΦ_.

In Table 1, we summarize the notation for resonant and phasonant frequencies (for cells and communication coefficient) used throughout the paper.

**Table 1:** Notation for resonant and phasonant frequencies

|  | Symbol | Meaning |
| --- | --- | --- |
| *Single linear cell* | $f_{res}$ | resonant frequency of impedance $Z$ (and $V_{max}$) |
| | $f_{phas}$ | phasonant frequency of $\Phi$ |
| *Linear networks* | $f_{ntwk,res,k}$ | resonant frequency of impedance $Z_{ntwk,k}$ (and $V_{max,k}$) |
| | $f_{ntwk,phas,k}$ | phasonant frequency of $\Phi_{ntwk,k}$ |
| *Non-linear networks* | $f_{res,k}$ | resonant frequency of $V_{max,k}$ |
| | $f_{res,k}^0$ | resonant frequency of the disconnected cell $V_{max,k}^0$ |
| | $f_{phas,k}$ | phasonant frequency of $\Phi_{ntwk,k}$ |
| | $f_{phas,k}^0$ | phasonant frequency of the disconnected cell $\Phi_{ntwk,k}^0$ |
| *K and $\Delta\Phi$* | $f_{res,K}$ | resonant frequency of coefficient $K$ |
| | $f_{ares,K}$ | antiresonant frequency of coefficient $K$ |
| | $f_{phas,\Delta\Phi}$ | phasonant frequency of coefficient $\Delta\Phi$ |

#### 2.4.4 Normalized communication coefficient *K*^*\**^

Understanding how the communication coefficient profiles *K*(*f*) are shaped by the network building blocks and how the latter contribute to the generation of preferred communication frequency ranges, requires comparison between the network peak profiles *V*_*max*,1_(*f*) and *V*_*max*,2_(*f*), and how they are shaped by the network building blocks. In order to facilitate this comparison we normalize these quantities

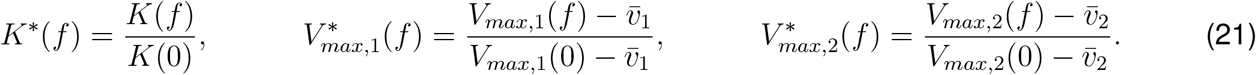

These normalized quantities are related by

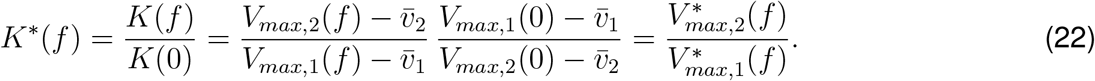

### 2.5 Numerical simulations

To compute the numerical solutions we used the modified Euler method (Runge-Kutta, order 2) [74] with a time step Δ*t* = 0.01 ms. Smaller values of Δ*t* have been used to check the accuracy of the results. All neural models and metrics, including phase-plane analysis, were implemented by self-developed Python (Python Software Foundation, https://www.python.org/) and MATLAB (The Mathworks, Natick, MA) routines. These routines are available at https://github.com/andrealbel/freq_communication.git.

## 3 Results

### 3.1 Network voltage response to oscillatory input currents: linear nodes and linear connectivity

We first discuss the simplified scenario where the network connectivity in the linear system (1)-(2) is linear

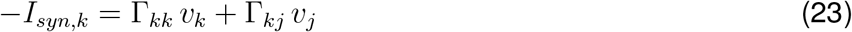

with Γ_*kj*_ constant (*j, k* = 1, 2). This system is amenable to analytical calculations (see Section 2.3.1 for details) and allows us to develop analytical, dynamical systems tools and test ideas that will be then extended to the more realistic nonlinear networks. We assume *A*_*in*,1_ *≥ A*_*in*,2_ (without loss of generality) and choose *A*_*in*,1_ as the normalization constant.

The network impedances are given by eqs. (10)-(11). They are functions of the extended impedances 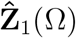 and 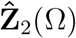, which are presented in the Supplementary Material S2 (with *g*_*L,k*_ substituted by *g*_*L,k*_ *−* Γ_*kk*_, *k* = 1, 2). In the absence of self-connections (Γ_11_ = Γ_22_ = 0), 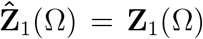 and 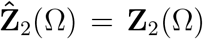 (the extended impedances of the individual cells are equal to their impedances).

By substituting the network impedances (10)-(11) into eq. (18), the amplitude communication coefficient reads

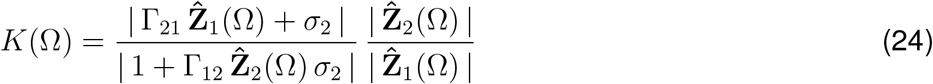

where *σ*_2_ = *A*_*in*,2_*/A*_*in*,1_. That is, the frequency-dependent properties of the amplitude and phase communication coefficients result from the interplay of the extended impedances of the participating cells, and do not directly reflect the frequency-dependent properties of the network impedance (discussed below).

If only cell 1 receives an oscillatory inputs (*A*_*in*,2_ = *σ*_2_ = 0), then

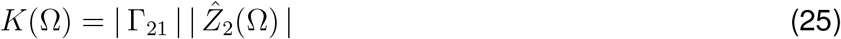

and, from eqs. (14) and (19),

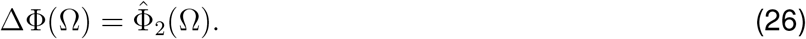

That is, the frequency-dependent properties of the amplitude and phase communication coefficients are inherited from the extended impedance and phase profiles of cell 2 (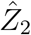 and 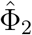) and are independent of the frequency-dependent properties of the directly perturbed cell 1 (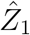 and 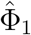).

According to this, for example, a linear network of passive cells (1D) does not exhibit *K*-resonance nor ΔΦ-phasonance regardless of which cell receives the oscillatory input and whether the cells exhibit network resonance or phasonance. A linear network of a resonator (2D) recurrently connected to a passive cell (1D), where for simplicity we assume there are no self-connections, shows *K*-resonance provided the oscillatory input arrives at the passive cell, but not when the oscillatory input arrives at the resonator. Similarly, if the 2D cell shows phasonance when disconnected, then the linear network shows ΔΦ-phasonance if the input arrives at the passive cell, but not otherwise.

In the next section we analyze in detail the frequency-dependent properties of a linear two-cell network of 1D cells. We first illustrate that although the network exhibits (network) resonance and phasonance in certain parameter regimes, it does not exhibits *K*-resonance nor ΔΦ-phasonance, as expected. We then develop a dynamical systems approach using an extended version of the phase-plane diagram to analyze the dynamic mechanisms that govern the generation of *K*-resonance and the differences between *K*-resonance and network resonance. We use this approach for the analysis of nonlinear networks where analytical calculations are not possible.

### 3.2 Two-1D-cell linear networks

The dynamics of a two-1D-cell linear network with an input arriving only at cell 1 are described by

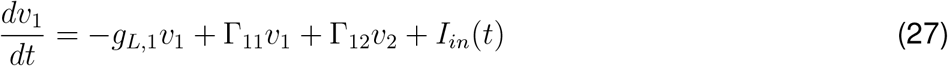

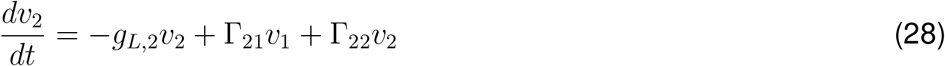

where *I*_*in*_(*t*) is given by (5) and we omit the membrane capacitance (*C*_1_ = *C*_2_ = 1).

The individual nodes are passive cells, and therefore they are low-pass filters and the Φ-profiles are positive and increasing (see Supplementary Material S2).

#### 3.2.1 Intrinsic properties revisited

The eigenvalues of the system (27)-(28) are given by (see eq. (S27) in the Supplementary Material)

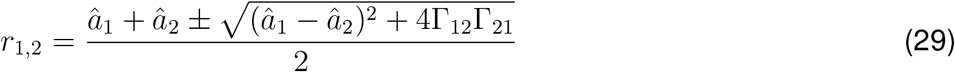

where *â*_*k*_ = *−g*_*L,k*_ + Γ_*kk*_, for *k* = 1, 2. Stability requires *â*_1_ + *â*_2_ *<* 0 and *â*_1_*â*_2_ *>* Γ_12_Γ_21_ or, more specifically,

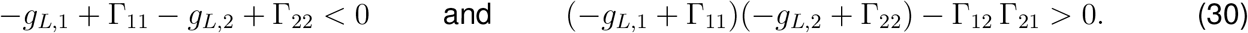

While stability of the individual cells requires *g*_*L*,1_ *>* 0 and *g*_*L*,2_ *>* 0, the stability of the network allows for *g*_*L*,1_ *<* 0 or *g*_*L*,2_ *<* 0 (or both), provided the self-connectivity parameters (Γ_11_ and Γ_22_) are negative enough. In the following, the intrinsic parameters *g*_*L*,1_ and *g*_*L*,2_ are assumed to absorb the self-connectivity parameters Γ_11_ and Γ_22_, respectively, without loss of generality and we will consider only positive values of both *g*_*L*,1_ and *g*_*L*,2_. The two conditions (30) become

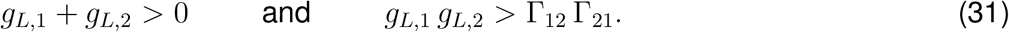

The second condition in eq. (31) imposes a restriction on the cross-connectivity parameters when they have the same sign, but not when their signs are opposite. In this case, the network exhibits damped oscillations when (*g*_*L*,1_ *− g*_*L*,2_)^2^ + 4Γ_12_Γ_21_ *<* 0, with natural frequency

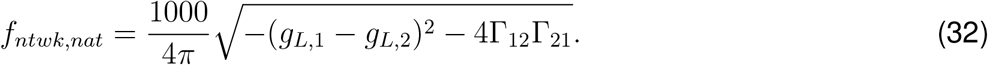

#### 3.2.2 Network resonance and phasonance

The network impedance and phase profiles and the network resonant and phasonant frequencies can be computed analytically. From (10)-(11) with Γ_11_ = Γ_22_ = 0 (without loss of generality) and *σ*_2_ = 0, the network impedance profiles are given by

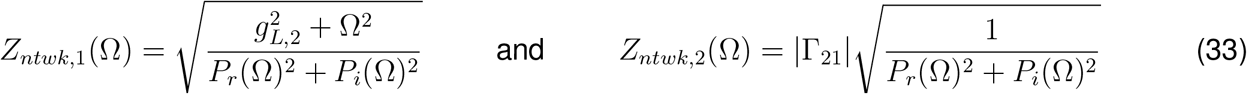

and the network phase profiles are given by

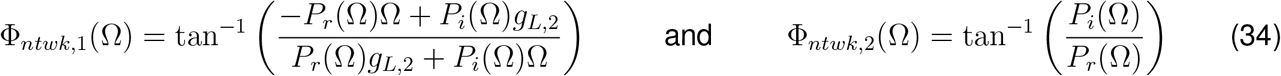

where

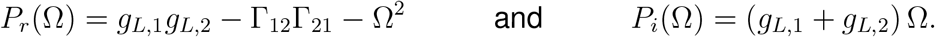

From (33), if both cells exhibit network resonance, the corresponding resonant frequencies are given by

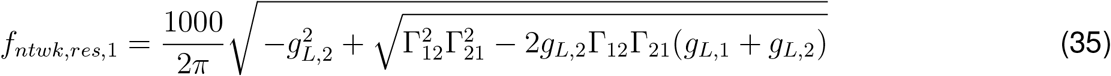

and

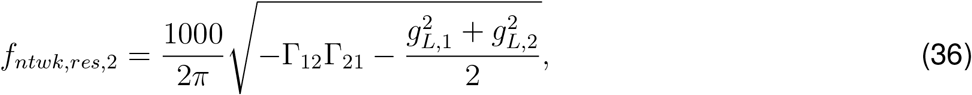

respectively. If cell 1 exhibit network phasonance, the phasonant frequency is given by

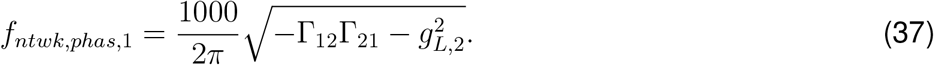

The cell 2 can never exhibit network phasonance since *P*_*i*_(Ω) in the second equation in (34) vanishes only for Ω = 0 (*f* = 0).

Network resonance in cells 1 and 2 and network phasonance in cell 1 require Γ_12_ and Γ_21_ to have different signs. This is straightforward from *f*_*ntwk,res*,2_ (36) and *f*_*ntwk,phas*,1_ (37). If this condition is not satisfied for *f*_*ntwk,res*,1_ (35), then Γ_12_Γ_21_ *−* 2*g*_*L*,2_ (*g*_*L*,1_ +*g*_*L*,2_) *>* 0, but the second stability condition in (31) implies 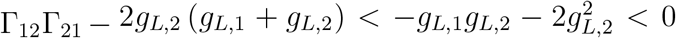. Note that the characteristic (natural, resonant and phasonant) frequencies do not necessarily coincide. They do when *g*_*L*,1_ = *g*_*L*,2_ = 0.

Figs. 3 illustrates the network impedance profiles for two-1D-cell mutually inhibitory and excitatory-inhibitory networks with input only to cell 1 (*A*_*in*,2_ = 0). In the excitatory-inhibitory network we consider two representative scenarios: (i) both cells 1 and 2 are network low-pass filters (Figs. 3-B1) for low enough values of the cross-connectivity parameters, and (ii) both cells 1 and 2 exhibit network resonance (Figs. 3-C1) for large enough values of the cross-connectivity parameters. Fig. 4 shows the phase profiles for the networks considered in Fig. 3. In the mutually inhibitory network cell 2 is delayed as compared to cell 1. For the excitatory-inhibitory scenarios: (i) both cells exhibit a delayed response for all values of the input frequency *f* (Fig. 4-B), and (ii) cell 1 exhibits network phasonance, while cell 2 has a delayed response for all values of the input frequency *f* (Fig. 4-C). There is an intermediate representative scenario (not shown) between these two where cell 1 exhibits network resonance, while cell 2 is a network low-pass filter and both cells have a delayed response for all values of the input frequency *f*.

**Figure 3:**
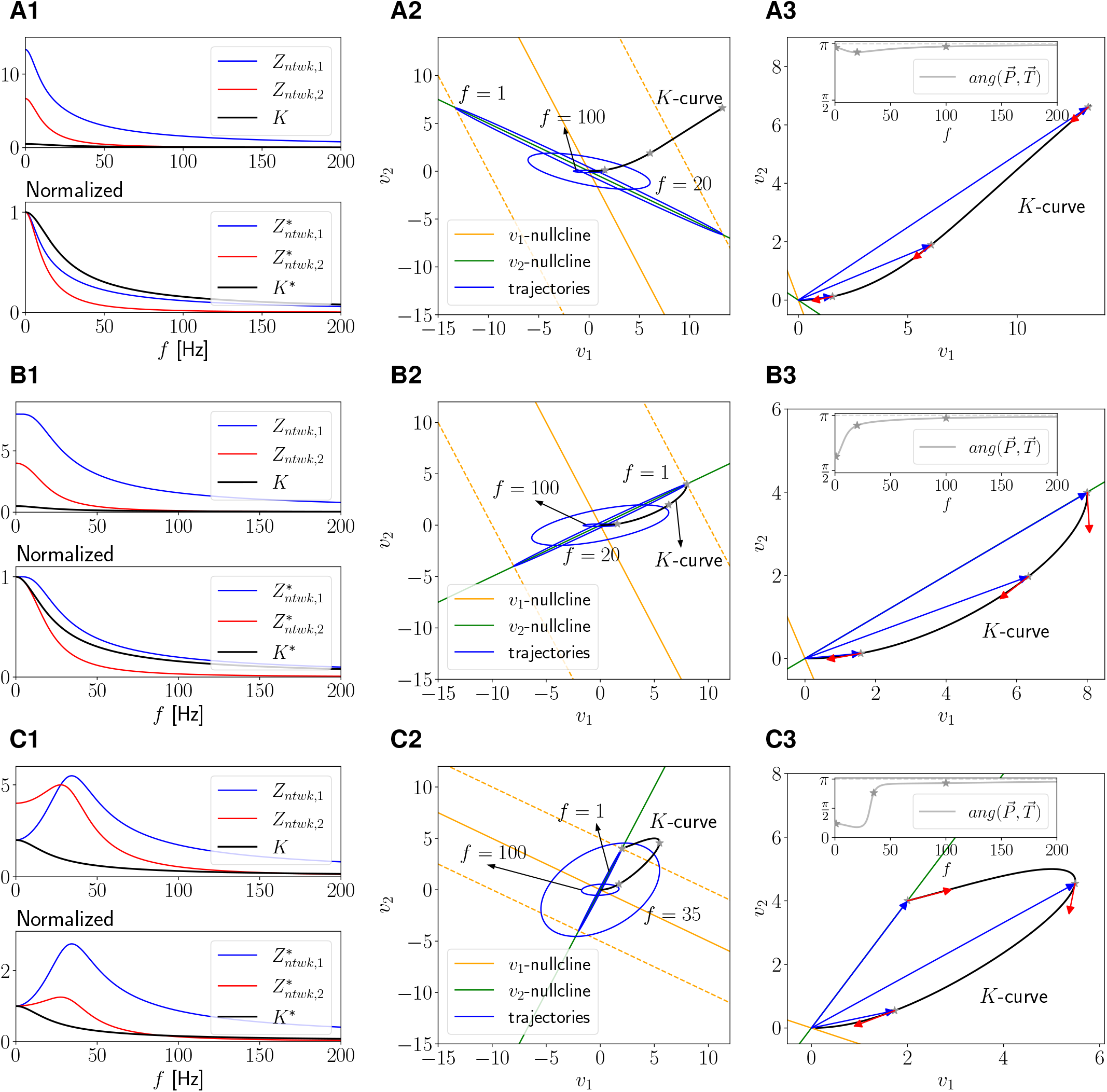
Response of a linear network of two 1D cells to oscillatory inputs. We used the linear system (27)-(28) receiving an oscillatory inputs (5) only to cell 1 (*A*_*in*,1_ = *A*_*in*_ and *A*_*in*,2_ = 0). The self-connectivity parameters are assumed to be absorbed by the corresponding linearized leak conductances and therefore we use Γ_11_ = Γ_22_ = 0 without loss of generality. **Left column**. Network impedances and communication coefficient *K* = *Z*_*ntwk*,2_*/Z*_*ntwk*,1_ (top). Normalized network impedances and communication coefficient (bottom). **Middle column**. Superimposed phase-plane diagram projections onto the *v*_1_-*v*_2_ plane. The solid orange and green lines are the *v*_1_ and *v*_2_ nullclines of the autonomous system, respectively, given by eqs. (40). The dashed orange lines are the extended *v*_1_-nullclines for the autonomous system for constant inputs equal to *±A*_*in*_, given by eq. (41). The blue curves are the (projections of the) response limit cycle trajectories for representative values of *f*. The black curve is the *K*-curve, it joins the peak values of the response limit cycles in the *v*_1_ and *v*_2_ directions and correspond to the network impedances in the left column. The gray stars correspond to the representative response limit cycles (blue curves). **Right column**. Magnified view of the *K*-curve, with the three representative position (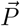, blue) and tangent (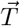, red) vectors. The tangent vectors were rescaled to improve the visualization. **Inset**. Angle between 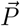 and 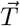 for the range of values of *f* in left panels. We used the following parameter values: *g*_*L*,1_ = *g*_*L*,2_ = 0.1, *A*_*in*_ = 1. **A**. Γ_12_ = *−*0.05, Γ_21_ = *−*0.05. **B**. Γ_12_ = *−*0.05, Γ_21_ = 0.05. **C**. Γ_12_ = *−*0.2, Γ_21_ = 0.2.

**Figure 4:**
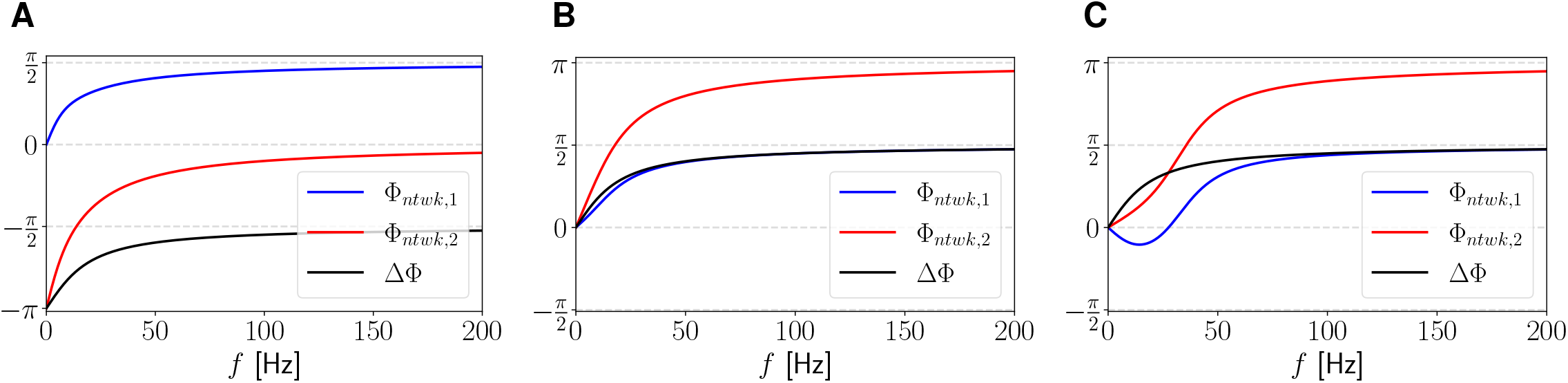
Response of a linear network of two 1D cells to oscillatory inputs. Phases. Network phase profiles for cells 1 and 2 (solid) and phase difference profile (black). We used the linear system (27)-(28) receiving an oscillatory inputs only to cell 1 (*A*_*in*,1_ = *A*_*in*_ and *A*_*in*,2_ = 0). We used the following parameter values: *g*_*L*,1_ = *g*_*L*,2_ = 0.1, *A*_*in*_ = 1. **A**. Γ_12_ = *−*0.05, Γ_21_ = *−*0.05. **B**. Γ_12_ = *−*0.05, Γ_21_ = 0.05. **C**. Γ_12_ = *−*0.2, Γ_21_ = 0.2.

The results discussed above do not change qualitatively if these two signs are inverted, the cells are not identical (*g*_*L*,1_ ≠ *g*_*L*,2_) or the cross-connectivity parameters have different absolute values. The relative size of the network impedance profiles depend on the ratio Γ_12_*/*Γ_21_, the stronger Γ_21_ (the connectivity from cell 1 to cell 2) for fixed values of Γ_12_, the larger *Z*_*ntwk*,2_ relative to *Z*_*ntwk*,1_ (not shown). In the Supplementary Material (S3), we illustrate the dependence of the characteristic frequencies *f*_*ntwk,nat*_, *f*_*ntwk,res*,1_, *f*_*ntwk,res*,2_ and *f*_*ntwk,phas*,1_ on the cross-connectivity parameters Γ_12_ and Γ_21_ (Fig. S1)

#### 3.2.3 Absence of *K*-resonance and ΔΦ-phasonance

From eqs. (25), (26), the linear system (27)-(28) considered here (*A*_*in*,2_ = *σ*_2_ = 0), inherits the frequency-dependent properties of the *K*- and ΔΦ-profiles from the extended impedance and phase profiles of cell 2. Specifically, from eq. (S21) in the Supplementary Material S2 with *g*_*L*,2_ substituted by *g*_*L*,2_ *−* Γ_22_

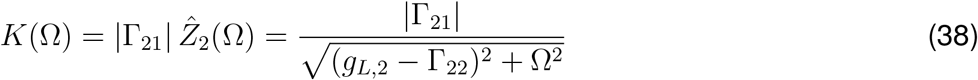

and

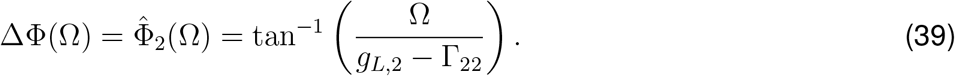

The *K*-profiles are monotonically decreasing in all cases (Fig. 3-A1, B1 and C1) and the ΔΦ-profiles are monotonically increasing and different from zero in all cases (Fig. 4). This, in spite of the presence of network resonance and phasonance in some parameter regimes. The two cells never peak simultaneously, which would require ΔΦ = 0. However, being linear systems, they are phase locked for all values of *f*.

### 3.3 Dynamical systems tools for the analysis of the frequency-dependent properties of the communication coefficient *K*(*f*)

For nonlinear networks, the *K*(*f*) and ΔΦ(*f*) cannot be computed analytically. Here we extend the methods introduced in [27, 37] to develop dynamical system tools for the qualitative analysis of both the dynamic mechanisms governing the generation of the *K*- and ΔΦ-profiles and their dependence on the model parameters. These tools allow us to better understand how *K*(*f*) is shaped by the network building blocks by looking at the interaction between the vector field of the autonomous (unforced) system and the oscillatory input that temporally modulates this vector field over a range of input frequencies. For simplicity we describe the method in a two-cell linear network of 1D cells. The extension to non-linear networks is straightforward. Further extension, for example, to networks having nodes with higher-dimensional dynamics requires the use of projections for visualization purposes.

Specifically, we consider the two-cell linear network (27)-(28) presented in Section 3.2 where the sinusoidal input (5) arrives only at cell 1 (*A*_*in*,2_ = 0) and Γ_11_ = Γ_22_ = 0 (without loss of generality as discussed above).

#### 3.3.1 The phase-space diagram

The *v*_1_- and *v*_2_-nullclines for the autonomous system are given by

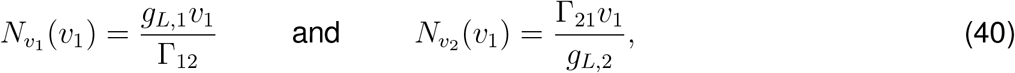

respectively. The corresponding extended (maximally displaced) *v*_1_-nullclines are defined by

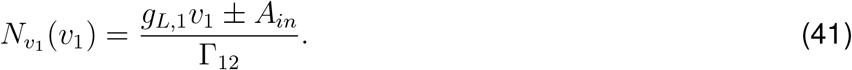

Following [27, 37], we rescale time by defining *t ← ft*. The linear system (27)-(28) becomes

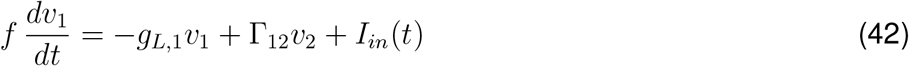

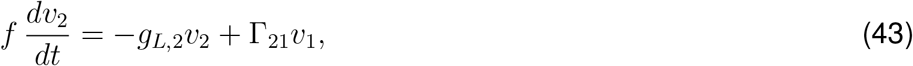

where *I*_*in*_(*t*) is now the sinusoidal input (5) with *f* substituted by 1. In other words, the oscillatory input has unit frequency for all values of *f* and the effect of the input frequency in the oscillatory input was translated to the speed of the response trajectories. The *v*_1_- and *v*_2_-nullclines for the autonomous system remain unchanged by this rescaling, which affects only the speed at which the trajectories evolve for each value of *f*.

Fig. 3-A shows a representative example. Fig. 3-A1 (top) shows the network impedances (*Z*_*ntwk*,1_ and *Z*_*ntwk*,2_) and the communication coefficient *K*. Because of the symmetry of the responses in linear systems,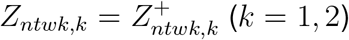 and because *A*_*in*_ = 1, *Z*_*ntwk,k*_ = *V*_*max,k*_ (*k* = 1, 2). Fig. 3-A1 (bottom) shows the normalized network impedances and communication coefficient. Fig. 3-A2 shows the phase-plane diagram corresponding to Fig. 3-A1. It is in fact the superimposed projections of the 3D phase-space diagrams onto the *v*_1_-*v*_2_ plane (the third dimension is *t*) for representative values of *f*. The solid orange and green lines are the *v*_1_- and *v*_2_-nullclines of the autonomous system, respectively, given by eqs. (40). The dashed orange lines are the maximally displaced *v*_1_-nullclines for the autonomous system (for constant inputs equal to *± A*_*in*_), given by eq. (41). The blue curves are the (projections of the) response limit cycle trajectories for representative values of *f*.

#### 3.3.2 Dynamic mechanisms of generation of the network voltage response profiles: “moving null-clines” and response limit cycles

As *t* progresses, the *v*_1_-nullcline is interpreted as moving cyclically in between the maximally displaced *v*_1_-nullclines (dashed-orange) reaching them at the oscillatory input’s peaks and troughs. As the *v*_1_-nullcline moves, so does the fixed-point. The response limit cycle trajectories track the motion of the fixed-point with a speed that depends on the input frequency *f*. Thus, the response limit cycles are shaped by the combined effect of the input frequency *f* and the properties of the underlying vector field (independent of *f*), which are partially captured by the nullclines of the autonomous system. The amplitude of the response limit cycles may decrease with increasing values of f (low-pass filter, no resonance), may increase for values of f within some range and then decrease for larger values of f (band-pass filter for one or both variables), or have more complex shapes. We refer the reader to [27, 37] for more details.

#### 3.3.3 The *K*-curve

The peak values *V*_*max*,1_ and *V*_*max*,2_ of the response limit cycles in the *v*_1_ and *v*_2_ directions form the so-called *K*-curve (in Fig. 3-A2 and A3, the black curve for the range of values of *f* in panel A1 and the three gray stars for the three representative response limit cycles). The *K*-curve is parametrized by *f* and its shape captures the relative change of *V*_*max*,1_ and *V*_*max*,2_ with *f*. As *f* increases, the *K*-curve moves from the mirror image of the intersection between the *v*_2_-nullcline and the right-most extended *v*_1_-nullcline (*f →* 0) to the origin (*f →∞*). Since the communication coefficient *K*(*f*) describes the relative changes of the maximum profiles, the *K*-curve is a geometric representation of the *K*-profile that allows us to analyze its properties and the mechanism by which it is generated from a dynamical systems point of view.

The effects of changes of the model parameters (*g*_*L*,1_, *g*_*L*,2_, Γ_12_ and Γ_21_) on the network impedances and the communication coefficient can be interpreted by looking at how they affect the nullclines and in turn the shapes of the response limit cycles. In the following sections we first use this dynamical systems tool to interpret our results for linear networks and then we expand this method to analyze nonlinear networks with linear nodes and chemical synapses, and to explain the emergence of communication of information resonance and phasonance in the network in response to oscillatory inputs.

#### 3.3.4 A geometric necessary condition for the generation of *K*-resonance

We define the position and tangent vectors of the *K*-curve as

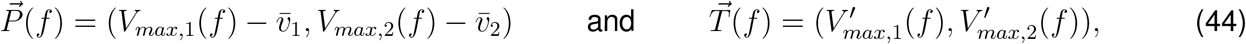

respectively. From the necessary condition (20), the alignment of 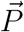 and 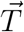 for a given frequency *f* could indicate the presence of resonance (or antiresonance) at this frequency. Geometrically, a change of curvature in the *K*-curve is a a necessary condition for the alignment of 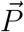 and 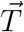 and therefore, for the presence of resonance in the *K* profile.

In Fig. 3-A, both *Z*_*ntwk*,1_ and *Z*_*ntwk*,2_ (and so *V*_*max*,1_ and *V*_*max*,2_) are decreasing functions of *f* (low-pass filters). The curvature of the *K*-curve does not change its sign since *Z*_*ntwk*,2_ decreases faster than *Z*_*ntwk*,1_ with increasing values of *f*, and thus, 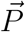 and 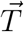 are not aligned for any frequency value *f*. Therefore, the *K* coefficient is monotonically decreasing and the *K*-curve captures this behavior.

#### 3.3.5 Two-1D-cell network: The *K*-curve does not change concavity type even in the presence of network resonance

Consistent with our findings in Section 3.2, the curvature of the *K*-curve in the *v*_1_-*v*_2_ phase-plane diagram (black curve in Fig. 3, middle and right panels) does not change sign. Moreover, the position and tangent vectors are not aligned for any input frequency *f* (Inset in Fig. 3, right panels). Specifically, if both *Z*_*ntwk*,1_ and *Z*_*ntwk*,2_ are decreasing functions of *f* (low-pass filters) and *Z*_*ntwk*,2_ decreases faster than *Z*_*ntwk*,1_ with increasing values of *f*, then *K* is monotonically decreasing (Fig. 3-A and B). If for some range of values of *f* both *Z*_*ntwk*,1_ and *Z*_*ntwk*,2_ are increasing functions of *f* and *Z*_*ntwk*,1_ increases faster than *Z*_*ntwk*,2_, then *K* is also monotonically decreasing within that range (Fig. 3-C).

The effects of changes in the model parameters (*g*_*L*,1_, *g*_*L*,2_, Γ_12_ and Γ_21_) on the network impedances for both cells and the communication coefficient can be interpreted by looking at how they affect the nullclines in the phase-plane diagram, which, in turn, affect the shapes of the response limit cycles and the *K*-curve. Increasing values of *g*_*L*,1_ increase the slope of the *v*_1_-nullcline, while increasing values of *g*_*L*,2_ decrease the slope of the *v*_2_-nullcline. Increasing values of |Γ_12_| decrease the slope of the *v*_1_-nullcline, while increasing values of |Γ_21_| increase the slope of the *v*_2_-nullcline. Changes in the signs of Γ_12_ and Γ_21_ change the directions of the *v*_1_- and *v*_2_-nullclines, accordingly (e.g., compare Figs. 3-A and B). In addition, decreasing values of *g*_*L*,1_ cause the *v*_1_-nullcline to be more shallow and therefore the intersection between the *v*_2_-nullcline and the right-most extended *v*_1_-nullcline occurs for a higher value of *v*_1_ and *v*_2_, thus increasing the network impedances and amplifying the response (not shown). Decreasing values of *g*_*L*,2_ also cause an amplification of the response by increasing the slope of the *v*_2_-nullcline and therefore increasing the intersection value between the *v*_2_-nullcline and the right-most extended *v*_1_-nullcline (not shown). Increasing levels of mutual inhibition (increasing negative values of Γ_12_ and Γ_21_) cause the *v*_1_ nullcline to be shallower and the *v*_2_-nullcline to be steeper, thus increasing the intersection value between the *v*_2_-nullcline and the right-most extended *v*_1_-nullcline and causing an amplification of the response (not shown). However, none of these changes in the model parameters cause the curvature of the *K*-curve to change sign or an alignment between position and tangent vectors for an intermediate frequency. Therefore, the *K* coefficient is always monotonically decreasing.

### 3.4 Two-1D-cell networks with nonlinear synaptic connectivity: Emergence of *K*-resonance

From our previous discussion, neither *K*-resonance nor ΔΦ-phasonance occur in two-1D-cell linear networks. Here we investigate the conditions under which these phenomena emerge as the result of the presence of the nonlinear connectivity.

From eqs. (1)-(2), the dynamics of a recurrently connected two-1D-cell network are given by

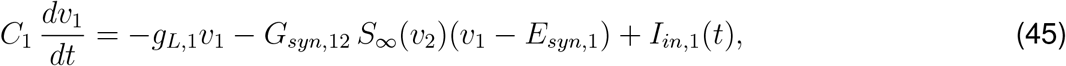

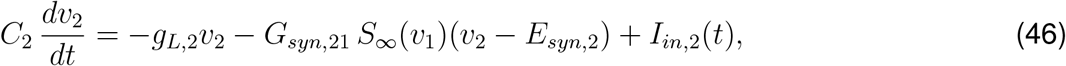

where *I*_*in*,1_(*t*) and *I*_*in*,2_(*t*) are given by (5) with amplitude *A*_*in*,1_ and *A*_*in*,2_, respectively, and the graded synaptic function *S*_*∞*_(*v*) is given by (4) with *v*_*hlf*_ = 0, *v*_*slp*_ = 1. In our analysis and simulations we use *A*_*in*,2_ = 0 (the oscillatory inputs arrives only to cell 1) and we refer to *A*_*in*,1_ simply as *A*_*in*_.

As for the linear networks, the individual nodes are low-pass filters and the Φ-profiles are positive and increasing. We discuss four scenarios: (i) mutual inhibition, (ii) mutual excitation, (iii) inhibition from cell 1 to cell 2 and excitation from cell 2 to cell 1, and (iv) excitation from cell 1 to cell 2 and inhibition from cell 2 to cell 1 (see diagrams in Fig. 1-A). We restrict our analysis to the parameter regimes where the autonomous (unperturbed) network exhibits a single stable equilibrium. The analysis of networks with more complex dynamics is beyond the scope of the present work.

#### 3.4.1 Linearized network

The linearization of system (45)-(46) around its fixed point 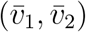 reads

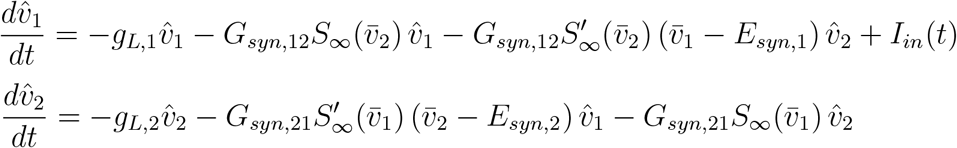

where

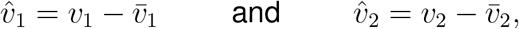

and we omit the membrane capacitance (*C*_1_ = *C*_2_ = 1). To the linear level of description, the nonlinear cross-connectivity terms in eqs. (45)-(46) can be explicitly unfolded into a self-connectivity contribution and a “true” cross-connectivity contribution. The analysis of this linearized systems is a direct application of the results presented in the previous section. These results extend to weakly nonlinear systems and nonlinear systems with inputs that activate the nonlinearities only weakly.

#### 3.4.2 Development of *K*-resonance as increasing values of *A*_*in*_ activate the network nonlinearities

Figs. 5 and 6 show representative scenarios for mutually inhibitory and excitatory-inhibitory networks, respectively and increasing values of *A*_*in*_ (from panels A to C). For the mutually inhibitory network, the two cells are network low-pass filters (Figs. 5, left). For low enough values of *A*_*in*_, the communication coefficient *K*(*f*) develops a peak (resonance) whose normalized value increases with increasing values of *A*_*in*_ (Figs. 5-A1 to C1, bottom). This reflects a switch in the relative rates at which *V*_*max*,1_(*f*) and *V*_*max*,2_(*f*) decrease. The *K*-resonant frequency increases as *A*_*in*_ increases, as the intersection between the two normalized peak profiles moves to the right (Figs. 5-B1 and C1).

**Figure 5:**
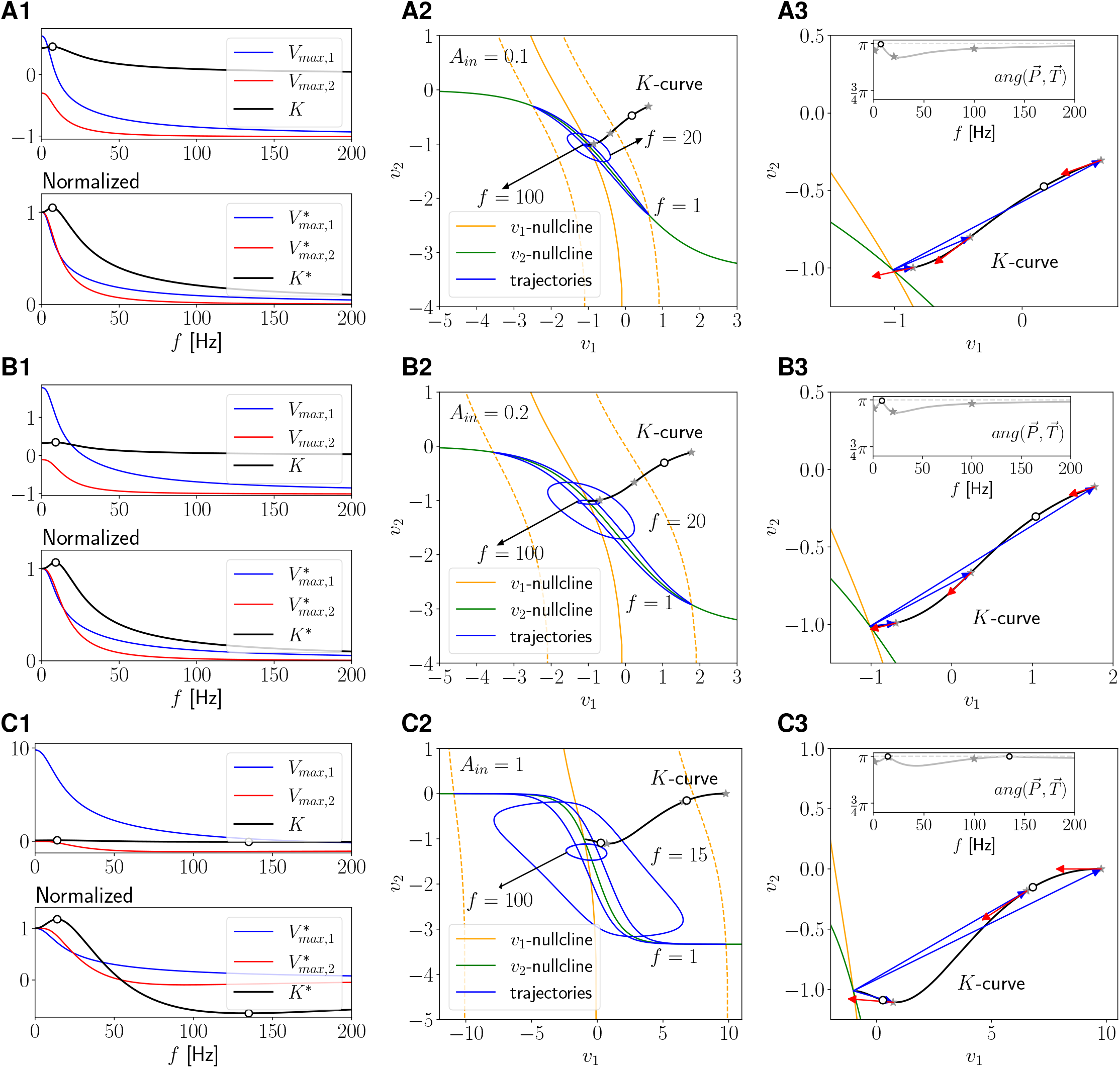
Response of a nonlinear mutually inhibitory network of two 1D cells to oscillatory inputs. We used the system (45)-(46) receiving an oscillatory inputs only to cell 1 (*A*_*in*,1_ = *A*_*in*_ and *A*_*in*,2_ = 0). The self-connectivity parameters (if they exist) are considered to be absorbed by the autonomous linear terms (*G*_*syn*,11_ = *G*_*syn*,22_ = 0) and *C*_1_ = *C*_2_ = 1. **Left column**. Network peak (*V*_*max*,1_ and *V*_*max*,2_) and communication coefficient (*K*) profiles (16) (top). Normalized profiles (bottom). The dots indicate the presence of *K*-resonance and antiresonance. **Middle column**. Superimposed phase-space diagram projections onto the *v*_1_-*v*_2_ plane. The solid orange and green lines are the *v*_1_ and *v*_2_ nullclines of the autonomous system, respectively, given by eqs. (47) and (48). The dashed orange lines are the extended *v*_1_-nullclines for the autonomous system for constant inputs equal to *±A*_*in*_, given by eq. (49). The blue curves are the (projections of the) response limit cycle trajectories for representative values of *f*. The black curve is the *K*-curve joining the peak values of the response limit cycles in the *v*_1_ and *v*_2_ directions. The gray stars correspond to the representative response limit cycles (blue curves). **Right column**. Magnified view of the *K*-curve, with representative position (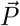, blue) and tangent (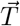, red) vectors. **Inset**. Angle between 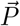 and 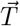 for the range of values of *f* in left panels. We used the following parameter values: *g*_*L*,1_ = *g*_*L*,2_ = 0.1, *G*_*syn*,12_ = *G*_*syn*,21_ = 0.02, *E*_*syn*,12_ = *E*_*syn*,21_ = *−*20. **A**. *A*_*in*_ = 0.1. **B**. *A*_*in*_ = 0.2. **C**. *A*_*in*_ = 1. 22

**Figure 6:**
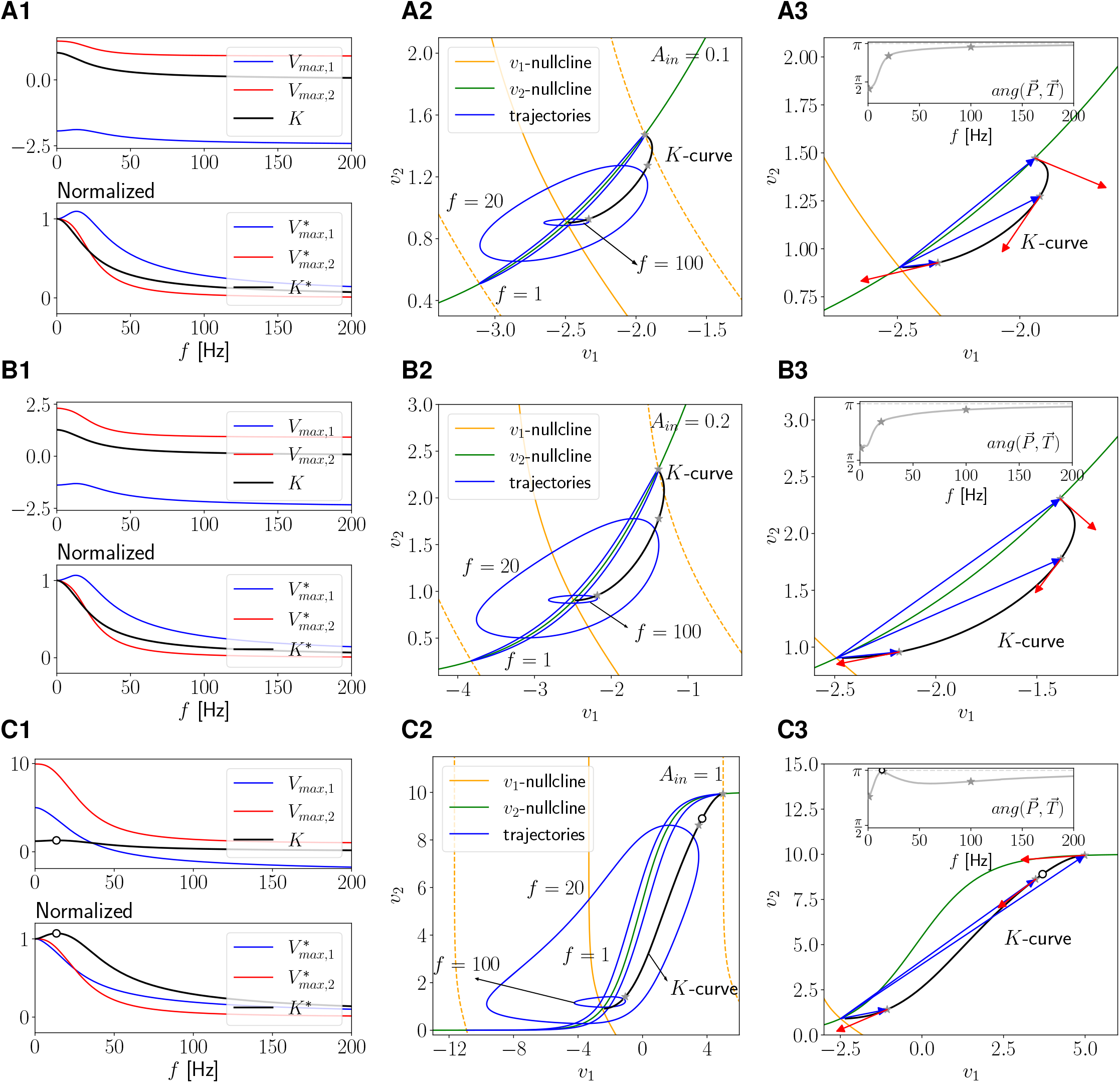
Response of a nonlinear excitatory-inhibitory network of two 1D cells to oscillatory inputs. We used the system (45)-(46) receiving an oscillatory inputs only to cell 1 (*A*_*in*,1_ = *A*_*in*_ and *A*_*in*,2_ = 0). The self-connectivity parameters (if they exist) are considered to be absorbed by the autonomous linear terms (*G*_*syn*,11_ = *G*_*syn*,22_ = 0) and *C*_1_ = *C*_2_ = 1. **Left-Middle-Right columns** as described in Fig. 5. We used the following parameter values: *g*_*L*,1_ = *g*_*L*,2_ = 0.1, *G*_*syn*,12_ = *G*_*syn*,21_ = 0.02, *E*_*syn*,12_ = *−*20, *E*_*syn*,21_ = 60. **A**. *A*_*in*_ = 0.1. **B**. *A*_*in*_ = 0.2. **C**. *A*_*in*_ = 1.

For the excitatory-inhibitory network, cell 1 transitions from being a band-pass filter for low values of *A*_*in*_ (Figs. 6-A1) to a low-pass filter for higher values of *A*_*in*_ (Figs. 6-B1 and C1), while cell 2 is a low-pass filter for all these values of *A*_*in*_. Similarly to the previous example, a peak in *K*(*f*) emerges as the result of a switch in the relative rates at which *V*_*max*,1_(*f*) and *V*_*max*,2_(*f*) decrease (Figs. 6-C1). Note that the presence of a peak in the receiving cell 1, does not cause a resonance in the communication coefficient *K* since the *K*-resonance is present when cell 1 is a low-pass filter.

#### 3.4.3 Phase-plane diagrams and K-curves

Here we extend the method introduced in subsection 3.3 to nonlinear systems in order to investigate the mechanisms of generation of *K*-resonance. The *v*_1_- and *v*_2_-nullclines for the autonomous systems are given by

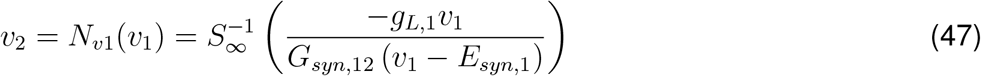

and

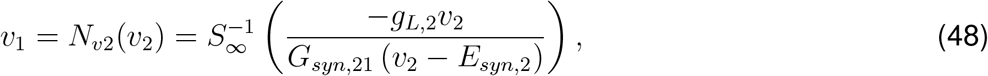

respectively. The corresponding extended (displaced) *v*_1_-nullclines are given by

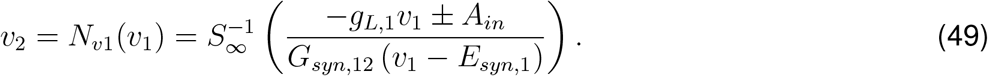

By rescaling time *t ← f t*, system (45)-(46) becomes

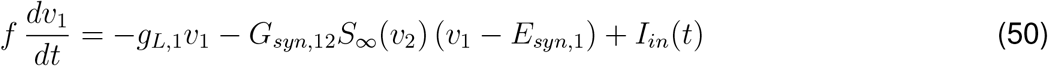

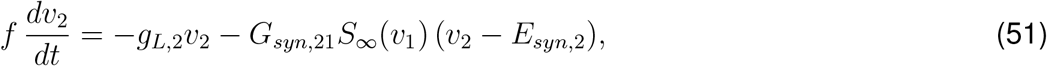

where now *I*_*in*_(*t*) is the sinusoidal input (5) with *f* substituted by 1.

Because both the *v*_1_- and *v*_2_-nullclines are nonlinear, the response limit cycles are affected by the model nonlinearities (Figs. 5 and 6, middle panels). For *f →* 0 (not shown), the response limit cycle trajectory is superimposed to the portion of the *v*_2_-nullcline in between the two extended *v*_1_-nullclines (dashed-red). For small enough values of *f >* 0, the response limit cycle trajectories evolve in a close vicinity of this portion of the *v*_2_-nullcline. For *f →∞*, the response limit cycle reduces to a point (the origin). For large enough values of *f*, the response limit cycles are quasi-elliptic trajectories with a major axis in the direction of the *v*_1_ axis, similarly to the linear case. For intermediate values of *f*, in between these extreme values, the response limit cycles expand, rotate and then shrink as *f* increases. Because of the model nonlinearities, the response limit cycles are not symmetric with respect to the fixed-point and therefore the network impedances and the peak values of *v*_1_ and *v*_2_ on the response limit cycle are different. We use the latter in our definition of the communication coefficient *K* (16).

#### 3.4.4 *K*-resonance: Change of concavity type in the *K*-curves and alignment between the position and tangent vectors

The factors that affect the shape of the *v*_2_-nullcline in between the two extended (nonlinear, dashed-orange) *v*_1_-nullclines, affect the nonlinear properties of the response limit cycle. Because the portion of the *v*_2_-nullcline in between the two extended *v*_1_-nullclines increases with increasing values of *A*_*in*_ (Figs. 5 and 6, from panels A to C), the nonlinear effects on the response limit cycles become more prominent as *A*_*in*_ increases.

For low enough values of *A*_*in*_ (panels A2 in Figs. 5 and 6), while the two systems are nonlinear, the responses are quasi-linear due to the relatively small area in between the two extended *v*_1_-nullclines (compare with panels A2 and B2 in Fig. 3, respectively). As *A*_*in*_ increases (panels C2 in Figs. 5 and 6), the area in between the two extended *v*_1_-nullclines increases and therefore the nonlinearities become more prominent. As this happens, the *K*-curve changes concavity type; It has an initial concave down portion, which switches to a concave-up final portion for higher values of *f*. These changes in concavity type along the *K*-curve allow the position and tangent vectors to be aligned for nonzero input frequency values. The shape of the *K*-curve reflects changes in the rates at which the network peak values *V*_*max*,1_(*f*) and *V*_*max*,2_(*f*) decrease, causing the development of the *K*-resonant peaks as discussed above.

In what follows, we will omit the angle between the position and tangent vectors for the *K*-curve in the figures and will only indicated the *K*-resonance along the curve.

#### 3.4.5 Feedforward and mutual inhibition: Emergence and amplification of *K*-resonance

Feedforward inhibition can generate *K*-resonance when the input amplitude is large enough (Fig. 7-A1 and -A2). Mutual inhibition (Fig. 1-A1) amplifies this *K*-resonance (Fig. 7-B1 and -B2). This amplification is more prominent the larger the difference between the feedforward and feedback connectivity parameters (Fig. 8-A). The presence of *K*-resonance requires the other parameter values (intrinsic properties) to be within some range (Fig. 9), and is absent (or very mild) for small enough values of both *A*_*in*,1_ and for connectivity parameters for which the nonlinear effects are not strong enough. In contrast to the linear case, *K*-resonance is present when cell 2 is a network low-pass filter.

**Figure 7:**
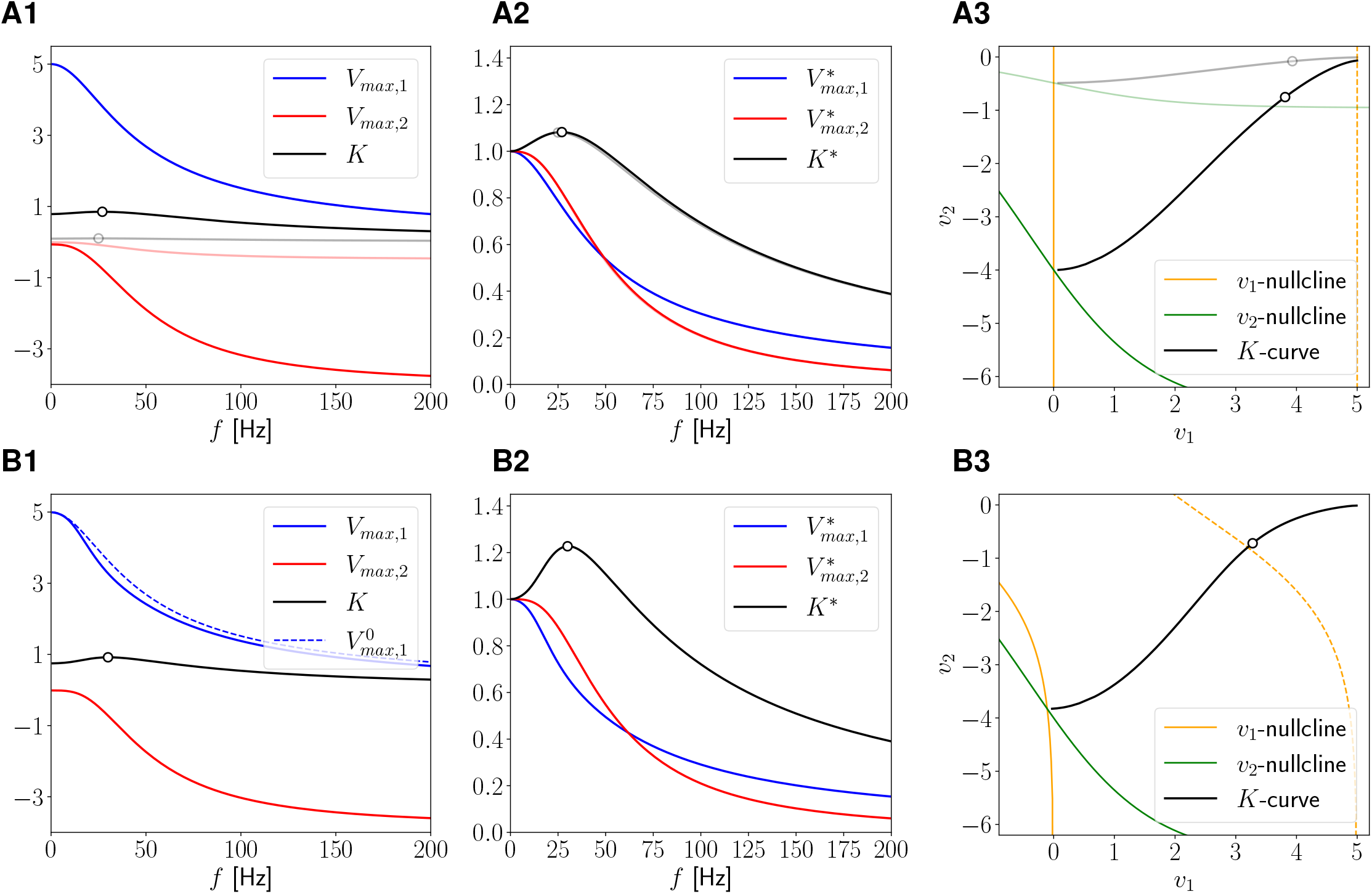
Two-1D-cell network with feedforward and mutual inhibition: *V*_*max*_ and *K*-profiles. We used the system (45)-(46) receiving an oscillatory inputs only to cell 1 (*A*_*in*,1_ = *A*_*in*_ and *A*_*in*,2_ = 0). The self-connectivity parameters (if they exist) are considered to be absorbed by the autonomous linear terms (*G*_*syn*,11_ = *G*_*syn*,22_ = 0) and *C*_1_ = *C*_2_ = 1. **Left column**. Network peak (*V*_*max*,1_ and *V*_*max*,2_) and communication coefficient (*K*) profiles (16). The dots indicate the presence of *K*-resonance. Peak voltage profiles for the disconnected cells (dashed) (achieved by assuming the cells receive an oscillatory input). **Middle column**. Normalized peak profiles and communication coefficient profile. **Right column**. Phase-plane diagram. The solid orange and green lines are the *v*_1_ and *v*_2_ nullclines of the autonomous system, respectively, given by eqs. (47) and (48). The dashed orange lines are the extended *v*_1_-nullclines for the autonomous system for constant inputs equal to *±A*_*in*_, given by eq. (49). The black curve is the *K*-curve it joins the peak values of the response limit cycles in the *v*_1_ and *v*_2_ directions. The dots indicate the presence of *K*-resonance. **A**. Feedforward inhibition: *G*_*in*,12_ = 0 and *G*_*in*,21_ = 0.1 (light colors *G*_*in*,21_ = 0.01). **B**. Mutual inhibition: *G*_*in*,12_ = 0.05and *G*_*in*,21_ = 0.1. We used the following parameter values: *g*_*L*,1_ = *g*_*L*,2_ = 0.2, *E*_*syn*,12_ = *E*_*syn*,21_ = *−*20 and *A*_*in*_ = 1.

**Figure 8:**
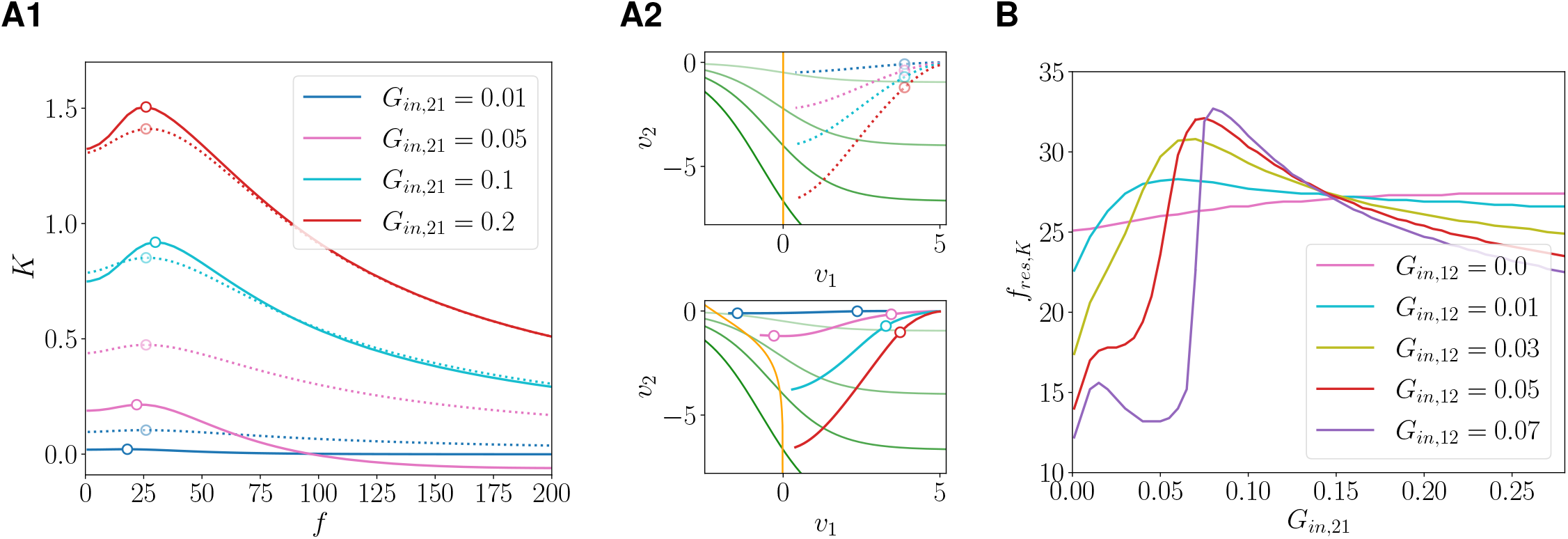
Two-1D-cell network with mutual inhibition: Dependence of the *K*-profiles on the connectivity parameters. The model is given by eqs. (45)-(46) with *A*_*in*_ = 1. **A1**. *K*-profiles for *g*_*L*,1_ = *g*_*L*,2_ = 0.2, *G*_*in*,12_ = 0 (dotted; feedforward inhibition), *G*_*in*,12_ = 0.05 (solid; mutual inhibition) and representative values of *G*_*in*,21_. The dots indicate the *K*-resonances and antiresonances. **A2**. *K*-curves in the phase-plane diagram for the feedforward (top) and mutual (bottom) inhibition cases in A1. The solid orange and green lines are the *v*_1_ and *v*_2_ nullclines of the autonomous system, respectively, given by eqs. (47) and (48). **B**. *K*-resonant frequencies as a function of *G*_*in*,21_, for representative values of *G*_*in*,12_ and *g*_*L*,1_ = *g*_*L*,2_ = 0.2.

**Figure 9:**
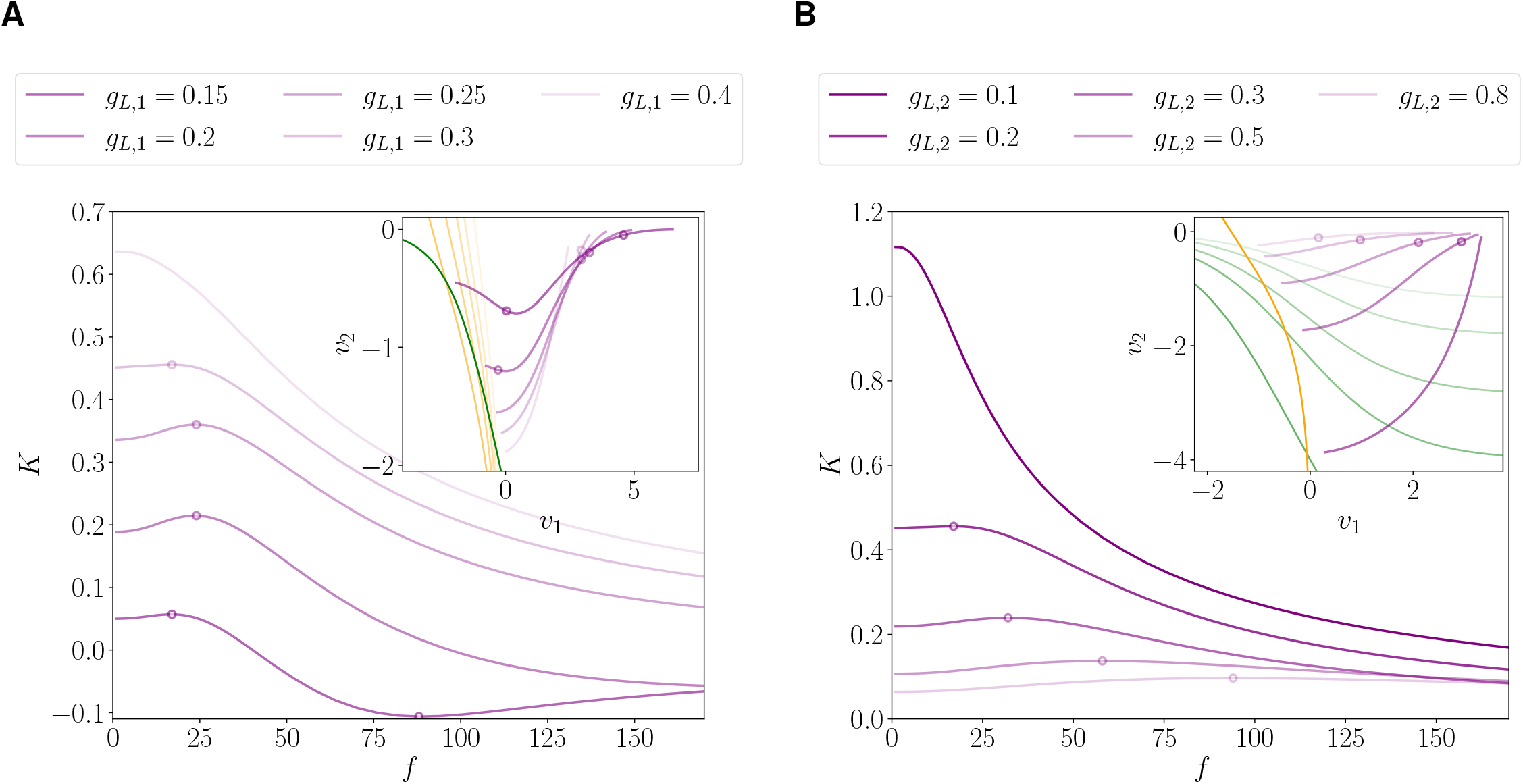
Two-1D-cell network with mutual inhibition: Dependence of the *K*-profiles on the intrinsic model parameters. The model is given by eqs. (45)-(46) with *A*_*in*_ = 1. **A**. *K*-profiles for *g*_*L*,2_ = 0.2, *G*_*in*,12_ = *G*_*in*,21_ = 0.05 and representative values of *g*_*L*,1_. The dots indicate the presence of *K*-resonance and antiresonance. **Inset**. *K*-curves in the phase-plane diagram. The solid orange and green lines are the *v*_1_ and *v*_2_ nullclines of the autonomous system, respectively, given by eqs. (47) and (48). **B**. *K*-profiles for *g*_*L*,1_ = 0.3, *G*_*in*,12_ = *G*_*in*,21_ = 0.05 and representative values of *g*_*L*,2_. The dots indicate the presence of *K*-resonance. **Inset** as described in A.

The mechanism of generation and amplification of *K*-resonance is captured by the change of concavity of the corresponding *K*-curves as *f* increases, which allows the alignment of the position and tangent vectors. This change is more prominent for the mutually inhibitory network (Fig. 7-B3) than for the feedforward network (Fig. 7-A3), and almost imperceptible for weakly feedforward network (Fig. 7-A3, light). The shapes of these *K*-curves reflect the underlying nonlinearities, which are more prominent for the mutually inhibitory than for the feedforward network (compare the phase-plane diagrams in Figs. 7-A3 and -B3).

For feedforward networks, the *K*-profiles and *K*-resonance are amplified by increasing values of the connectivity parameter *G*_*in*,21_ (dotted curves in Fig. 8-A1). As *G*_*in*,21_ increases, the *v*_2_-nullcline displays a stronger nonlinear behaviour in the relevant portion of the phase-plane diagram and the *K*-curves show a more prominent change of concavity (Fig. 8-A2). As a result, the *Q*_*K*_ values increase, the *K*-resonant profiles become sharper and the *K*-resonant frequencies *f*_*res,K*_ slightly increase (purple curve in Fig. 8-B). When the feedback connectivity parameter *G*_*in*,12_ increases (solid curves in Fig. 8-A1) and *G*_*in*,21_ is low enough, the *K*-curves are affected by the nonlinear *v*_1_-nullcline and cover a small range of *v*_2_ values on the phase-plane. The corresponding *K*-profiles are reduced (as compared with the feedforward cases) and can have negative values and antiresonances (see also Fig. 5-C). Additionally, in these cases (low fixed values of *G*_*in*,21_) the *K*-resonant frequencies are non-monotonic when the parameter *G*_*in*,12_ increases (Fig. 8-B). As *G*_*in*,21_ continue to increase, the *K*-curves are more similar to the ones for the feedforward networks and thus the *K*-profiles are amplified, however the *K*-resonance weakens and the frequency *f*_*res,K*_ decreases (Fig. 8-B). By increasing the values of the connectivity parameters further, new network equilibria are created and the analysis presented above ceases to be directly applicable.

The ΔΦ profile generated by the feedforward connectivity (Fig. S2-A) is slightly deformed by the presence of mutual inhibition, as compared to the disconnected network (Fig. S2-B). As expected for inhibitory patterns, cell 2 is delayed as compared to cell 1 and the phase-locking properties are frequency-dependent, transitioning from antiphase and decreasing their phase-difference as *f* increases.

Fig. 9 illustrates the dependence of the *K*-profiles with the intrinsic parameters of the individual cells for a mutually inhibitory network (*G*_*in*,12_ = *G*_*in*,21_). For the lowest values of *g*_*L*,1_ and *g*_*L*,2_ fixed (Fig. 9-A) the network exhibits *K*-resonance and *K*-antiresonance, while for *g*_*L*,1_ fixed and the largest values of *g*_*L*,2_ (Fig. 9-B) the network exhibits *K*-resonance. As *g*_*L*,1_ increases (*g*_*L*,2_ fixed; Fig. 9-A), the *v*_1_-nullcline moves to the right in the phase-plane diagram (Fig. 9-A inset), and the fixed-point shifts toward a region where the nonlinearity for the first variable has less impact on the network’s responses. The *K*-curves cover a smaller range of values of *v*_1_ in the phase-plane diagram as *g*_*L*,1_ increases, and the change of concavity becomes less prominent and then vanishes (e.g., *g*_*L*,1_ = 0.4). As a result, the *K*-profiles raise and are simultaneously attenuated by increasing values of *g*_*L*,1_. The *K*-resonant frequency, first increases and then decreases until it vanishes. As *g*_*L*,2_ increases (*g*_*L*,1_ fixed; Fig. 9-B), the *v*_2_-nullcline raises in the phase-plane diagram (Fig. 9-B inset), giving rise to a change of concavity of the *K*-curve but this curve cover a smaller range of values of *v*_2_. As a result, the *K*-profile is attenuated and the *K*-resonance is generated as *g*_*L*,2_ increases and the *K*-resonant frequency increases.

#### 3.4.6 Feedforward and mutual excitation: Emergence and attenuation of *K*-resonance

Feedforward excitation can also generate *K*-resonance (Fig. 10-A1 and A2). However, in contrast to the mutually inhibitory network, mutual excitation (Fig. 1-A2) attenuates the *K*-profiles (Fig. 10-B1 and B2). Neither feedforward nor mutual excitation produce network resonance (the participating cells remain low-pass filters). In addition, there is no ΔΦ-phasonance (Fig. S3).

**Figure 10:**
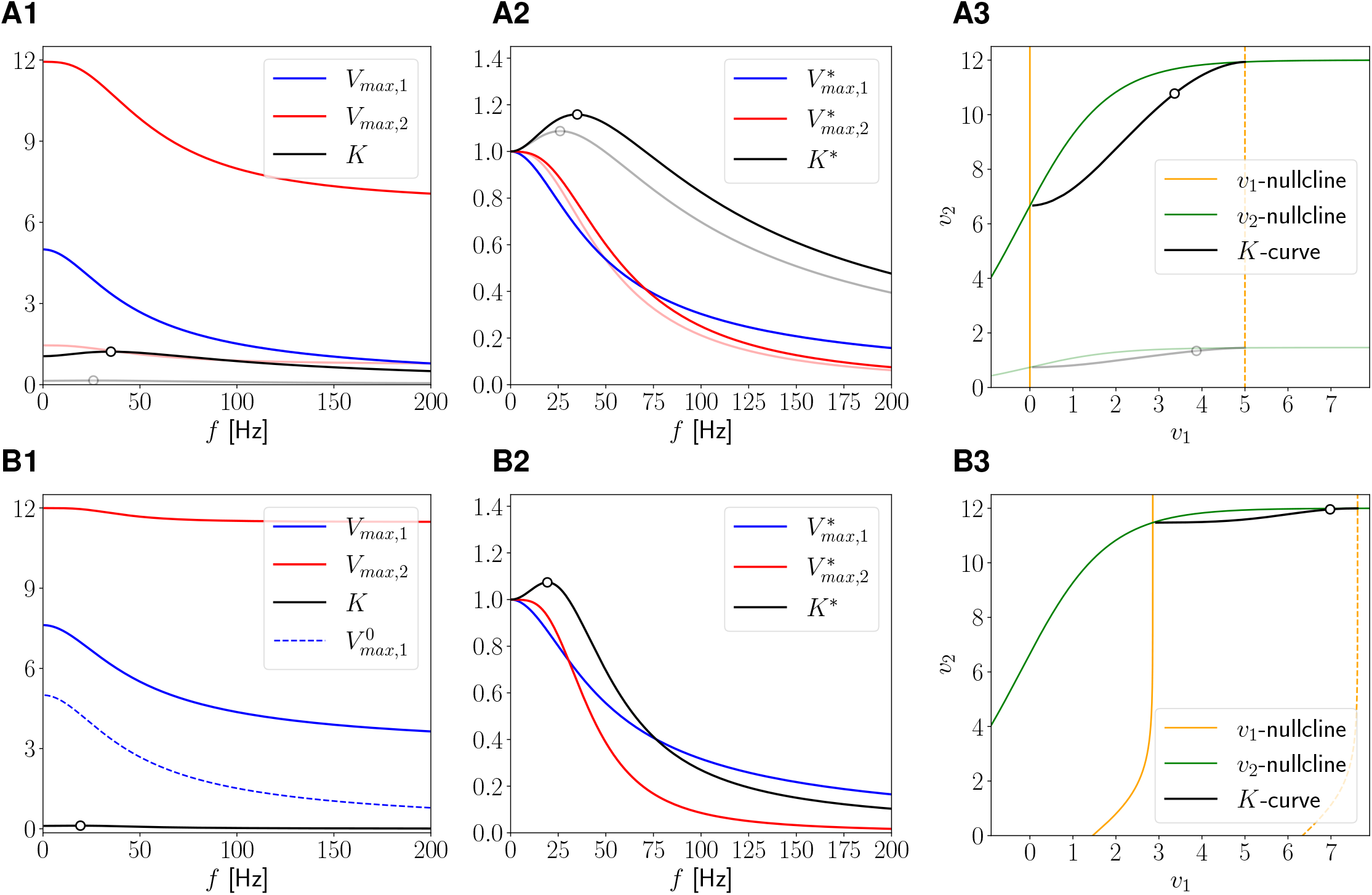
Two-1D-cell network with feedforward and mutual excitation: *V*_*max*_ and *K*-profiles. We used the system (45)-(46) receiving an oscillatory inputs only to cell 1 (*A*_*in*,1_ = *A*_*in*_ and *A*_*in*,2_ = 0). The self-connectivity parameters (if they exist) are considered to be absorbed by the autonomous linear terms (*G*_*syn*,11_ = *G*_*syn*,22_ = 0) and *C*_1_ = *C*_2_ = 1. **Left column**. Network peak (*V*_*max*,1_ and *V*_*max*,2_) and communication coefficient (*K*) profiles (16). The dots indicate the presence of *K*-resonance. Peak voltage profiles for the disconnected cells (dashed) (achieved by assuming the cells receive an oscillatory input). **Middle column**. Normalized peak profiles and communication coefficient profile. **Right column**. Phase-plane diagram. The solid orange and green lines are the *v*_1_ and *v*_2_ nullclines of the autonomous system, respectively, given by eqs. (47) and (48). The dashed orange lines are the extended *v*_1_-nullclines for the autonomous system for constant inputs equal to *±A*_*in*_, given by eq. (49). The black curve is the *K*-curve it joins the peak values of the response limit cycles in the *v*_1_ and *v*_2_ directions. The dots indicate the presence of *K*-resonance. **A**. Feedforward excitation: *G*_*ex*,12_ = 0 and *G*_*ex*,21_ = 0.05 (light colors *G*_*ex*,21_ = 0.005). **B**. Mutual excitation: *G*_*ex*,12_ = 0.01 and *G*_*ex*,21_ = 0.05. We used the following parameter values: *g*_*L*,1_ = *g*_*L*,2_ = 0.2, *E*_*syn*,12_ = *E*_*syn*,21_ = 60 and *A*_*in*_ = 1.

The mechanism of generation of *K*-resonance in the feedforward excitatory networks is similar to the mechanism described for feedforward inhibitory networks. In the phase-plane diagram, the *K*-curve shifts from being concave down to concave up as *f* increases (Fig. 10-A3), allowing the alignment of the position and tangent vectors, and therefore causing the emergence of *K*-resonance. As the connection parameter *G*_*ex*,21_ increases the concave up portion of the *K*-curve increases, causing an amplification of the *K*-resonance. In contrast, increasing values of *G*_*ex*,12_ causes the *v*_1_-nullcline to move to a region in the phase-plane diagram where the nonlinearities are less prominent (Fig. 10-B3), thus causing the attenuation of the *K*-resonance, which vanishes as *G*_*ex*,12_ continues to increase. The *K*-profiles raise by increasing values of *g*_*L*,1_ or decreasing values of *g*_*L*,2_, but they quickly loose their preferred frequency (not shown).

#### 3.4.7 Inhibitory-excitatory networks: Emergence of resonance in cell 2 and attenuation of the *K*-profiles

In these networks, cell 1 inhibits cell 2 (*G*_*in*,21_ *>* 0), which in turn excites cell 1 (*G*_*ex*,12_ *>* 0) (Fig. 1-A3). In the absence of excitation (the feedforward inhibitory network discussed above), network resonance is not present in either cell and the *K*-profile can exhibit *K*-resonance. If the input amplitude *A*_*in*_ is large enough, these networks generate resonances in cell 2 and reduce the values of the communication coefficient *K* for small input frequencies (Fig. 11-A1). In addition, these networks show antiphase synchronization between the two cells and ΔΦ is negative and non-monotonic (Fig. S4), in contrast to the previous cases discussed above, which are all monotonic.

**Figure 11:**
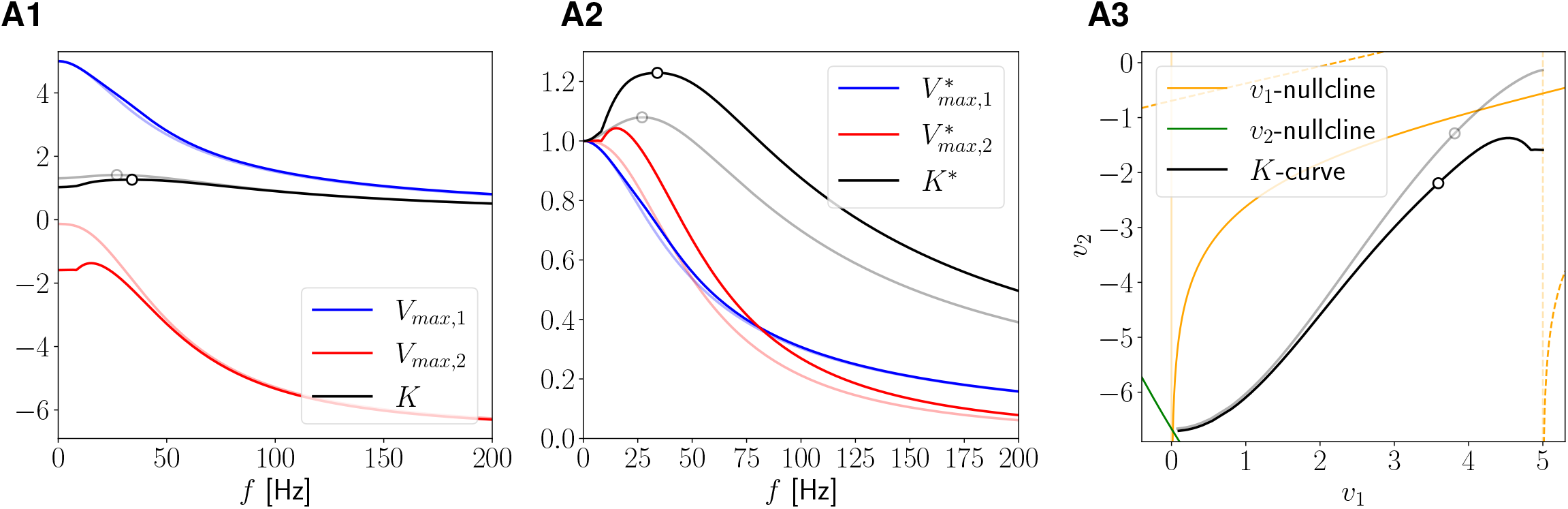
Two-1D-cell inhibitory-excitatory network. (connectivity parameters: *G*_*in*,21_ and *G*_*ex*,12_). **Left-Middle-Right columns** as descripted in Fig. 10. We used equations (45)-(46) with *A*_*in*_ = 1 and the following additional parameter values: *g*_*L*,1_ = *g*_*L*,2_ = 0.2, *G*_*in*,21_ = 0.2, *G*_*ex*,12_ = 0.05 (light colors feedforward inhibition, i.e., *G*_*ex*,12_ = 0), *E*_*syn*,21_ = *−*20 and *E*_*syn*,12_ = 60.

As discussed above, feedforward inhibition can generate *K*-resonances. In the phase-plane diagram, the *K*-curve is concave down for small values of *f*, and then shift to concave up (Fig. 11-A3, gray curve) as *f* increases. When the feedback excitation is active (*G*_*ex*,12_ *>* 0), the *K*-curve covers a smaller range of values of *v*_2_. In particular, the values of *K* for small input frequencies are smaller than in the feedforward inhibitory networks. Thus, the excitatory feedback attenuates the *K*-resonance if it exists, or generates *K*-resonance but with smaller *K* values than in the feedforward cases.

Fig. 12 illustrates how the balance of excitation, inhibition and the intrinsic properties of the participating nodes affect the resonant properties of the *K*-profiles. *K*-resonance are more prominent the larger *G*_*in*,21_, which is inherited from the feedforward inhibitory network (compare the solid and dashed curves for *G*_*in*,21_ = 0.4 and *G*_*in*,21_ = 0.1, respectively in Fig. 12-A). In contrast, the *K*-profiles for the lower levels of *G*_*in*,21_ are more sensitive to increasing values of *G*_*ex*,12_, which cause an attenuation of these phenomena.

**Figure 12:**
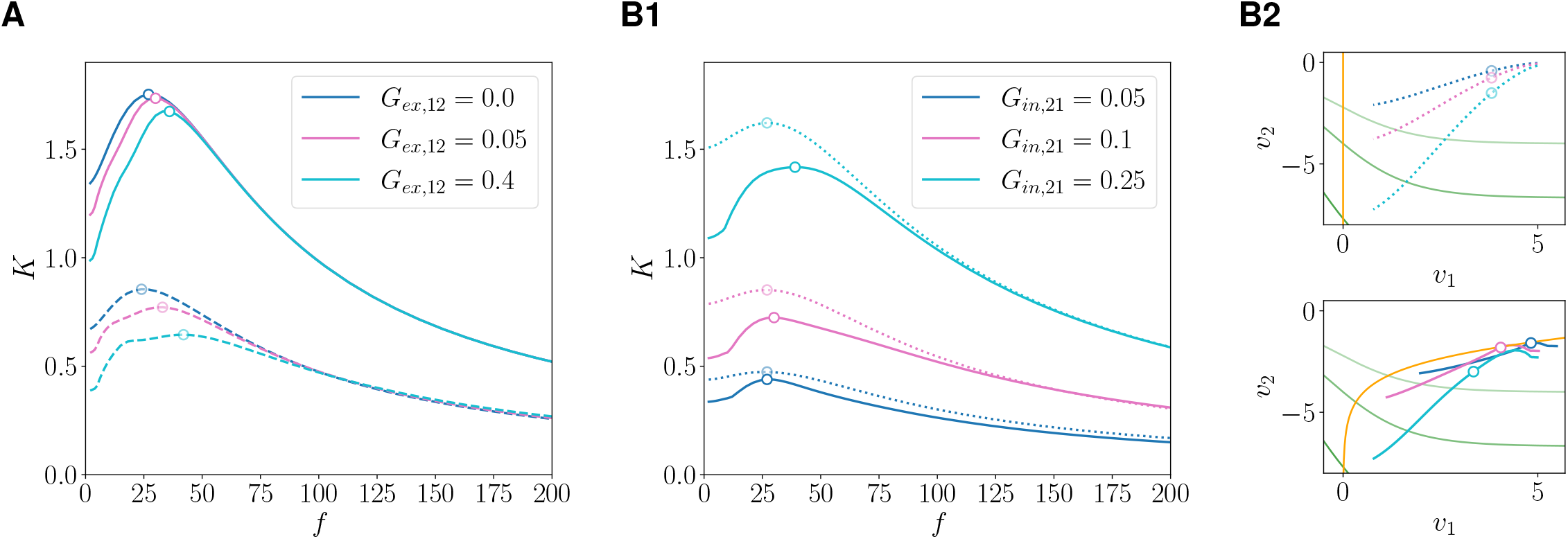
Two-1D-cell inhibitory-excitatory network: Dependence of the K -profiles on the connectivity parameters. We used eqs. (45)-(46) with *A*_*in*_ = 1, *E*_*syn*,21_ = *−*20 and *E*_*syn*,12_ = 60. **A**. *K*-profiles for *g*_*L*,1_ = *g*_*L*,2_ = 0.1, *G*_*in*,21_ = 0.1 (dashed) and *G*_*in*,21_ = 0.4 (solid). The dots indicate the presence of *K*-resonance. **B1**. *K*-profiles for *g*_*L*,1_ = *g*_*L*,2_ = 0.2, *G*_*ex*,12_ = 0 (dotted) and *G*_*ex*,12_ = 0.1 (solid). The dots indicate the presence of *K*-resonance. **B2**. *K*-curves in the phase-plane diagram for the feedforward inhibition (top) and feedback excitation (bottom) cases in B1. The solid orange and green lines are the *v*_1_ and *v*_2_ nullclines of the autonomous system, respectively, given by eqs. (47) and (48).

Increasing values of *G*_*in*,21_, raise the *K*-profiles and cause an amplification of the *K*-resonance (Fig. 12-B1, solid). While the *K*-profiles are larger the larger *g*_*L*,1_ and *g*_*L*,2_ (compare panels A and B1), the *K*-resonance is more prominent the smaller *g*_*L*,1_ and *g*_*L*,2_. All this is inherited from the feedforward inhibitory network (dotted) that we already analyze with help of the *K*-curves (Fig. 12-B2, top). When the feedback exitation is active, the *v*_1_-nullcline (orange in Fig. 12-B2, bottom) displaced the *K*-curves in the phase-plane and they cover a reduced range of *v*_2_ values. This change reduce the *K* values for small input frequencies and thus the *K*-resonance becomes more prominent than in the inhibitory feedforward networks.

We note that if *A*_*in*_ is increased further in this type of network, the network displays mixed-mode oscillations (MMOs) and the responses become distorted. The analysis of this type of patterns is beyond the scope of this work.

#### 3.4.8 Excitatory-inhibitory networks: Amplification of *K*-profiles and attenuation of *K*-resonance

In this type of networks, cell 1 excites cell 2 (*G*_*ex*,21_ *>* 0), which in turn inhibits cell 1 (*G*_*in*,12_ *>* 0) (Fig. 1-A4). With this architecture, both cells exhibit network resonance and the network exhibits *K*-resonance, but they are relatively mild (Fig. 13-A1 and A2). In the phase-plane diagram (Fig. 13-A3), the presence of feedback inhibition causes the *v*_1_-nullcline to shift, moving the equilibrium toward values where cell 2 approaches the inactivation zone. This causes the network impedance and *K*-profiles to be less prominent than for the feedforward excitatory network discussed above. Note that in contrast to the other network architectures, cell 1 exhibits phasonance (Fig. S5), however, ΔΦ is always positive and monotonically increasing.

**Figure 13:**
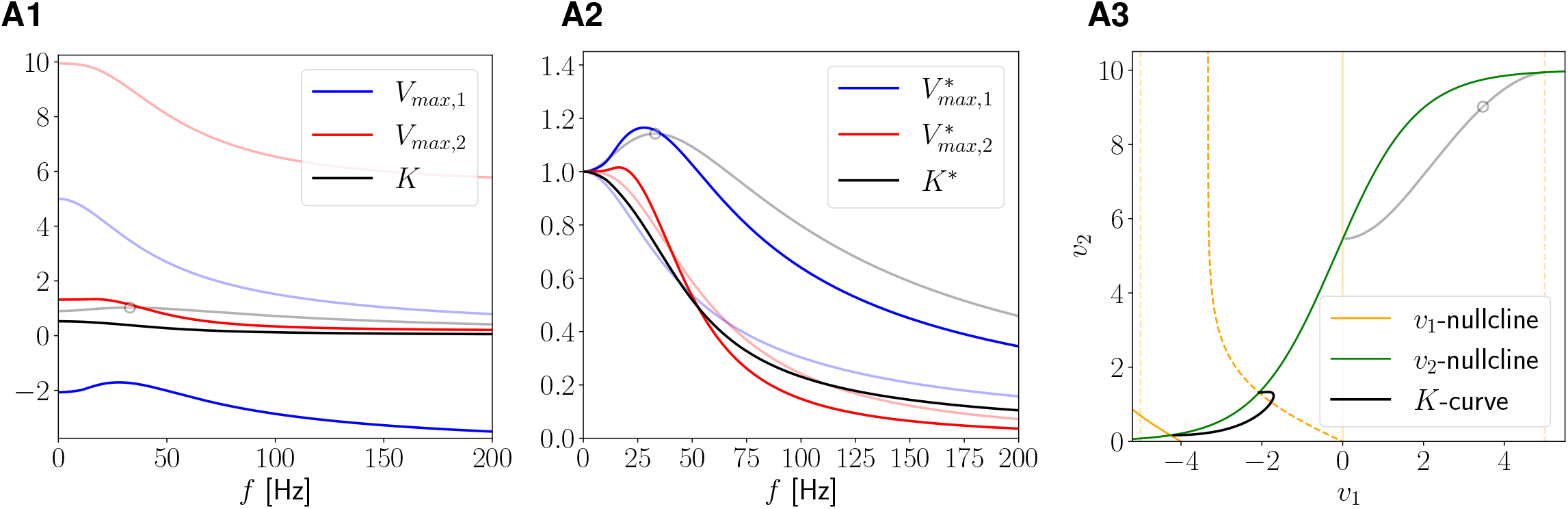
Two-1D-cell excitatory-inhibitory network. (connectivity parameters: *G*_*ex*,21_ and *G*_*in*,12_). **Left-Middle-Right columns** as descripted in Fig. 10. We used equations (45)-(46) with the following additional parameter values: *g*_*L*,1_ = *g*_*L*,2_ = 0.2, *G*_*ex*,21_ = 0.04, *G*_*in*,12_ = 0.1 (light colors feedforward excitation, i.e., *G*_*in*,12_ = 0), *E*_*syn*,21_ = 60 and *E*_*syn*,12_ = *−*20.

### 3.5 Two-2D-cell networks with nonlinear synaptic connectivity

In Section 3.4 we investigated in detail the mechanisms of generation of preferred communication frequencies in two-cell networks of passive cells where the individual nodes are low-pass filters. In these networks, *K*-resonance and ΔΦ-phasonance requires the presence of nonlinear connectivity and are absent when the connectivity is linear (see Section 3.1).

Here, we extend our investigation to two-cell networks where at least one of the cells is a resonator (2D). From equations (1)-(2) the dynamics of these networks are given by

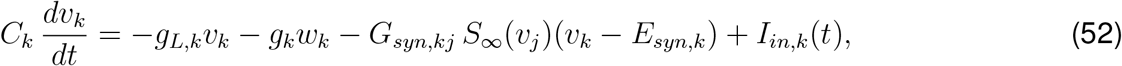

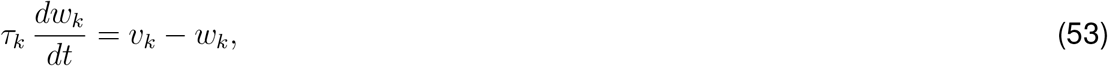

for *k* = 1, 2. As in the previous section, only cell 1 receives a sinusoidal input, thus, *I*_*in*,1_(*t*) is given by (5) and *I*_*in*,2_(*t*) = 0.

From our discussion in Section 3.1, eqs. (25)-(26), the presence of resonance and phasonance in the isolated cell 2, but no in the isolated cell 1 is communicated to *K*(*f*) and ΔΦ(*f*) in the corresponding linear networks. A linear network consisting of a resonator (RN) and passive (PN) nodes exhibits *K*-resonance if the input arrives in the PN, but not if it arrives in the RN. A similar result holds for ΔΦ-phasonance provided the RN also exhibits phasonance.

Below, we analyze the effects of the nonlinearities on the frequency-dependent properties of *K*(*f*) and ΔΦ(*f*) by focusing on representative examples. We restrict our analysis to the parameter regimes for which the autonomous (unperturbed) network has a single stable equilibrium. The investigation of more complex intrinsic network scenarios (e.g., intrinsic bistability and oscillations [43, 50]) requires further development of the dynamical systems tools, and is beyond the scope of the present work.

#### 3.5.1 Mutually inhibitory resonant 2D- and passive 1D-cell: Generation of network resonances, *K*-resonances and antiresonances

We consider mutually inhibitory networks consisting of one RN (resonator, 2D, resonant frequency 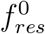) and one PN (passive, 1D). We chose the parameter values for the individual cells so that their peak impedances coincide when they are isolated. We divide our study in two cases according to which cell type receives the oscillatory input: PN (Case A, Fig. 1-B1) and RN (Case B, Fig. 1-B2). We show representative examples in Figure 14.

**Figure 14:**
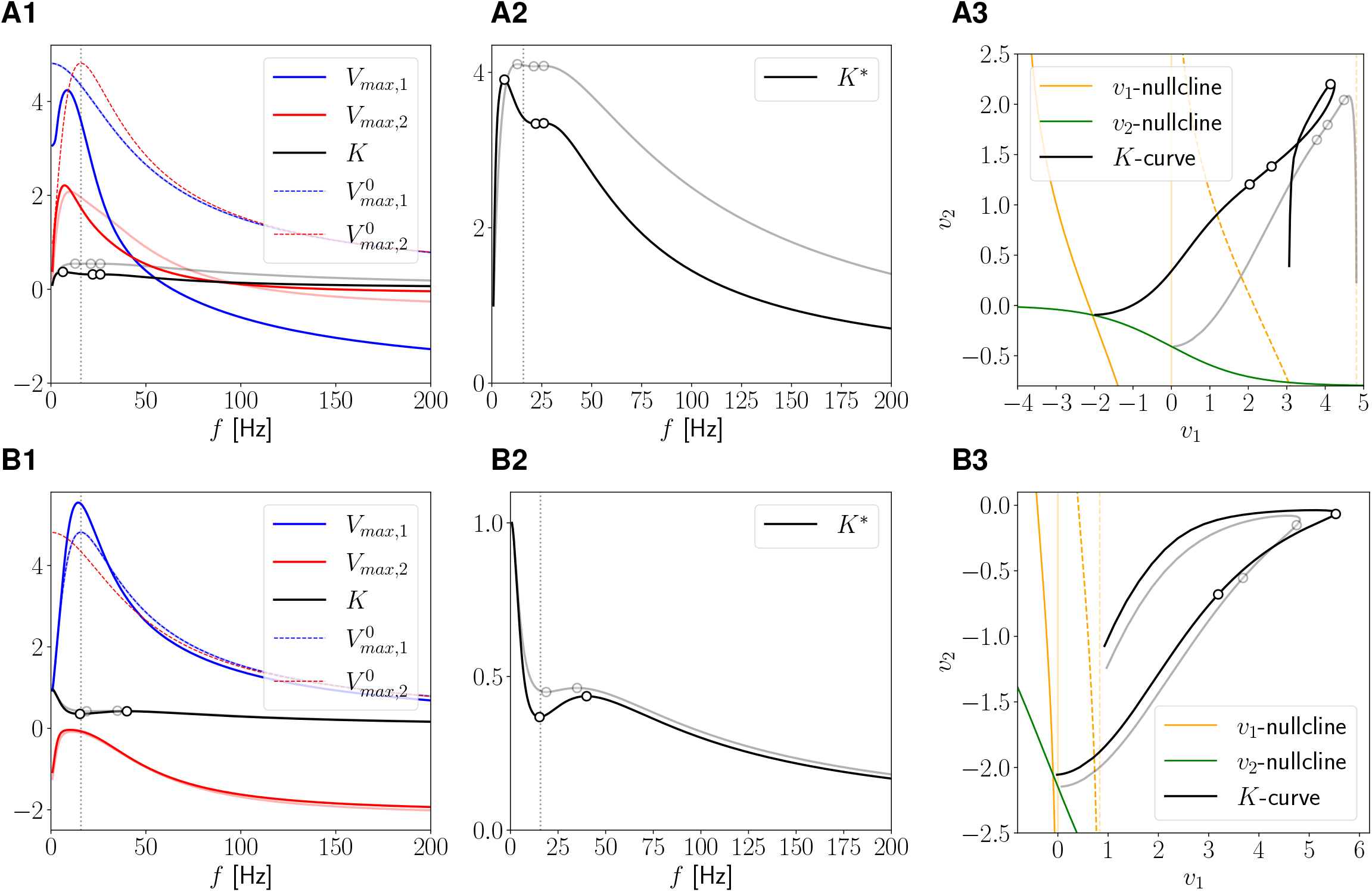
Two-cell network: Mutually inhibitory resonant (RN) and passive (PN) nodes. We used eqs. (52)-(53) with *g*_*L*_ = 0.2, *g* = 1 and *τ* = 120 (RN), *g*_*L*_ = 0.2078, and *g* = 0 (PN), and *G*_*in*,21_ = *G*_*in*,12_ = 0.05. The resonant frequency for the isolated RN is 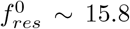 (vertical dotted). The network natural frequency is *f*_*ntwk,nat*_ *∼* 10.35. **Left column**. Network peak (*V*_*max*,1_ and *V*_*max*,2_) and communication coefficient (*K*) profiles (16). The dots indicate the presence of *K*-resonance. Peak voltage profiles for the disconnected cells (dashed) (achieved by assuming the cells receive an oscillatory input). For comparison, we consider the feedforward inhibitory network, with *G*_*in*,12_ = 0 (light blue, light red and gray). **Middle column** Normalized communication coefficient profile (gray for feedforward inhibitory network). **Right column**. Phase-plane diagram. The solid orange and green lines are projections of the *v*_1_ and *v*_2_-nullcline of the autonomous system. The dashed orange lines are the extended *v*_1_-nullclines for the autonomous system for constant inputs *±A*_*in*_. The black curve is the *K*-curve it joins the peak values of the response limit cycles in the *v*_1_ and *v*_2_ directions. The dots indicate *K*-resonances and antiresonances. Light colors for feedforward inhibitory network. **A**. Cell 1 is the PN and cell 2 is the RN. **B**. Cell 1 is the RN and cell 2 is the PN.

In both cases we observe network resonances with resonant frequencies smaller than 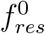 (Fig. 14-A1 and B1), but there are disparities in the qualitative properties of the *K*-profiles and ΔΦ profiles. This is primarily due to the differences in the the filtering propertiess of the cell receiving the input, which are more prominent at low input frequencies, and different in both cases.

In Case A (cell 1 is a PN and cell 2 is a RN; Fig. 14-A1), the feedforward inhibition attenuates the *V*_*max*,2_ profile (light red) as compared to the one for the isolated PN, and causes a decrease in the resonant frequency of cell 2 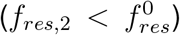. The addition of feedback inhibition induces a resonance in cell 1 with 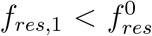 and attenuates the *V*_*max*,1_ profile. In Case B (cell 1 is a RN and cell 2 is a PN; Fig. 14-B1), the feedforward inhibition induces a resonance in cell 2, while the addition of feedback inhibition causes a small decrease in the resonant frequency of cell 1 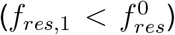 and amplifies the *V*_*max*,1_ profiles at frequencies near its resonance frequency.

The *K*-curves for Cases A (Fig. 14-A3) and B (Fig. 14-B3) have a qualitatively different structure. Specifically, the *K*-curve for Case A with feedforward inhibition decreases very slowly along variable *v*_1_ and rapidly along variable *v*_2_ (gray curve in Fig. 14-A3). After reaching its maximum *v*_2_ value, the *K*-curve switches concavity several times as *f* continuous to increase, thus meeting the conditions for the alignment between the tangent and position vectors at several frequency values, before approaching the fixed point as *f → ∞*. When the feedback inhibition is active, the *v*_1_-nullcline shifts to the left in the phase plane, causing the *K*-curve to start at lower *v*_1_ values (compare black and gray curves in Fig. 14-A3). The *K*-curve then increases along both variables as *f* increases and turnes around, which is associated with resonance in both nodes. We observe a self-intersection in the *K*-curve followed by two changes in concavity. Again, in this case, the conditions are met for the tangent and position vectors to be aligned at several frequency values.

In Case B, the *v*_1_-nullcline is only slightly modified when comparing the network with feedforward and mutual inhibition (Fig. 14-B3). Consequently, the corresponding *K*-curves are very similar, showing two points of alignment of the tangent and position vectors, the first of which occurs very close to the region where variable *v*_1_ reaches its maximum value.

When comparing the nullclines for the two cases, we observe that the extended *v*_1_-nullclines (for constant inputs *± A*_*in*_) deviate further from the *v*_1_-nullcline in Case A than in Case B. This leads to lower *K* values for input frequencies near zero for Case A as compared to networks for Case B. In all cases, we observe that for large values of the input frequency, the *K*-curves move through the phase-plane in a manner similar to the curves analyzed for networks of two PNs, reaching the fixed point with upward concavity. For the mutual inhibitory networks, the position of the initial point of the *K*-curves in the phase-plane (with respect to the final part of the curve) determines their behavior at low frequency values and the existence or absence of self-intersections.

The observations about the *K*-curves explain the qualitative differences of the *K*-profiles between the two Cases (Fig. 14-A2 and -B2). In both cases, in the presence of feedback inhibition (Fig. 14-A2 and -B2, black), the *K*-profile exhibits a peak (*K*-resonance) and a trough (*K*-antiresonance) at frequencies near the resonant frequency for the isolated cell 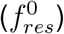. However, the *K*-profile in Case A shows a second resonance near the antiresonance and almost at the same value, and the resonant more prominent resonant frequency in Case A is smaller than the antiresonance frequency (Fig. 14-A2), while the resonant frequency in Case B is larger than the antiresonance frequency (Fig. 14-B2).

In both cases, feedback inhibition attenuates the *K*-curves as compared to feedforward inhibition (Fig. 14-A2 and -B2, gray) and amplifies the differences between the resonances and antiresonances. The biggest differences of the *K*-curves between feedback (black) and feedforward inhibition (gray) occur in a frequency band around the resonant frequency of the RN 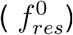. By increasing the values of 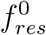, *K* could become a low pass filter (not shown).

We have also observed disparities between the phase profiles for the two cases (Fig. S6). In Case A, cell 1 shows a phasonance at small frequencies and cell 2 shows an antiphase synchronization (Φ_*ntwk*,2_ = *−π*) at frequencies near 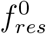 (Fig. S6-A). In Case B, cell 1 shows a phasonance at frequencies near 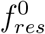 and cell 2 shows an antiphase synchronization at frequencies near the network natural frequency *f*_*ntwk,nat*_ (Fig. S6-B). Finally, in both cases, the ΔΦ-profiles show antiphase synchronization (ΔΦ = *−π*) at frequencies that are not directly related with 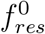 or the network natural frequency.

#### 3.5.2 Resonant 2D-node connected to passive 1D-node with inhibition-excitation: Attenuation of network resonances

We consider network architectures where cell 1 inhibits cell 2, which in turn excites cell 1. We present our results in Fig. 15. We already analyzed this type of architecture in subsection 3.4.7 for networks of two passive cells. We chose the same parameter values for the individual cells as in Section 3.5.1 (Fig. 14), so their peak impedances coincide when they are isolated (Figs. 14 and 15, light colors). We divide our study as in the same two cases as in Section 3.5.1 (see Fig. 1-B3 and B4, respectively).

**Figure 15:**
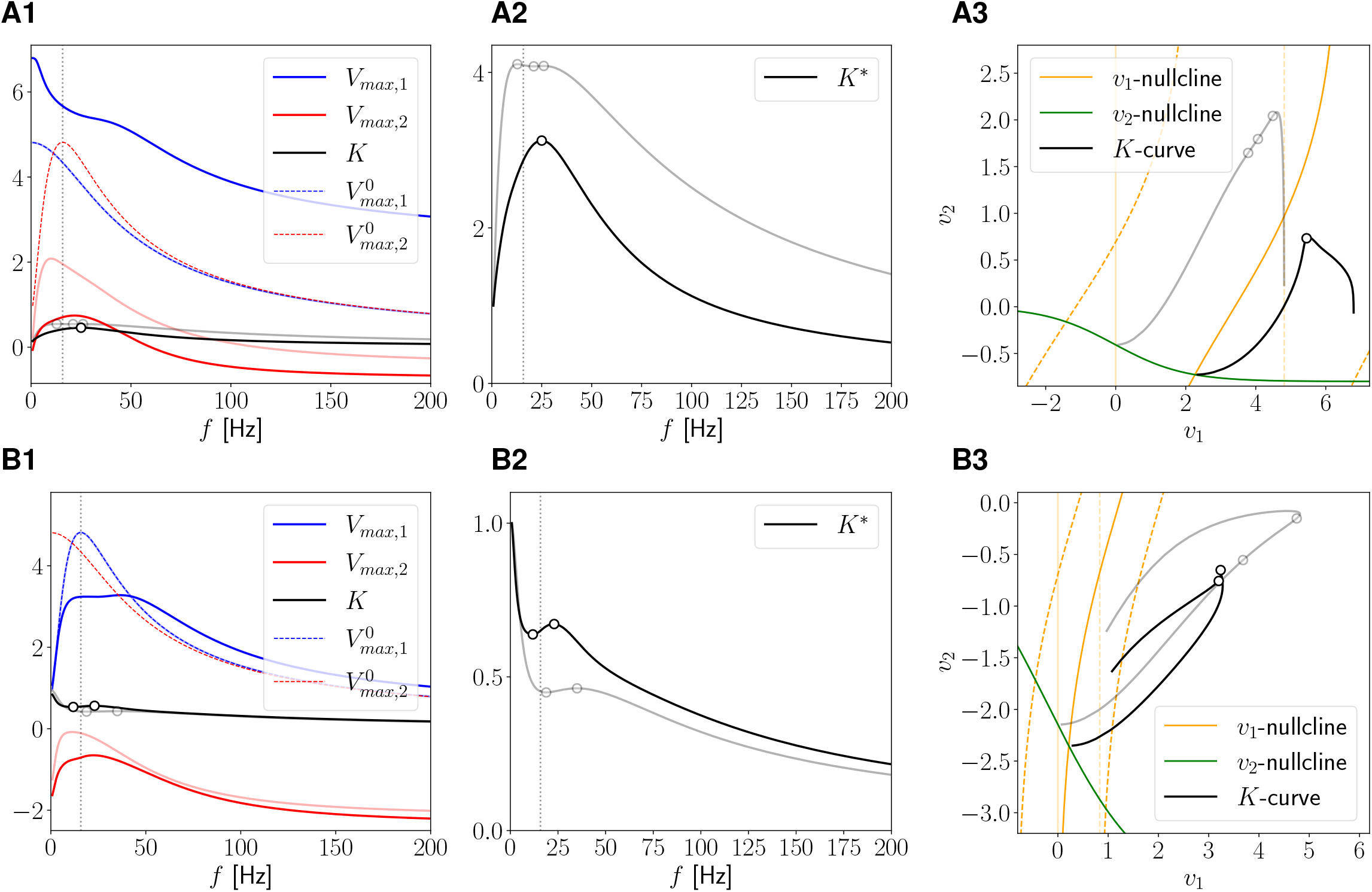
Inhibitory-excitatory network: Resonant (RN) and passive (PN) nodes. We used eqs. (52)-(53) with *g*_*L*_ = 0.2, *g* = 1 and *τ* = 120 (RN), *g*_*L*_ = 0.2078, *g* = 0 (PN). The resonance frequency for the isolated cell is 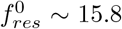 (vertical dotted). **Left-Middle-Right columns** as descripted in Fig. 14. **A**. Cell 1 is the PN and cell 2 is the RN. The connection parameters are *G*_*in*,21_ = 0.05 and *G*_*ex*,12_ = 0.025. The network natural frequency is *f*_*ntwk,nat*_ *∼* 28.28. **B**. Cell 1 is the RN and cell 2 is the PN. The connection parameters are *G*_*in*,21_ = 0.05 and *G*_*ex*,12_ = 0.05. The network natural frequency is *f*_*ntwk,nat*_ *∼* 40.05.

In Case A (cell 1 is a PN; Fig. 15-A) feedback excitation leads to an increase in the *V*_*max*,1_ profile and a decrease in *V*_*max*,2_ profile, along with an increase in the resonance frequency (*f*_*res*,2_) as compared to the feedforward network.

The *v*_1_-nullcline shifts to the right in the phase-space diagram (Fig. 15-A3), causing the *K*-curve to shift accordingly. This results in the above mentioned changes in the voltage peak profiles.

The *K* profile for the feedforward inhibitory network shows two resonances and one antiresonance in between at frequencies close to the resonant frequency 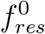 for the isolated RN (Fig. 15-A2, gray). As the result of feedback excitation, the *K*-profile is attenuated (Fig. 15-A2, black), the number of resonances decreases to one, and the antiresonance vanishes. This attenuation effect is similar to that observed for the analogous networks of passive cells (Section 3.4.7).

In Case B (cell 1 is a RN; Fig. 15-B), feedback excitation causes an attenuation of the network resonances and generates antiresonance and a new resonance in the response profiles for cell 1 (Fig. 15-B1, blue). Because the (resonant) peaks and (antiresonant) trough of the *V*_*max*,1_ profile are significantly close to each other, the effect of feedback excitation can be interpreted as broadening the resonant frequency band (around the resonant frequency 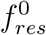 for the isolated RN) as the *V*_*max*,1_ profile is attenuated. Feedback excitation also attenuates the *V*_*max*,2_ profile and increases the resonant frequency (Fig. 15-B1, red), but in contrast to the *V*_*max*,1_ profile, there is no significant change in the resonant frequency band.

The *K*-curve cover a narrower range of values in both variables (Fig. 15-B3), and it shows two frequencies at which the tangent and position vectors align, beyond which the curve exhibits no further changes in curvature.

The corresponding *K*-profile is amplified by feedback excitation (Fig. 15-B2) and the resonance and antiresonance frequencies created by feedback inhibition Fig. 15-B2, gray) become closer to each other and decrease in the presence of feedback excitation Fig. 15-B2, black).

Regarding the phases, we observe that cell 1 shows a phasonance in networks where the isolated cell 1 is resonant (Fig. S7-B), while cell 2 shows antiphase synchronization in both cases (Fig. S7-A and B). Additionally, in both cases the phase difference shows antiphase synchronization (ΔΦ = *− π*) in frequencies near the *K*-resonance and *K*-antiresonance frequencies.

#### 3.5.3 Two resonant nodes with mutual inhibition

Here, we extend our results in Section 3.5.1 to include two resonators in the network (Fig. 1-C). Specifically, we consider two mutually inhibitory resonators (RN1 and RN2) with resonant frequencies 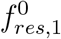 and 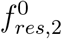 (when RN1 and RN2 are isolated). As above, the oscillatory input arrives only at cell 1. The parameter values for RN1 and RN2 were chosen so that their (isolated) impedances coincide, while they have different resonant frequencies. We divide our study in two cases according to the relative value of their resonant frequencies: 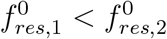 (Case A) and 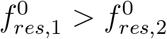 (Case B).

In both cases, the response of each cell in the network is similar to the corresponding case examined in Section 3.5.1. More specifically, both node show network resonance at frequencies equal to or between the resonant frequencies of the isolated cells (Fig. 16). The resonant voltage peak profiles in Case A (Fig. 16-A) are slightly sharper than in Case B (Fig. 16-B). The voltage peak profile for cell 1 is higher the larger its (isolated) resonant frequency (compare Figs. 16-A and -B). The voltage peak profile for cell 2 is also higher at its largest (isolated) resonant frequency, but the corresponding resonant frequencies are closer to the smallest (isolated) resonant frequency.

**Figure 16:**
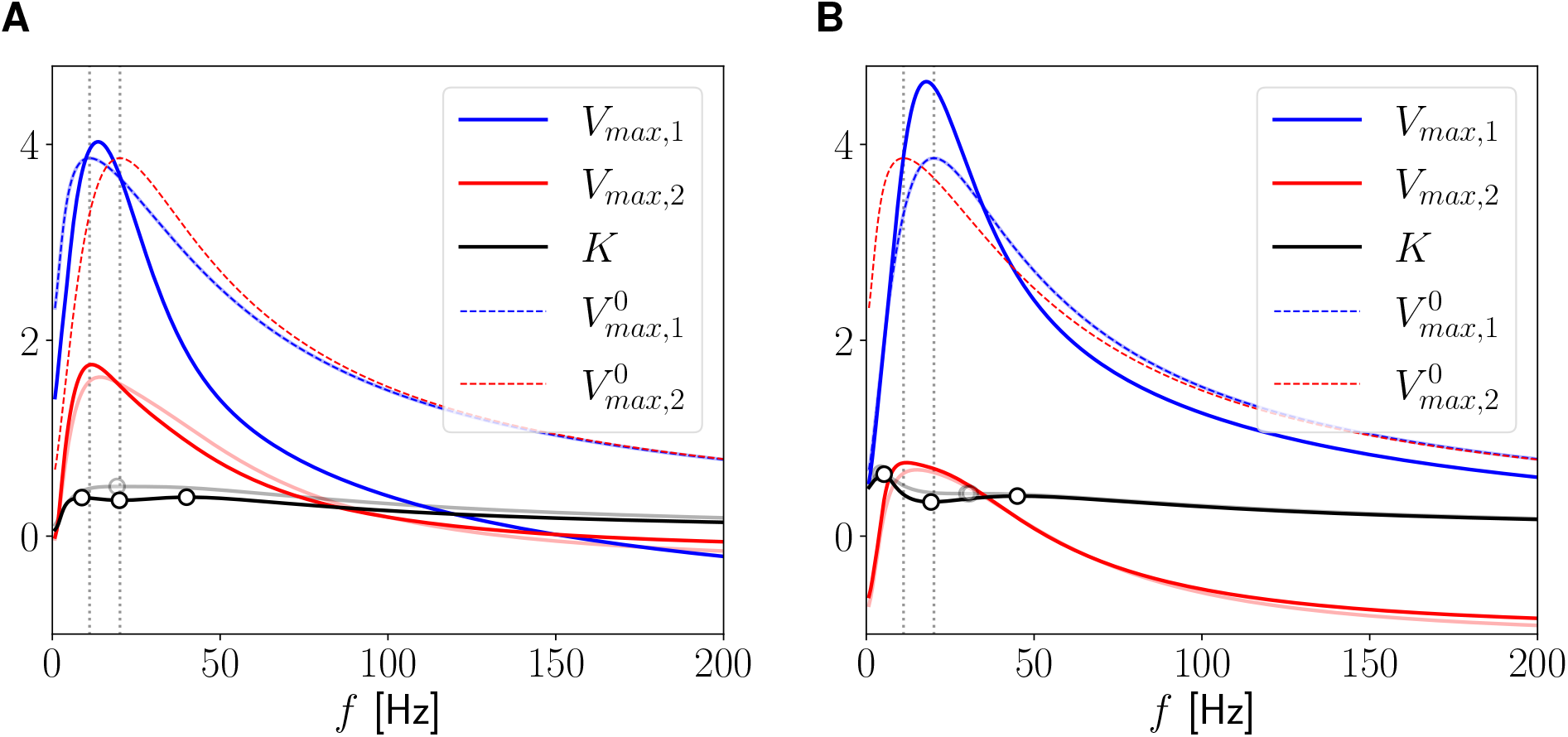
Two mutually inhibitory resonant 2D nodes (RN1 and RN2). We used eqs. (52)-(53) with *g* = 0.2 *τ* = 75.63 (RN1) and *g* = 1.4 and *τ* = 101.58 (RN2). The resonant frequencies of the isolated resonant nodes (vertical-dashed) are 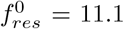 (RN1) and 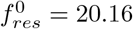 (RN2). **A, B**. Network peak (*V*_*max*,1_ and *V*_*max*,2_) and communication coefficient (*K*) profiles (16). The dots indicate the presence of *K*-resonance and antiresonance. Peak voltage profiles for the disconnected cells (dashed) (achieved by assuming the cells receive an oscillatory input). For comparison, we consider the feedforward inhibitory network, with *G*_*in*,12_ = 0 (light blue, light red and gray). **A**. Cell 1 is RN1 and cell 2 is RN2. **B**. Cell 1 is RN2 and cell 2 is RN1. We used the following additional parameter values: *g*_*L*,1_ = *g*_*L*,2_ = 0.25 and *G*_*in*,12_ = *G*_*in*,21_ = 0.05. The network natural frequency is *f*_*ntwk,nat*_ *∼* 14.44.

The *K*-curves (Fig. 17) are also similar to the corresponding ones in Section 3.5.1, but they show some differences for the smaller values of the input frequency *f*. In Case A, the *K*-curve presents a self intersection and three frequencies at which the position and tangent vectors are aligned. In contrast, in Case B, there is no self-intersection. However, unlike the situation in the mutually inhibitory network of PNs (subsection 3.5.1, Case B), the behavior for small input frequencies allows for the existence of an additional frequency alignment point between the position and tangent vectors. These observations explain the existence of resonances and antiresonances in the corresponding *K*-profiles (Fig. 17-C). In Case A, the *K* values at the resonant and antiresonant frequencies are relatively close to each other (Fig. 17-C, blue), while in Case B, these differences are larger (Fig. 17-C, pink). In both cases, the antiresonant frequency is closest to the largest one of the isolated cells.

**Figure 17:**
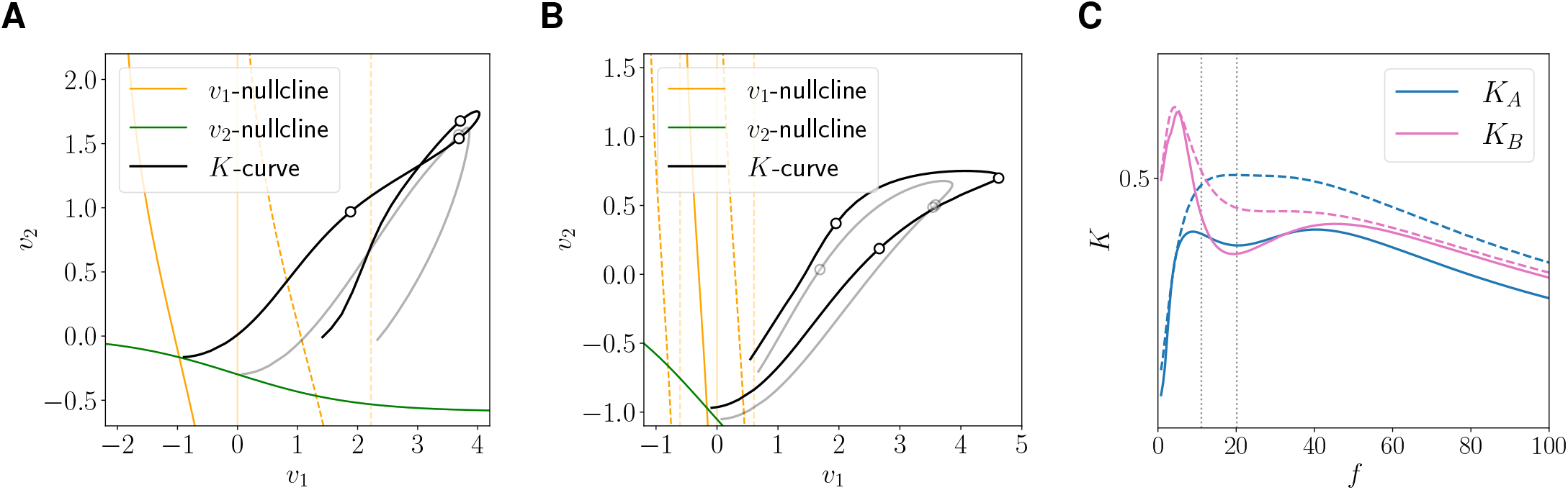
Two mutually inhibitory resonant 2D nodes (RN1 and RN2): *K*-curves and *K*-profiles. We used eqs. (52)-(53) with the parameter values as in Fig. 16. **A, B**. Phase-plane diagram for the examples considered in Fig. 16. The solid orange and green lines are projections of the *v*_1_ and *v*_2_-nullcline of the autonomous system. The dashed orange lines are the extended *v*_1_-nullclines for the autonomous system for constant inputs *±A*_*in*_. The black curve is the *K*-curve it joins the peak values of the response limit cycles in the *v*_1_ and *v*_2_ directions. The dots indicate the presence of *K*-resonance and antiresonance. **C**. *K*-profiles for case A and B. Feedforward inhibitory (dashed) and mutual inhibitory (solid) networks. The vertical dotted lines correspond to the resonant frequencies of the isolated cells (see Fig. 16).

Interestingly, in case A, both the resonance and the antiresonance are created by feedforward inhibition (Fig. 17-C, dashed-blue) and they become more prominent by the addition of feedback inhibition (Fig. 17-C, solid-blue). In contrast, in case B, feedforward inhibition creates a broad resonant band (Fig. 17-C, dashed-pink), which transitions into a resonant-antiresonant pattern (Fig. 17-C, solid-pink) by the addition of feedback inhibition.

Fig. 18 compares the *K*-profiles for three representative mutually inhibitory network scenarios where the cell two is a resonator (RN1) mutually connected to an identical resonator (orange), another resonator (RN2, purple) and a passive cell (PN, green). As the value of 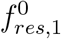 (for the isolated cell 1) increases, the values of the resonant and antiresonant frequencies also increase. The peak-to-trough amplitude of the *K*-profiles increases from the green (PN) through the orange (RN1) to the purple (RN2) scenarios. comm

**Figure 18:**
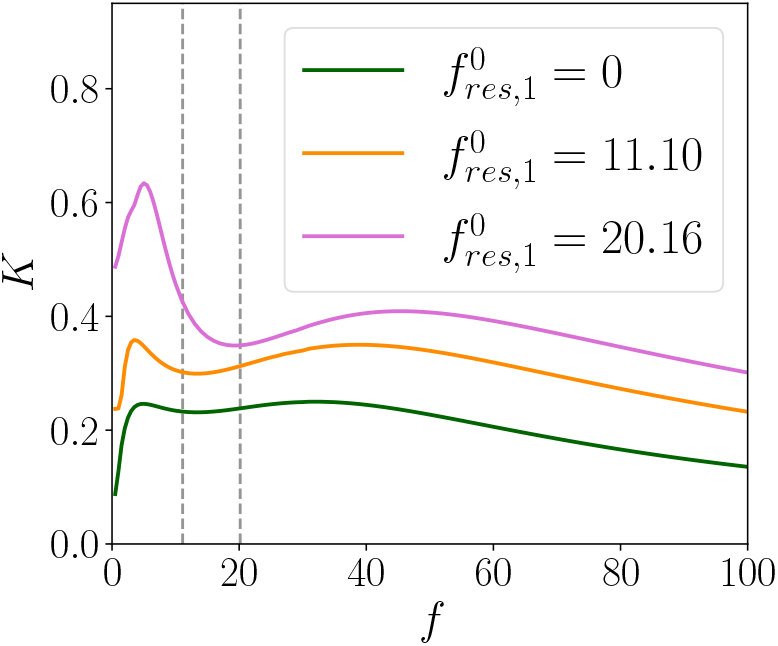
Comparison of the *K*-profile for three representative mutually inhibitory network scenarios. We used (52)- with *G*_*in*,12_ = *G*_*in*,21_ = 0.05. The three networks consist of a resonator (RN1) connected to an identical resonator (RN1), another resonator (RN2) and a passive cell (PN). The cell receiving the input (cell 1) changes and the cell 2 is always the RN1. The parameter values for the resonant cells (RN1 and RN2) are as in Fig. 16. The parameter values for the passive cell (PN) are *g*_*L*_ = 0.259 and *g* = 0. Values of 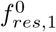 for the isolated cell 1 (vertical dotted).

If the inhibition between the cells increases or the values of the leak conductance *g*_*L*_ decrease, the resonances and antiresonance in the communication coefficient *K* are more pronounced. But, by modifying these parameter the network shows intrinsic sustained oscillations and our analysis of impedance profiles ceases to be valid. In addition, if the parameter *g*_*k*_ is increased, the corresponding isolated cell could show either an intrinsic natural frequency or, in the network, the equilibrium coexists with a limit cycle. Here too, in both cases the analysis developed before ceases to be valid.

## 4 Discussion

Neurons and networks of neurons exhibit preferred frequency responses to oscillatory inputs in amplitude (resonance) and phase (phasonance) [1,4,19,20,26,32,35]. Preferred frequency responses can be inherited from one level of organization to another (e.g., from neurons to networks) or be created at different levels of organization in an independent manner by a variety of mechanisms [4]. Two important mechanistic issues that remain poorly understood are (i) whether and under what conditions a neuronal network exhibits resonance and phasonance in one or more of the participating neurons and (ii) how input signals are communicated between neurons in the network in a frequency-dependent manner. In this paper we set out to investigate these issues by using minimal network models (Fig. 1) consisting of two cells recurrently connected via graded synaptic inhibition or excitation where one of the cells receives oscillatory input currents within a frequency range. We used analytical calculations, numerical simulations and we developed dynamical system tools for the qualitative analysis and biophysical interpretation of the nonlinear models where analytical calculations were not possible.

The individual neurons are either passive cells (low-pass filters, 1D) or resonators (band-pass filters, 2D), defined as neurons that exhibit resonance in the absence of intrinsic oscillations. This choice was motivated by our desire to separate the effects of resonance from other possibly interfering effects. We purposely left out of the scope of this paper the study of networks having more complex intrinsic dynamics (e.g., intrinsic bistability and oscillations [43, 50]) in order to identify the basic dynamic principles that control the generation of preferred network responses to oscillatory inputs.

The response of a neuron receiving an oscillatory input current is typically measured in terms of the impedance function, which allows to study the amplitude and phase frequency content of this response. In linear models, the impedance amplitude profile is equivalent to the voltage peak-to-trough envelope profiles. However, this is not the case for nonlinear models. Because neuronal signals are generated through voltage threshold mechanisms, we examined the peak envelope profiles for each cell in the network to compute a generalized version of the impedance and phase profiles, and used these profiles to understand the frequency-dependent properties of the network response to oscillatory inputs.

To study the properties of the communication of oscillatory information between neurons in a network, we borrowed the notion of coupling coefficient for electrically coupled networks (gap junctions) in response to external inputs [52–57, 59, 60, 65, 66] and defined the communication coefficient *K* as the ratio between the responses of the indirectly and directly activated neurons. We analyzed the frequency-dependent properties of *K* in terms of these of the participating neurons and the conditions under which the nodes function as filters that either facilitate or hinder the transmission in synaptically connected networks. We also analyzed the synchronization and frequency-dependent phase-locking properties of the responses using the phase coefficient ΔΦ. For linear networks (linear nodes and linear connectivity, *K* and ΔΦ constitute the complex communication coefficient.

In order to understand how the network response and communication profiles are controlled by the intrinsic properties of the participating cells and the synaptic connectivity, we extended the dynamical systems methods introduced in [37] and developed the notion of the *K*-curve, a curve parametrized by the input frequency, which is a geometric representation of the coefficient *K* in the phase-space diagram.

We first considered two-cell networks with linear connections and a sinusoidal input to cell 1. The response of linear networks to sinusoidal input is analytically solvable using standard tools, as shown in the Supplementary Material S4. As we showed, the network impedance profile of each neuron can exhibit resonance, and cell 1 can have a phasonance. However, the communication coefficient (for 1D or 2D linear nodes) is always a multiple of the individual extended impedance of cell 2, and the phase difference equals the individual extended phase of cell 2. Thus, the frequency-dependent transmission of information in the network from cell 1 to cell 2 is inherited from cell 2, and independent of cell 1. Specifically, in the case where both isolated nodes are passive (1D), the connected nodes may show a preferred amplitude frequency, but the *K* does not; It acts as a low-pass filter, and the nodes do not have synchronized in phase peaks. In contrast, if node 2 is a resonator, the information is transmitted with a preferred frequency close (or equal if there is no self-connection) to the resonance frequency of the isolated node 2, and both neurons have synchronized in phase peaks at the frequency where node 2 shows a phasonance.

In networks with graded synaptic (nonlinear) connections, the network’s response and particularly the behavior of the communication coefficient becomes more complex, potentially functioning as a low-pass or a band-pass filter, with or without antiresonances. We studied this coefficient in networks with two (1D or 2D) cells and different architectures (see diagrams in Fig. 1). We used the *K*-curve (defined as the curve joining the peak values of the responses to oscillatory inputs) to study the communication coefficient *K* as the intrinsic or connection parameters vary. In all cases, the attributes of the communication coefficient *K* are associated with how the network response (and so the *K*-curve) (parametrized by the input frequency) traverses the phase-space as the input frequency increases and how the nonlinearity modifies the shape of this trajectory. Not only the input frequency is important, but also its amplitude, which allows the nonlinearity to be expressed in the responses of each node. The nonlinearities have a greater impact in the filter properties of the communication coefficient *K* for small frequencies (smaller than 50 Hz) for the scenario considered. At high frequencies, both responses tend to shrink around the equilibrium, and so the coefficient *K* tends to zero.

More specifically, in networks of passive cells, the feedforward inhibition or excitation (from cell 1 to cell 2) generates a preferred frequency for information transmission (*K*-resonance), i.e., it maximizes the amplitude of cell 2 relative to cell 1. And, as expected, more inhibition or excitation from cell 1 to cell 2 (increasing values of *G*_*syn*,21_) increases information transmission through the network. However, how the feedback affects this transmission depends on the feedforward connection. If the feedforward is inhibitory, both types of feedback amplify the resonance amplitude (*Q*_*K*_). If the feedforward is excitatory, the feedback connection attenuates the *K*-profiles and the resonance, if it exists, becomes less significant since the amplitude variation in cell 2 is small compared to that in cell 1. The intrinsic properties of the cells also affect the results. In the mutual inhibitory network if the conductance of cell 1 (*g*_*L*,1_) increases, the coefficient reaches higher values and becomes a low-pass filter (because *K*(0) increases), whereas if the conductance of cell 2 (*g*_*L*,2_) increases, the values of *K* decrease but a resonance (of small amplitude) is generated.

In the case of networks where at least one node is resonant, the inhibitory connection generates network resonance in both nodes that is lower than that of the isolated node, or between the resonance frequencies if both nodes are resonators. The communication coefficient exhibits an antiresonance at frequencies close to the resonance frequency of the resonant node (or to the higher resonance frequency of the isolated node, if both are resonant). Moreover, the coefficient can show two resonances at frequencies near the antiresonance. The combination of resonances and antiresonances produces a frequency range in which the coefficient *K* reaches values close to its maximum. In inhibitory networks with symmetric connections, we observe that the coefficient *K* reaches higher values, mainly at low frequencies, when the node receiving the input (cell 1) has a higher resonant frequency than the connected node (cell 2).

Thus, we show that the nonlinearities associated to graded-type connections generates greater variability in the transmission of information between nodes in the network, and that this depends both on the properties of the nodes and on the intensity and type of connection between them. We observe that the nonlinearity is more strongly expressed in the responses of each node and in the transmission when the input has low frequencies and sufficiently large amplitudes. Beyond a certain frequency, the effects of the nonlinearities become less noticeable, the responses shrink around the system’s equilibrium, and the communication coefficient decreases to zero.

In this paper we studied minimal networks with the goal of identifying the basic mechanisms that control the frequency-dependent behaviors of the nodes’ response to oscillatory inputs and the coefficients *K* and ΔΦ and developing the appropriate tools for this study. Future research should focus on more complex networks having nonlinear nodes and more intricate architectures, and the interaction of multiple inputs to the network with possible different frequency content acting simultaneously. This requires extending the dynamical systems methods to account for the presence of multiple fixed-points as well as intrinsic oscillations.

Our results and the dynamical systems tools developed here have implications for networks of electrically coupled neurons where the coupling coefficient was used to characterize the effective strength and transmission of information in response to constant [51–65] and oscillatory inputs [65, 66]. Because gap junctions are linear, the basic properties of the coupling coefficient were analytically illustrated by using linear models. However, our results suggest that the presence of nonlinearities generated by the intrinsic ionic currents in the participating nodes may generate significant qualitative departures from the linear prediction, which remain to be investigated.

Because of the properties of instantaneously fast graded synapses [41–49] our results have direct implications for networks of firing rate models of Wilson-Cowan type (1D nodes) [68] and Wilson-Cowan type with adaptation (2D nodes) [47–49]. The communication coefficients can be defined and used in these networks to investigate the frequency-dependent properties of the communication of information between nodes in response to oscillatory inputs and the dynamical systems tools developed here can be extended to understand the underlying mechanisms. Additional research is needed to investigate whether the observed phenomena persist in networks of spiking neurons.

## Acknowledgments

AB acknowledges supports from the Universidad Nacional del Sur grant PGI 24/L131 and CONICET, Argentina. HGR acknowledges support from the National Science Foundation grants DMS-1608077 and IOS-2002863. HGR is a Corresponding Researcher in CONICET, Argentina and a Graduate Faculty Member in the Graduate Program in Neuroscience (GPN) in the Center for Molecular and Behavioral Neuroscience (CMBN) at Rutgers University.

## Supplementary Material

### S1 Linearization of single-cell conductance-based models

#### S1.1 Conductance-based models: single cell

We use the following biophysical (conductance-based) models of Hodgkin-Huxley type [75, 76] to describe the neuronal subthreshold dynamics

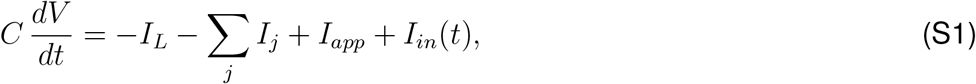

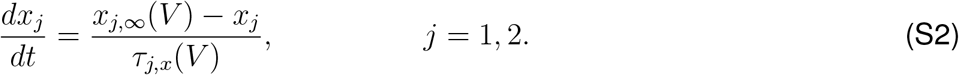

In the current-balance equation (S1), *V* is the membrane potential (mV), *t* is time (ms), *C* is the membrane capacitance (*µ*F/cm^2^), *I*_*app*_ is the applied bias (DC) current (*µ*A/cm^2^), *I*_*L*_ = *G*_*L*_ (*V − E*_*L*_) is the leak current, and *I*_*j*_ = *G*_*j*_ *x*_*j*_, (*V − E*_*j*_) are generic expressions for ionic currents (with *j* an index) with maximal conductance *G*_*j*_ (mS/cm^2^) and reversal potentials *E*_*j*_ (mV) respectively. The dynamics of the gating variables *x*_*j*_ obey the kinetic equations (S2) where *x*_*j,∞*_(*V*) and *τ*_*j,x*_(*V*) are the voltage-dependent activation/inactivation curves and time constants respectively. For simplicity, the generic ionic currents *I*_*j*_ we consider here are restricted to have a single gating variable *x*_*j*_ and to be linear in *x*_*j*_. This is typically the case for persistent sodium (*I*_*Nap*_), *h*- (hyperpolarization-activated, mixed-cation, inward; *I*_*h*_), and slow-potassium (M-type) (*I*_*Ks*_) currents. Our discussion and results can be easily adapted and generalized to include ionic currents having two gating variables [19, 38, 77, 78] raised to powers different from one such as T-type calcium and A-type potassium currents [1]. The input current *I*_*in*_(*t*) (*µ*A/cm^2^) in eq. (S1) has the form (5), *I*_*in*_(*t*) = *A*_*in*_ *sin*(Ω *t*) with Ω = 2 *π f /* 1000 (*f* is the input frequency in Hz).

Here we focus on 2D models describing the dynamics of *V* and one gating variable. Additional currents whose gating variables evolve on a very fast time scale (as compared to the other variables) can be included by using the adiabatic approximation *x*_*j*_ = *x*_*j,∞*_(*V*). Here we include one such fast current *I*_2_ = *G*_2_ *x*_2,*∞*_(*V*) (*V − E*_2_). Additional fast currents can be included without significantly changing the formalism used here.

#### S1.2 Linearization: conductance-based linearized models

Linearization of the autonomous part (*I*_*in*_(*t*) = 0) of system (S1)-(S2) around the fixed-point 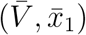 yields [19, 26, 77]

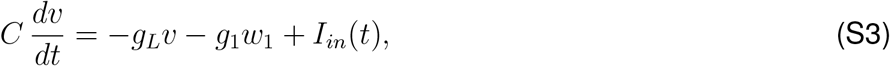

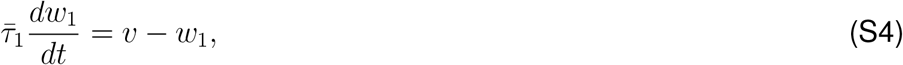

where

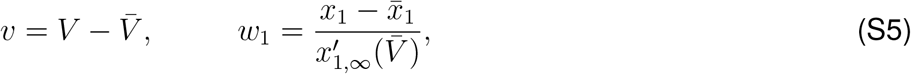

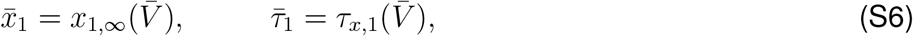

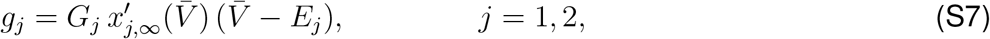

and

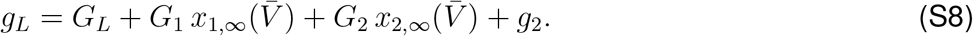

Note that the gating variables *w*_1_ and *w*_2_ in (S5) have units of voltage ([*v*] = [*w*_1_] =V).

The effective leak conductance *g*_*L*_ (S8) contains information about the biophysical leak conductance *G*_*L*_, the ionic conductances, and their associated voltage-dependent activation / inactivation curves. The fast ionic current *I*_2_ contributes to *g*_*L*_ with an additional term *g*_2_. The signs of the effective ionic conductances *g*_*j*_ determine whether the associated gating variables are resonant (*g*_*j*_ *>* 0) or amplifying (*g*_*j*_ *<* 0) [1, 19]. Specific examples are the gating variables associated to *I*_*h*_ (resonant), *I*_*Ks*_ (resonant), *I*_*Nap*_ (amplifying) and *I*_*Kir*_ (amplifying). All terms in *g*_*L*_ are positive except for the last one that can be either positive or negative. Specifically, *g*_*L*_ can become negative for negative enough values of *g*_2_.

All the linearized conductances are affected not only by the respective biophysical conductances, but also by the magnitudes and signs of the activation (*σ <* 0) and inactivation (*σ >* 0) curves and their derivatives, which are typically given by expressions of the form

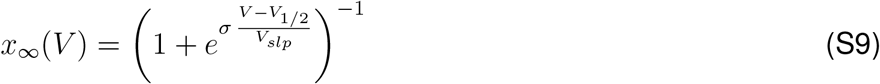

and

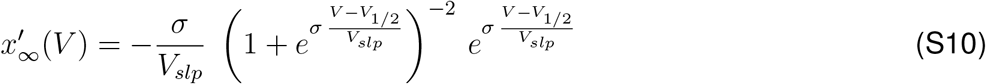

where *V*_1*/*2_ and *V*_*slp*_ *>* 0 are constants.

### S2 Impedance amplitude and phase profiles for 2D linear systems: individual cells

For a linear system

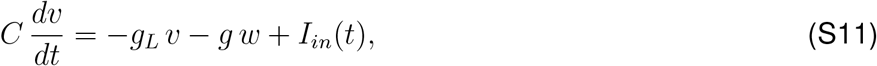

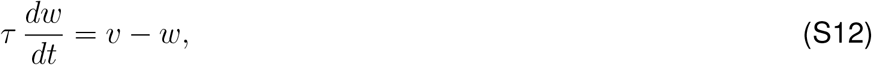

receiving a sinusoidal input current of the form (5)

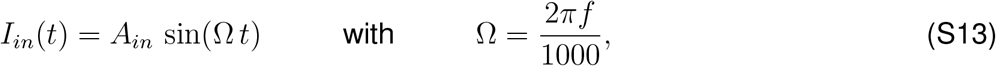

the voltage response is given by

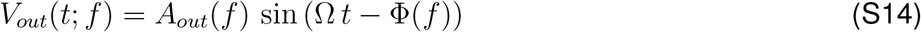

where *A*_*out*_(*f*) is the amplitude and Φ(*f*) is the phase-shift (or phase), which captures the difference between the peaks of *I*_*in*_(*t*) and *V*_*out*_(*t*; *f*) normalized by the period. For the linear system (S11)-(S12), the impedance is given by [19, 26, 77])

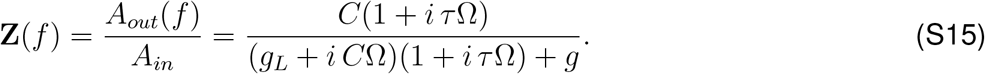

The *Z*- and Φ-profiles are given, respectively by

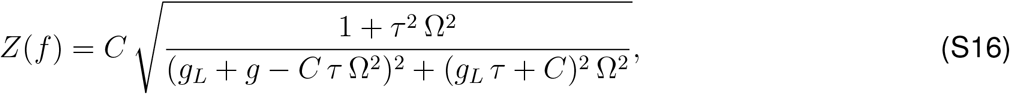

and

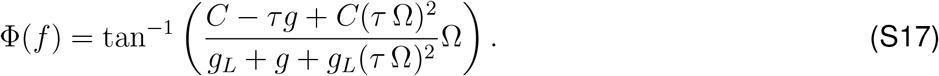

System (S11)-(S12) exhibits *resonance* (band-pass filter) if *Z*(*f*) peaks at a non-zero (resonant) frequency *f*_*res*_ (Figs. 2-A2) and *phasonance* if Φ(*f*) vanishes at a non-zero (phasonant) frequency *f*_*phas*_ (Figs. 2-C2).

From (S16), the resonant frequency is given by

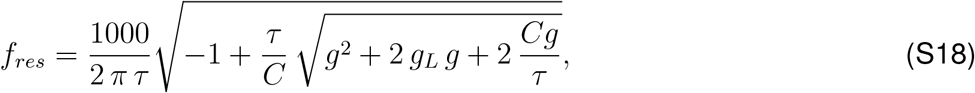

and the impedance peak *Z*_*max*_ is given by

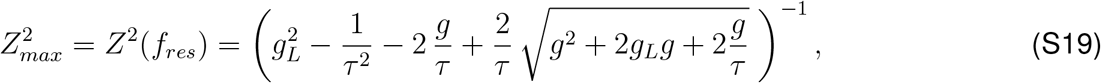

where for simplicity we used *C* = 1. From (S17), the phasonant frequency is given by

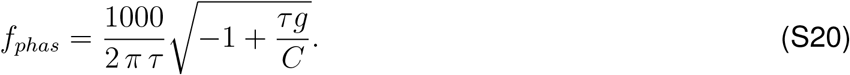

For passive (one-dimensional) cells (*g* = 0), the *Z*-profile reduces to a low-pass filter

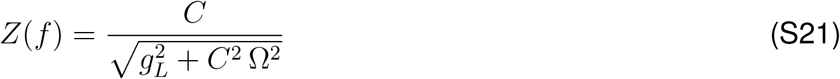

and the Φ-profile reduces to the positive increasing function

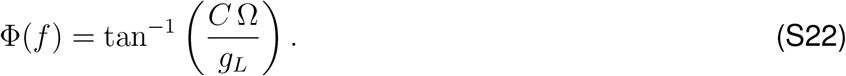

The eigenvalues for the autonomous part of system (S11)-(S12) (no inputs) are given by

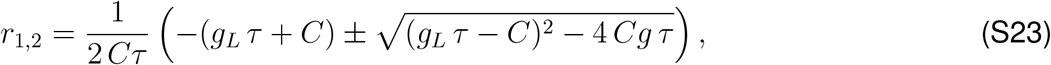

and the natural frequency, if it exists, is given by

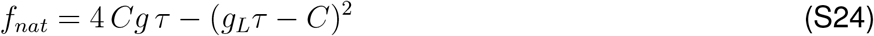

provided this quantity is positive.

### S3 Two-1D-cell network: Resonant and phasonant frequencies, and eigenvalues

#### S3.1 Resonant and phasonant frequencies

In Fig. S1 we illustrate the dependence of the natural (32), resonant (35)-(36), and phasonant (37) frequencies for the two cells in the network on the cross-connectivity parameters for representative values of *g*_*L*,1_ and *g*_*L*,2_, which control the shape of the *Z*- and Φ-profiles of the individual cells. Because in all cases this dependence is through the product Γ_12_Γ_21_, we use this product as the control parameter (abscissa). In all cases, the characteristic frequencies are increasing functions of the cross-connectivity strength |Γ_12_Γ_21_|, but the relative values of these frequencies vary as well as the ability of the network to exhibit one phenomenon in the absence of the other(s).

For the limiting case *g*_*L*,1_ = *g*_*L*,2_ = 0, then *f*_*ntwk,res*,1_ = *f*_*ntwk,phas*,1_ = *f*_*ntwk,res*,2_ = *f*_*ntwk,nat*_ for all values of the cross-connectivity product Γ_12_Γ_21_. If the individual cells are identical (*g*_*L*,1_ = *g*_*L*,2_), from (32), *f*_*ntwk,nat*_ depends only on the cross-connectivity product, and from (36) and (37), *f*_*ntwk,res*,2_ = *f*_*ntwk,phas*,1_ (not shown). Therefore we focus on relatively large and different values of *g*_*L*,1_ and *g*_*L*,2_.

For the parameter values in Fig. S1-A, intrinsic damped-oscillations may occur in the absence of network resonance. As Γ_12_Γ_21_ decreases, first cell 1 exhibits network resonance, then cell 1 exhibits network phasonance, and finally cell 2 exhibits network resonance. As the difference between *g*_*L*,1_ and *g*_*L*,2_ increases (Fig. S1-B), the green curve falls completely below the blue one, and cell 1 can exhibit network resonance in the absence of intrinsic damped oscillations. As this difference increases further (Fig. S1-C), *f*_*ntwk,phas*,1_ increases above *f*_*ntwk,nat*_, and therefore cell 1 can exhibit both network resonance and phasonance in the absence of intrinsic damped oscillations. In the limiting case *g*_*L*,2_ = 0, from (35) and (37), *f*_*ntwk,res*,1_ = *f*_*ntwk,phas*,1_ and the corresponding curves are superimposed (not shown). While this case is not realistic, for small enough values of *g*_*L*,2_, the corresponding curves for *f*_*ntwk,res*,1_ and *f*_*ntwk,phas*,1_ are almost superimposed (not shown).

In Fig. S1-D, the difference between *g*_*L*,1_ and *g*_*L*,2_ is the same as in Fig. S1-A, but with a different sign. This causes *f*_*ntwk,res*,2_ and *f*_*ntwk,phas*,1_ to be inverted in comparison to Fig. S1-A, and therefore cell 2 can exhibit network resonance in the absence of network phasonance in cell 1. This behavior persists as the difference between *g*_*L*,1_ and *g*_*L*,2_ increases (Fig. S1-E). When the ratio decreases, the relative position of the curves remains the same, but more “squeezed” (Fig. S1-F), at least for a significantly large range of parameter values.

#### S3.2 Characteristic polynomial for a two-cell network

The characteristic polynomial for a two-cell network of the form (S28)-(S29) is given by

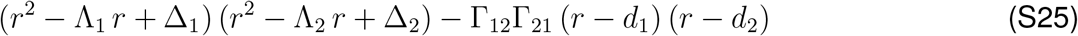

where

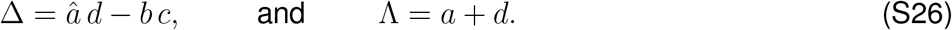

For two-cell networks of 1D cells, the characteristic polynomial reduces to

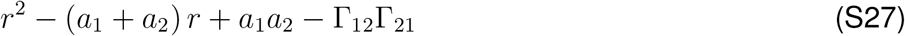

**Figure S1:**
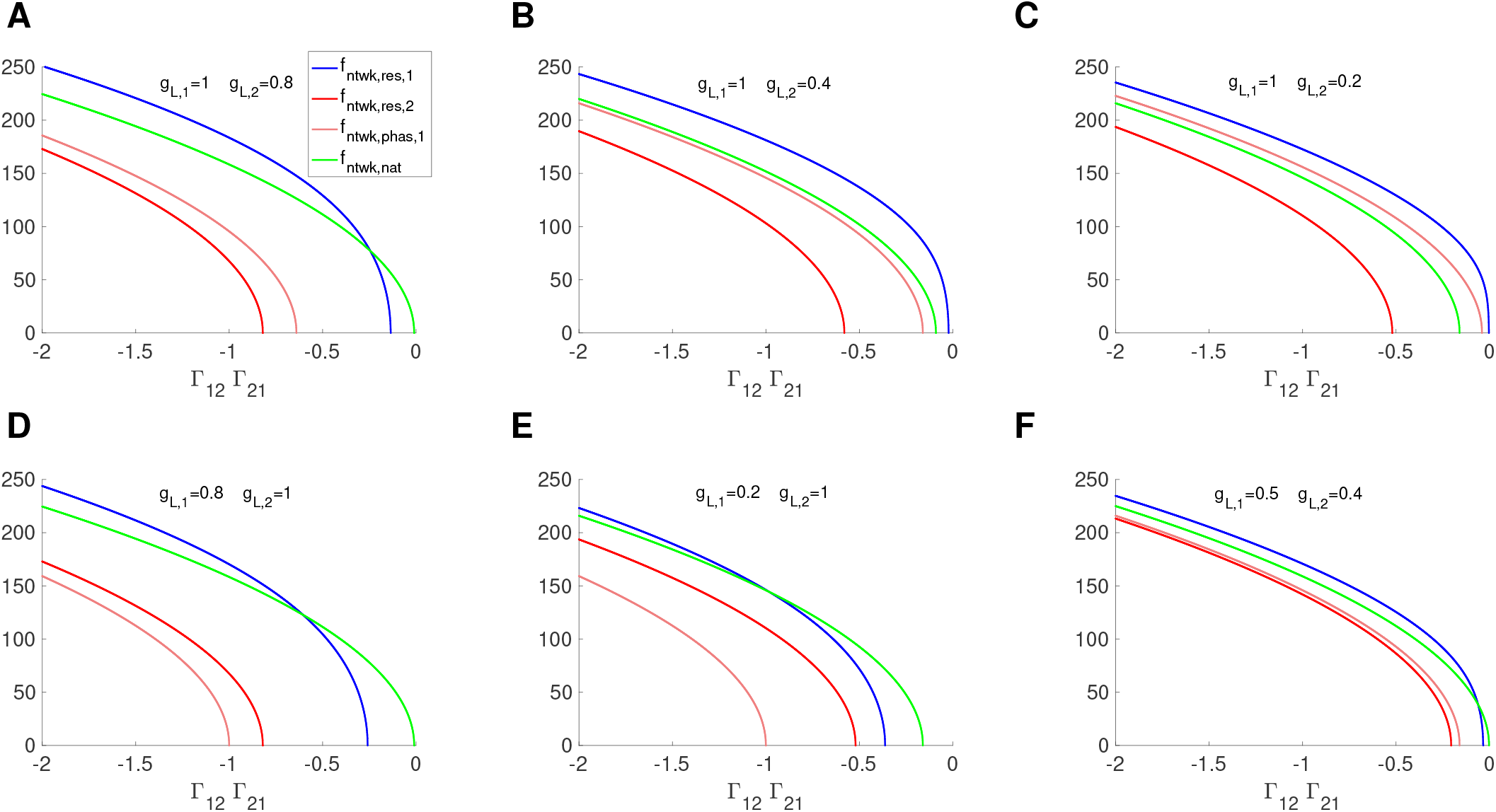
Dependence of the network characteristic frequencies on the cross-connectivity parameters for two-1D-cell linear networks. The natural, resonant and phasonant frequencies are given by eqs. (32), (35), (36) and (37), respectively. We used the following additional parameter values: Γ_11_ = Γ_22_ = 0 (Γ_11_ and Γ_22_ can be thought of being absorbed in *g*_*L*,1_ and *g*_*L*,2_, respectively).

### S4 Network impedance profiles for linear systems: networks with linear nodes and connectivity

We consider networks consisting of *N* nodes with 2D linear dynamics and linear connectivity. The dynamics are described by

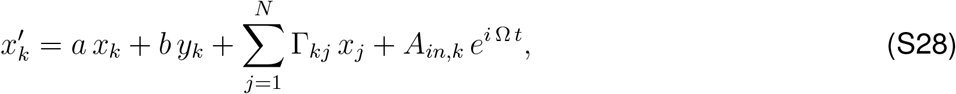

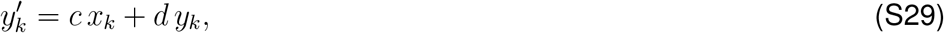

for *k* = 1, …, *N*. In (S28)-(S29), *a, b, c* and *d* are constant, Ω *>* 0 is the input frequency and *A*_*in,k*_ ≥ 0 are the input amplitudes. The connectivity coefficients Γ_*kj*_ indicate a connection from cell *j* to cell *k*. The coefficients *a, b, c* and *d* may be different in different nodes in the network. For simplicity in the notation we omit the subindex, which will be reintroduced later in the notation for the impedance profiles.

We use the notation **Z**_**k**_(Ω), *Z*_*k*_(Ω) and Φ_*k*_(Ω) (*k* = 1, …, *N*) to refer to the (complex) impedance, impedance amplitude (or simply, impedance) and phase profiles, respectively, of the individual nodes. The analytical expressions for 2D and 1D (*b* = 0) models are given in the Appendix S2.

The equation (S28) can be rewritten as

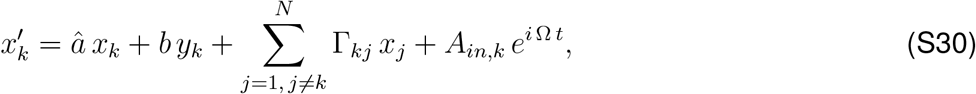

for *k* = 1, …, *N* where *â* = *a* + Γ_*kk*_ includes the coefficient (Γ_*kk*_) of the autonomous component of the coupling term (Γ_*kk*_ *x*_*k*_). We use the notation 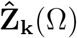 to refer to the (extended) impedance profile of the individual nodes including this term; i.e., the impedance profiles of the system consisting of the autonomous component in (S30) and equation (S29). Similarly, 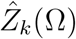 and 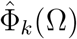 refer to the extended impedance amplitude and phase profiles of these systems, respectively.

The particular solution to system (S28)-(S29) has the form

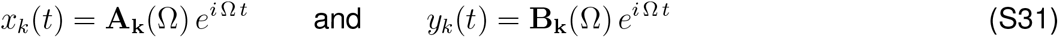

for *k* = 1, …, *N*. Substituting (S31) into (S29)-(S30) and rearranging terms we obtain

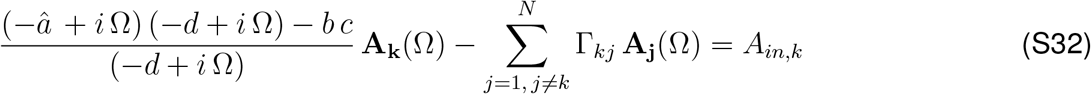

for *k* = 1, …, *N*. From (S42) in the Appendix S2 with *a* substituted by *â* we obtain

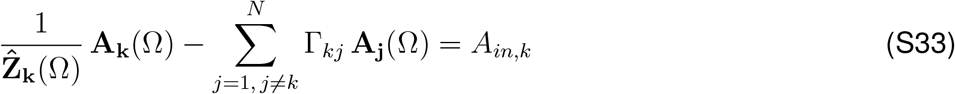

for *k* = 1, …, *N*.

This is a complex linear system of algebraic equations that can be written in compact form as

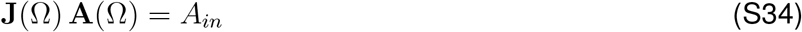

where

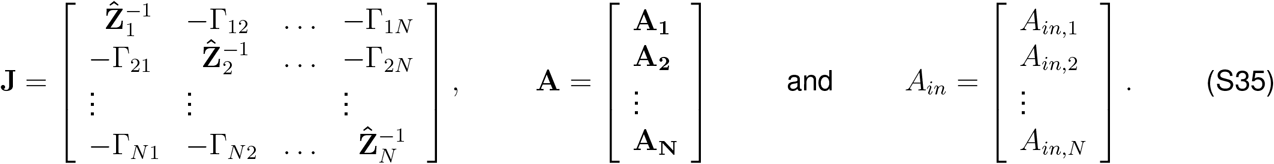

Note that for simplicity the explicit dependence of **J** and **A** on Ω has been omitted in (S35).

The solution to (S34) is formally given by

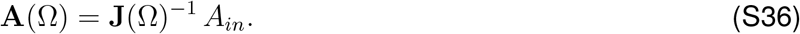

In order to invert the complex matrix **J**(Ω) one needs to express 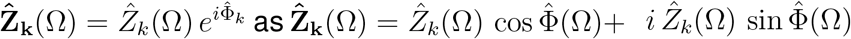 and **A**_k_(Ω) = *A*_*k,real*_(Ω)+*iA*_*k,imag*_(Ω). Alternatively, **A**_**k**_ = |*A*_*k*_(Ω) |*e*^*i*Ψ(Ω)^ = |*A*_*k*_(Ω) |cos Ψ(Ω)+ *i A*_*k*_(Ω) sin Ψ(Ω).

The vector **A**(Ω) represents the response of the network to the oscillatory inputs. For a two-cell network (*N* = 2) we obtain

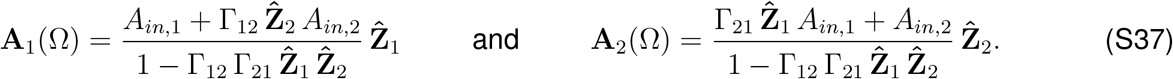

In order to extend the concept of impedance to networks we need to find an appropriate normalization. In the impedance for the individual neurons, this normalization is given by the input amplitude. If the input arrives only to one node in the network, it is natural to choose this input amplitude as the normalization constant. Otherwise, one has to choose an appropriate one among the various input amplitudes. Here we choose the largest of all and we index the sequence of input amplitudes in decreasing order of magnitude. Thus, eq. (S36) reads

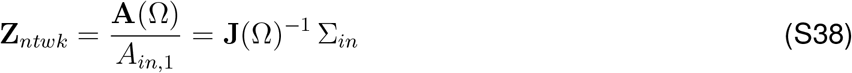

where Σ_*in*_ = [1, *σ*_2_, …, *σ*_*N*_]^*T*^ with *σ*_*k*_ = *A*_*in,k*_*/A*_*in*,1_ (*k* = 2, …, *N*). For a two-cell network, we obtain eqs. (10)-(11).

Assuming we have chosen an appropriate normalization, we call the *Z*_*ntwk,k*_(Ω) and Φ_*ntwk,k*_(Ω) (*k* = 1, …, *N*) the network impedance amplitude and phases, respectively. While one could conceive other, global measures of the voltage response, such as the addition of the responses of the two cells or the quotient between the two, the one we choose is the most informative for the purpose of this paper.

### S5 Nonlinear networks: Phase profiles and phase communication coefficient

**Figure S2:**
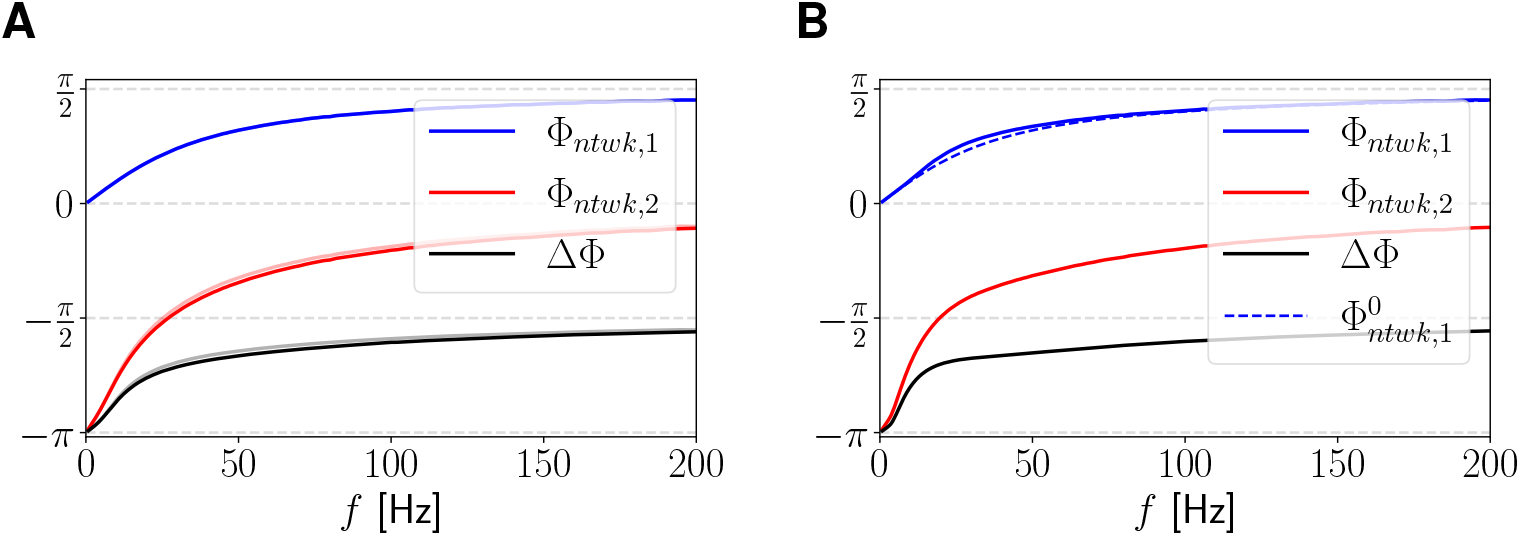
Two-1D-cell network with feedforward and mutual inhibition: Phases and ΔΦ-profiles. Network phase profiles (solid), individual phase profiles (dashed) and ΔΦ-profile (black). We used the system (45)-(46) receiving an oscillatory inputs only to cell 1 (*A*_*in*,1_ = *A*_*in*_ and *A*_*in*,2_ = 0). The self-connectivity parameters (if they exist) are considered to be absorbed by the autonomous linear terms (*G*_*syn*,11_ = *G*_*syn*,22_ = 0) and *C*_1_ = *C*_2_ = 1. **A**. Feedforward inhibition: *G*_*in*,12_ = 0 and *G*_*in*,21_ = 0.1 (light colors *G*_*in*,21_ = 0.01). **B**. Mutual inhibition: *G*_*in*,12_ = 0.05 and *G*_*in*,21_ = 0.1. We used the following additional parameter values: *g*_*L*,1_ = *g*_*L*,2_ = 0.2, *E*_*syn*,12_ = *E*_*syn*,21_ = *−*20 and *A*_*in*_ = 1.

**Figure S3:**
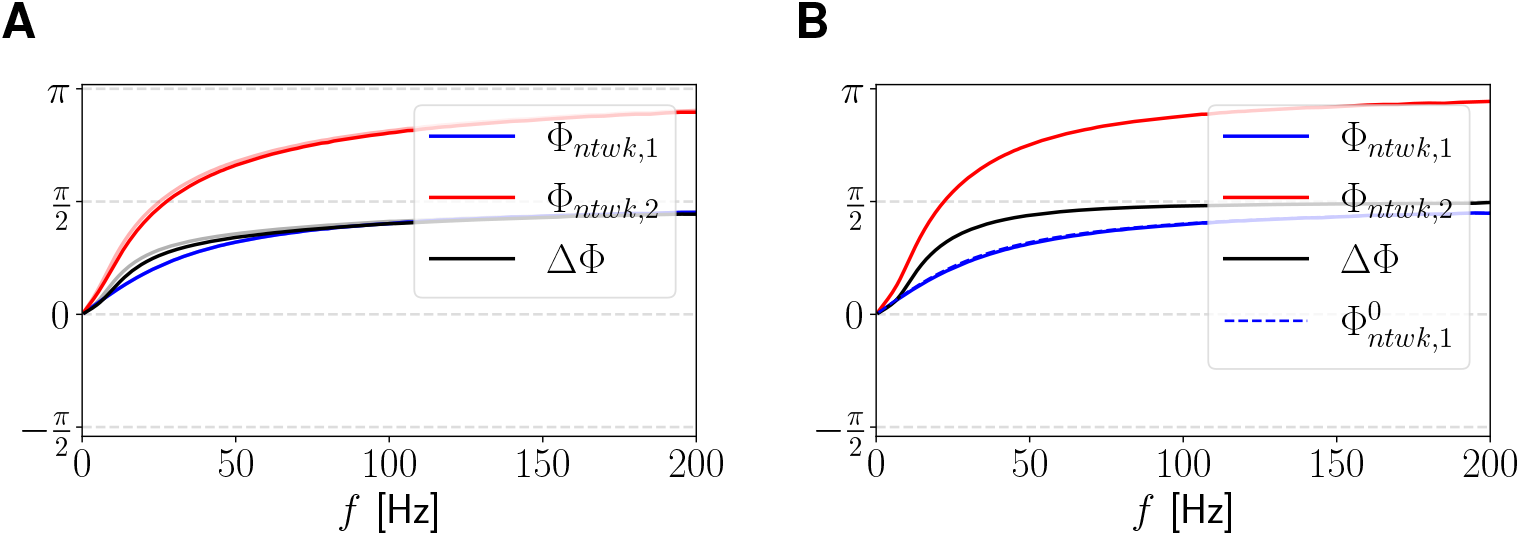
Two-1D-cell network with feedforward and mutual excitation: Phases and ΔΦ-profiles. Network phase profiles (solid), individual phase profiles (dashed) and ΔΦ-profile (black). We used the system (45)-(46) receiving an oscillatory inputs only to cell 1 (*A*_*in*,1_ = *A*_*in*_ and *A*_*in*,2_ = 0). The self-connectivity parameters (if they exist) are considered to be absorbed by the autonomous linear terms (*G*_*syn*,11_ = *G*_*syn*,22_ = 0) and *C*_1_ = *C*_2_ = 1. **A**. Feedforward excitation: *G*_*ex*,12_ = 0 and *G*_*ex*,21_ = 0.05 (light colors: *G*_*ex*,21_ = 0.005). **B**. Mutual excitation: *G*_*ex*,12_ = 0.01 and *G*_*ex*,21_ = 0.05. We used the following additional parameter values: *g*_*L*,1_ = *g*_*L*,2_ = 0.2, *E*_*syn*,12_ = *E*_*syn*,21_ = 60 and *A*_*in*_ = 1.

**Figure S4:**
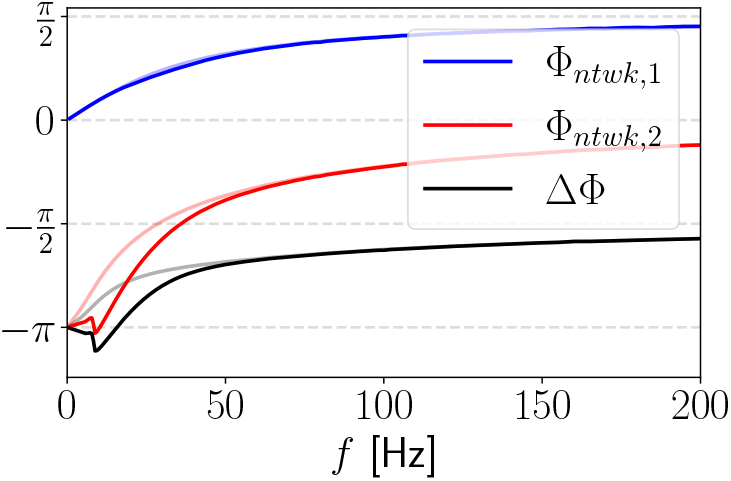
Two-1D-cell inhibitory-excitatory network: Phases and ΔΦ-profiles. Network phase profiles (solid) and ΔΦ-profile (black). We used equations (45)-(46) with *A*_*in*_ = 1 and the following additional parameter values: *g*_*L*,1_ = *g*_*L*,2_ = 0.2, *G*_*in*,21_ = 0.2, *G*_*ex*,12_ = 0.05 (light colors: feedforward inhibition, i.e., *G*_*ex*,12_ = 0), *E*_*syn*,21_ = *−*20 and *E*_*syn*,12_ = 60.

**Figure S5:**
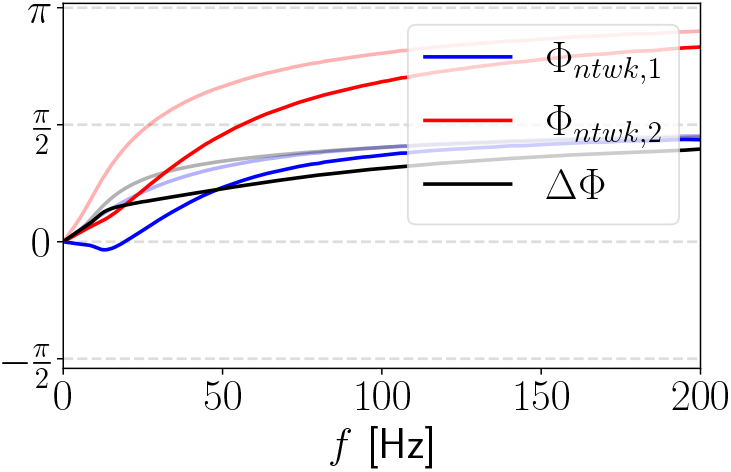
Two-1D-cell excitatory-inhibitory network. Phases and ΔΦ-profiles. Network phase profiles (solid) and ΔΦ-profile (black). We used equations (45)-(46) with *A*_*in*_ = 1 and the following additional parameter values: *g*_*L*,1_ = *g*_*L*,2_ = 0.2, *G*_*ex*,21_ = 0.04, *G*_*in*,12_ = 0.1 (light colors: feedforward excitation, i.e., *G*_*in*,12_ = 0), *E*_*syn*,21_ = 60 and *E*_*syn*,12_ = *−*20.

**Figure S6:**
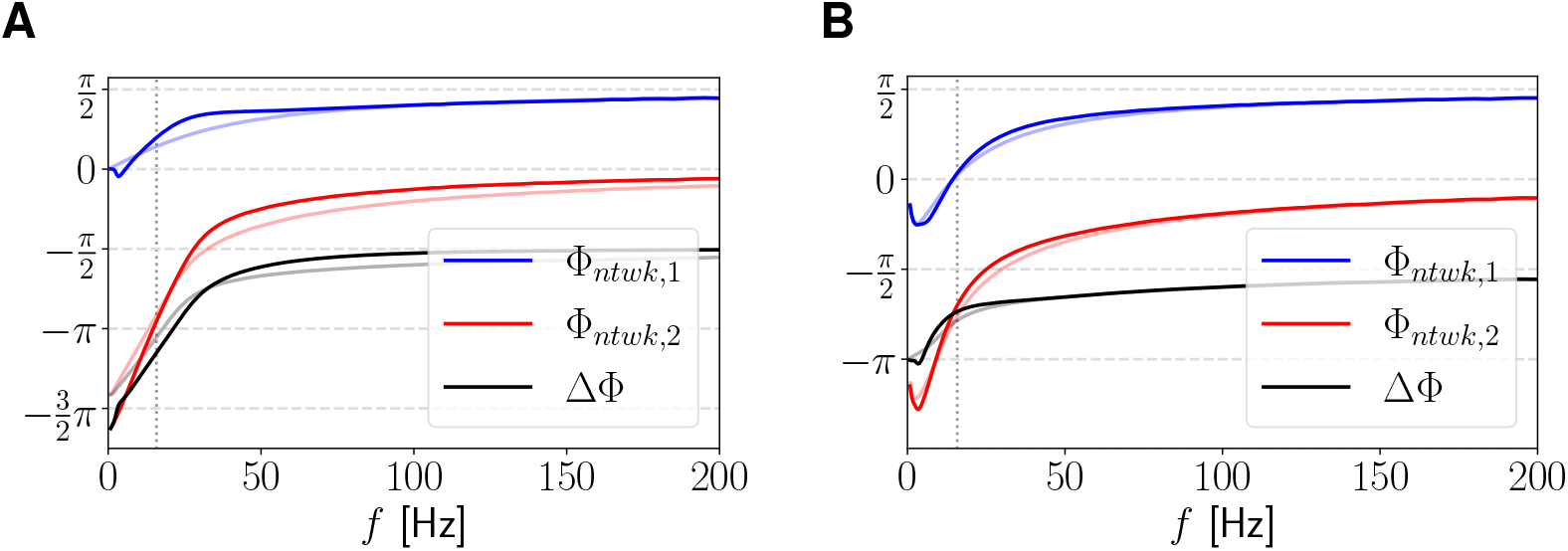
Mutually inhibitory resonant (RN) and passive (PN) nodes. Phases. Network phase profiles (solid) and phase difference profile (black). We used eqs. (52)-(53) with the following parameter values: *g*_*L*_ = 0.2, *g* = 1 and *τ* = 120 (RN) and *g*_*L*_ = 0.2078, *g* = 0 (PN). With this choice, the impedances for the two nodes have the same peak value. For the RN, *f*_*res*_ *∼* 15.8 (vertical dotted). The synaptic maximal conductances are *G*_*in*,21_ = *G*_*in*,12_ = 0.05. For comparison, we use a feedforward inhibitory network (*G*_*in*,12_ = 0, and therefore *f*_*nat*_ = 0; light blue, light red and gray). **A**. Cell 1 is the PN and cell 2 is the RN. **B**. Cell 1 is the RN and cell 2 is the PN.

**Figure S7:**
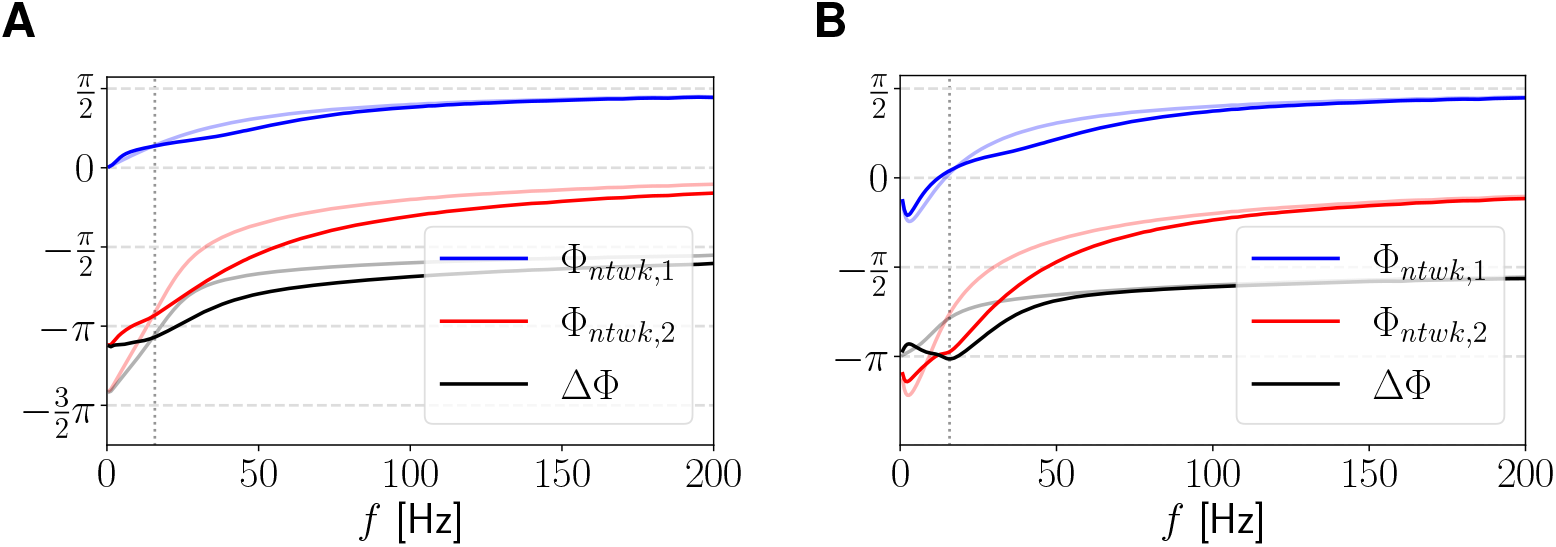
Inhibitory-excitatory network: Resonant node connected with passive node. Phases. We consider equations (52)-(53) with the following parameter values: resonant node (RN): *g*_*L*_ = 0.2, *g* = 1 and *τ* = 120, the resonance frequency for the isolate cell is *f*_*res*_ *∼* 15.8 (vertical dotted), passive node (PN): *g*_*L*_ = 0.2078, *g* = 0. **A**. Cell 1 is the PN and cell 2 is the RN. The connection parameters are *G*_*in*,21_ = 0.05 and *G*_*in*,12_ = 0.025. The network natural frequency is *f*_*ntwk,nat*_ *∼* 28.28. **B**. Cell 1 is the RN and cell 2 is the PN. The connection parameters are *G*_*in*,21_ = 0.05 and *G*_*in*,12_ = 0.05. The network natural frequency is *f*_*ntwk,nat*_ *∼* 40.05.

### S6 Impedance amplitude and phase profiles for 2D linear systems: general calculation

In order to analytically compute the impedance amplitude and phase profiles for 2D linear systems motivated by neuronal models we use

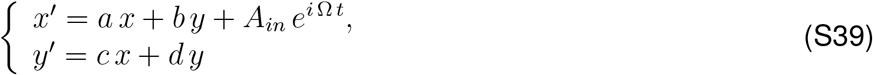

where *a, b, c* and *d* are constant, Ω *>* 0 and *A*_*in*_ *≥* 0 is the input amplitude.

#### Impedance and phase profiles

The particular solution to system (S39) has the form

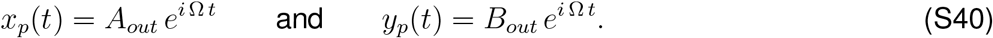

Substituting (S40) into (S39) and rearranging terms we obtain

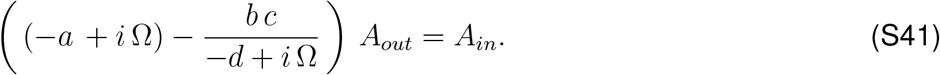

The impedance results

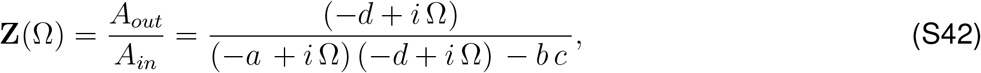

which is a complex quantity with amplitude |**Z**(Ω)| and phase Φ(*f*). For simplicity in the notation we use *Z*(Ω) for the impedance amplitude.

The impedance amplitude and phase are given, respectively, by

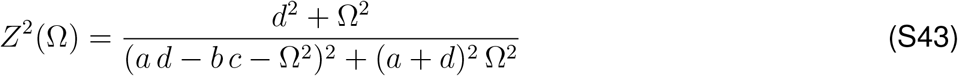

and

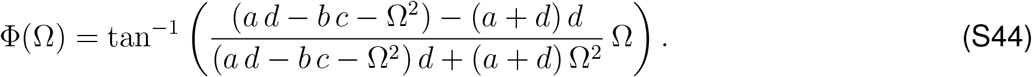

The resonance frequency, if it exists, is given by

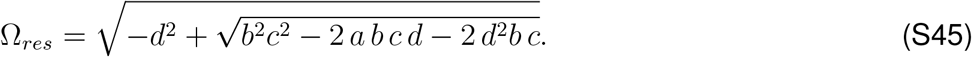

#### Impedance and phase profiles: 1D linear systems

For 1D linear systems, by making *b* = 0 in (S42) the impedance **Z** reduces to

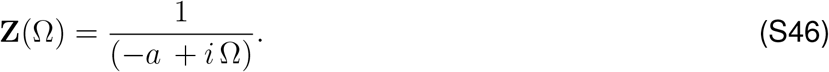

The impedance amplitude and phase are given, respectively, by

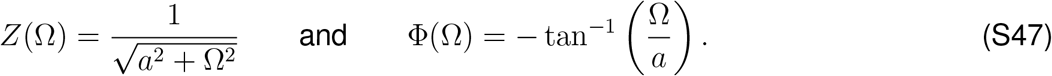

#### Characteristic polynomial for the autonomous system

The characteristic polynomial for the corresponding homogeneous 2D system (*A*_*in*_ = 0) is given by

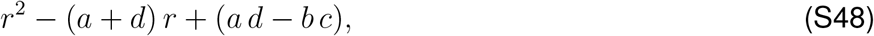

and the roots are

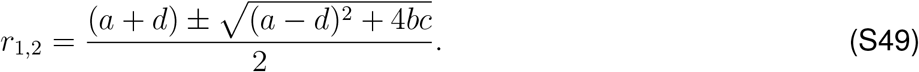

From the above equation, if 4*bc* + (*a − d*)^2^ *<* 0, the natural frequency of system (S39) is given by

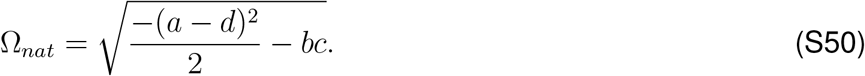

